# Hagfish genome illuminates vertebrate whole genome duplications and their evolutionary consequences

**DOI:** 10.1101/2023.04.08.536076

**Authors:** Daqi Yu, Yandong Ren, Masahiro Uesaka, Alan J. S. Beavan, Matthieu Muffato, Jieyu Shen, Yongxin Li, Iori Sato, Wenting Wan, James W. Clark, Joseph N. Keating, Emily M. Carlisle, Richard P. Dearden, Sam Giles, Emma Randle, Robert S. Sansom, Roberto Feuda, James F. Fleming, Fumiaki Sugahara, Carla Cummins, Mateus Patricio, Wasiu Akanni, Salvatore D’Aniello, Cristiano Bertolucci, Naoki Irie, Cantas Alev, Guojun Sheng, Alex de Mendoza, Ignacio Maeso, Manuel Irimia, Bastian Fromm, Kevin J. Peterson, Sabyasachi Das, Masayuki Hirano, Jonathan P. Rast, Max D. Cooper, Jordi Paps, Davide Pisani, Shigeru Kuratani, Fergal J. Martin, Wen Wang, Philip C. J. Donoghue, Yong E. Zhang, Juan Pascual-Anaya

## Abstract

Whole genome duplications (WGDs) are major events that drastically reshape genome architecture and are causally associated with organismal innovations and radiations^1^. The 2R Hypothesis suggests that two WGD events (1R and 2R) occurred during early vertebrate evolution^2, 3^. However, the veracity and timing of the 2R event relative to the divergence of gnathostomes (jawed vertebrates) and cyclostomes (jawless hagfishes and lampreys) is unresolved^4–6^ and whether these WGD events underlie vertebrate phenotypic diversification remains elusive^7^. Here we present the genome of the inshore hagfish, *Eptatretus burgeri*. Through comparative analysis with lamprey and gnathostome genomes, we reconstruct the early events in cyclostome genome evolution, leveraging insights into the ancestral vertebrate genome. Genome-wide synteny and phylogenetic analyses support a scenario in which 1R occurred in the vertebrate stem-lineage during the early Cambrian, and the 2R event occurred in the gnathostome stem-lineage in the late Cambrian after its divergence from cyclostomes. We find that the genome of stem-cyclostomes experienced two additional, independent genome duplications (herein CR1 and CR2). Functional genomic and morphospace analyses demonstrate that WGD events generally contribute to developmental evolution with similar changes in the regulatory genome of both vertebrate groups. However, appreciable morphological diversification occurred only after the 2R event, questioning the general expectation that WGDs lead to leaps of morphological complexity^7^.

---

Whole genome duplications (WGDs) are dramatic events commonly invoked causally in organismal evolution ^1^. The generally accepted ‘2R Hypothesis’^2, 3^ suggests that two rounds of WGD occurred during early vertebrate evolution (referred as 1R and 2R), however, their timing and macroevolutionary consequences remain unclear^7–9^. Most studies agree that 1R occurred before the divergence of living vertebrates, but debate centres on whether 2R predated^10, 11^ or postdated^5, 6, 12, 13^ the divergence between cyclostomes and gnathostomes (Fig. 1c). Reconstruction of the ancestral vertebrate karyotype is fundamental to unravel the timing of 2R^5, 11, 14–16^ but this goal has been stymied by a dearth of cyclostome genomes. The recently described genome of the sea lamprey (*Petromyzon marinus*) has been interpreted to support 2R occurring before^11^ or after^5^ the gnathostome-cyclostome split, or not at all (with the karyotype diversity explained as the result of large-scale segmental duplications^4, 17^). Analysis of the Arctic lamprey (*Lethenteron camtschaticum*) genome has suggested that 2R occurred in the gnathostome lineage and that two additional WGD events might have occurred in the lamprey lineage^6, 18^, perhaps shared with the hagfish lineage^6, 19^ (Fig. 1c). However, the lack of a hagfish genome assembly, the only major vertebrate group without a reference genome, has impeded assessment of the conservation of these events during early vertebrate evolution. Here we describe the outcome of sequencing and comparative analysis of the genome of the inshore hagfish, *Eptatretus burgeri* (Fig. 1a, b).

**Figure 1.**
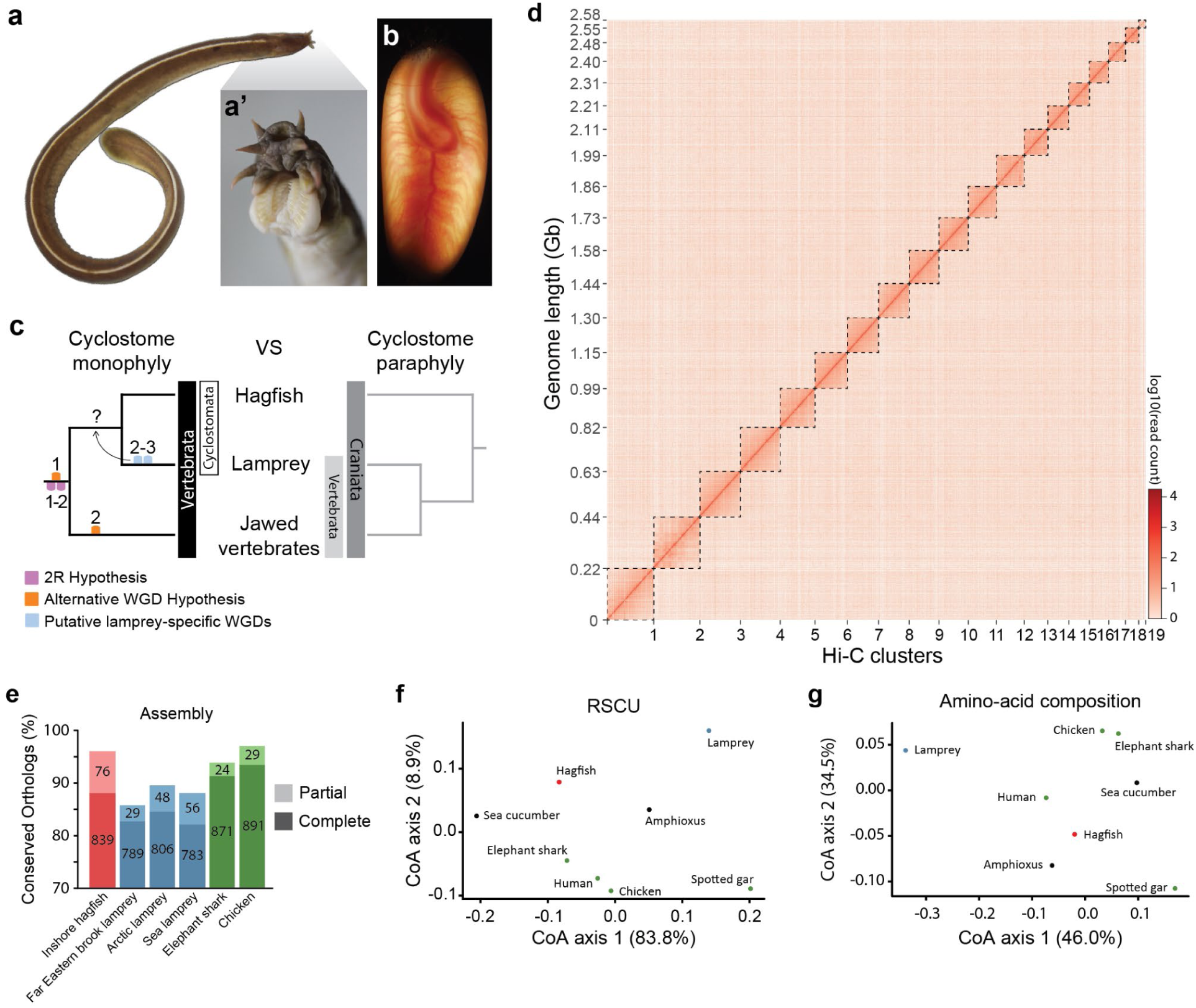
Genome of the inshore hagfish, *Eptatretus burgeri.* **a**, Dorsal view of a young adult of the inshore hagfish *Eptatretus burgeri*, with the head to the top right. The teeth apparatus (and not a jaw) can be observed in a magnification of the head region of a fixed adult individual (**a’**). **b**, Fertilised egg of *E. burgeri* with a developing embryo at stage Dean 53 (REF. 41). Blood vessels can be observed from the exterior. **c**, Two competing hypotheses of vertebrate phylogeny. WGD events corresponding to the 2R Hypothesis (lilac), to an alternative vertebrate 2R hypothesis (orange) and to those recently proposed in the lamprey lineage (light blue) are marked. Whether the lamprey-specific events did actually occur in a stem cyclostome remains elusive. **d**, Hi-C contact heatmap of the corrected lungfish genome assembly, ordered by cluster (chromosome) length. Dashed boxes indicate the cluster boundaries. **e**, Completeness assessment of the genome assembly of the inshore hagfish *E. burgeri* genome (red), three lamprey species (blue) and two jawed vertebrates (green). Number of conserved metazoan orthologs (metazoa_odb10 dataset, containing 954 BUSCOs) are indicated for each case. **f,** Correspondence analysis (CoA) on relative synonymous codon usage (RSCU) values was performed using the nucleotide sequences of all predicted genes concatenated for individual species. The percentages of variance are indicated for each axis. **g**, CoA of amino-acid composition, with the percentage of variance indicated for each axis. In (**f**, **g**): red, hagfish; blue, lamprey; green, jawed vertebrates; black, invertebrates.

## Chromosome-scale assembly and genome annotation

Like in the lamprey^20^, the hagfish genome undergoes somatic programmed DNA rearrangement, in the way of chromosome elimination^21^, making it crucial to obtain a reference assembly from a germline source. We sequenced DNA extracted from the testis of a single, sexually mature male of *E. burgeri* and generated a preliminary draft assembly using ∼240X of short-read Illumina data assisted by a Chicago *in vitro* proximity ligation assay at Dovetail Genomics^22^ (Supplementary Information). We estimated the genome of *E. burgeri* at 3.12 Gb based on k-mer frequency distribution (Extended Data Fig. 1a and Supplementary Information), in line with other hagfish species (∼2.2-4.5 Gb)^23^. Chromosome conformation capture (Hi-C) data, obtained from the testis DNA of a second individual, were used to further scaffold the genome into a final assembly (version 4.0), containing 19 contact clusters –which we consider as chromosomes for subsequent analyses— (Fig. 1d), and 9,295 unplaced scaffolds and contigs (Methods and Supplementary Information). The genome was annotated following the Ensembl annotation pipeline^24^, assisted by RNA-seq from 9 different adult tissues, and previous embryonic and juvenile transcriptomics data^19^ (Supplementary Table 11; see Methods). We generated a final gene dataset of 16,513 protein coding genes (with 27,960 transcripts), 446 long intergenic non-coding (linc)RNAs and a minority of other classes of non-coding RNA genes (Extended Data Fig. 1b). 180 microRNA (miRNA) genes were found in the *E. burgeri* genome conserved with the hagfish *Myxine glutinosa*^25^ belonging to 77 miRNA families and catalogued at MirGeneDB.org^26^. Note that although the haploid number of *E. burgeri* is 26, somatic tissues lose 8 pairs of microchromosomes during development by somatic chromosome elimination^21^ (somatic n=18). Cluster 19 and unplaced contigs/scaffolds likely correspond to these difficult-to-assemble microchromosomes, which presumably consist mainly of highly repetitive sequences and contain almost no protein-coding genes^21^ (Supplementary Information). Consistently, 98.3% (16,240/16,513) of annotated genes are located in Clusters 1-18.

BUSCO analyses show high levels of completeness of the hagfish genome (96.0 and 94.2% of single orthologs are present in the assembly and annotation, respectively; Fig. 1e, Extended Data Fig. 1c). GC-content distribution pattern analysis of the hagfish and other chordate genomes shows that the *E. burgeri* genome represents an intermediate condition between the lamprey and other chordates (Extended Data Fig. 1d), although having an overall content similar to that of the lamprey (46.7% and 48.1%, respectively). While lamprey protein-coding gene sequences have been demonstrated to pose difficult challenges for comparative analyses due to their high GC-content ^27^ (64.0%), the lower content in hagfish coding sequences (50.0%) is within the typical range of most gnathostomes and non-vertebrate chordates (42.5%-53.4%; Extended Data Fig. 1e, f, Supplementary Information). Lamprey represents an outlier in terms of both codon usage bias and amino acid composition, while the hagfish is more similar to other vertebrates (Fig. 1f, g). Hagfish genome contains, on average, significantly longer introns and intergenic regions than other vertebrates (*P* < 2.2 × 10−16, two-sided Wilcoxon sum-rank test), while the average length of coding sequences is similar to other chordates (Extended Data Fig. 1g, Supplementary Fig. 4, Supplementary Tables 18-22). This might explain the larger size of hagfish genomes and than those of lampreys^23^. Altogether, the hagfish genome may represent a better cyclostome model for relevant analyses including gene tree reconstruction and comparative genomic analyses.

## Phylogenetic position of the hagfish and gene family evolution

Whether the hagfish forms a clade with lampreys (Cyclostomata) or represents the sister to all other vertebrates (including lampreys) has depended on whether molecular or morphological evidence were considered (Fig. 1c). Morphological studies historically supported cyclostome paraphyly but more recent analyses have recovered cyclostome monophyly (reviewed in REF. 28). Phylogenies inferred from molecular evidence have almost exclusively recovered cyclostome monophyly (reviewed in REF. 25). We used Bayesian inference to reconstruct the phylogeny of vertebrates (Fig. 2, Extended Data Fig. 2a), strongly supporting a monophyletic Cyclostomata. We calculated the likelihood of gene duplication and loss patterns under the competing phylogenetic hypotheses^29^ (see Methods), finding that patterns of gene gains and losses better fit cyclostome monophyly. To further compare the two alternative hypotheses of hagfish relationships, an approximately unbiased test^30^ was performed which strongly rejected cyclostome paraphyly (log likelihood difference = 7947.7, AU = 0.004, multiscale bootstrap probability < 0.001). These results corroborate previous molecular analyses and recent morphological studies^28^ supporting the view that cyclostomes are monophyletic.

**Figure 2.**
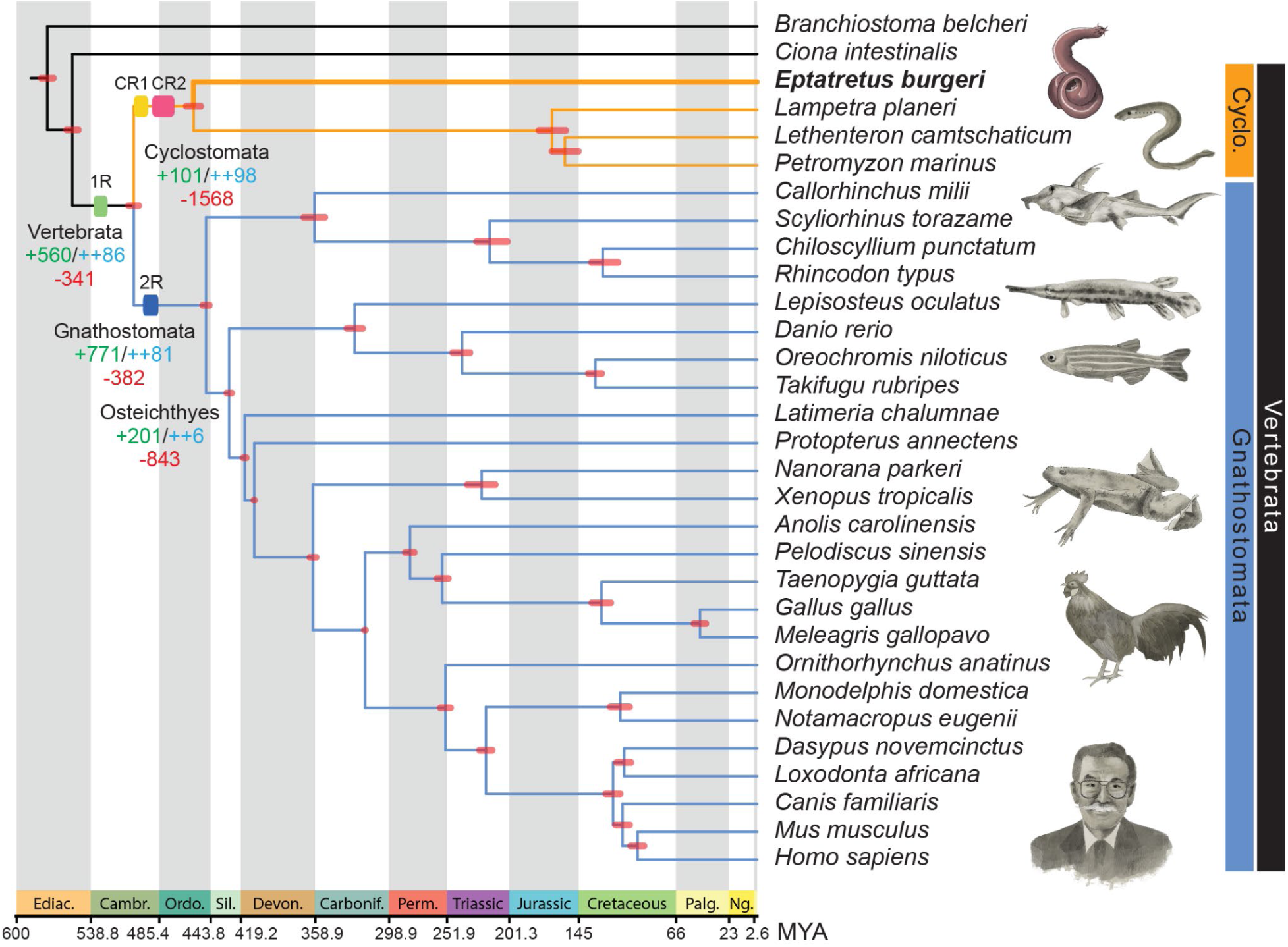
Calibrated and dated vertebrate evolution. Time calibrated, rooted phylogeny of vertebrates and two non-vertebrate species with 95% credibility intervals for clade divergence times indicated by red bars on nodes. Phylogenetic tree was obtained with Bayesian inference (Extended Data Fig. 2a), and all nodes were recovered with a posterior probability of 1. Numbers of gene family gains (green, novel HG; blue, novel core HG) and losses (red) are indicated in selected nodes (see text). Dated WGD events, including 1R, 2R and cyclostome-specific events (CR1 and CR2) described in this study, are indicated with coloured rectangles. The hagfish position is highlighted with a thicken line and bold font. Geological periods are color-coded at the bottom: Ediac., Ediacaran; Cambr., Cambrian; Ordo., Ordovician; Sil., Silurian; Devon., Devonian; Carbonif., Carboniferous; Perm., Permian; Palg., Paleogene; Ng., Neogene.

To better understand the genomic changes accompanying major transitions in chordate evolution, we used a phylogenetic-aware comparative genomic approach^31–33^ to infer ancestral gene complements and gene family gains and losses across the vertebrate tree (Fig. 2; Extended Data Fig. 3; Supplementary Information). We observed two peaks of gene novelty in both the vertebrate and gnathostome stem-lineages (novel genes: +560 and +771, respectively), also characterised by the lowest amount of gene losses (−341 and −382, respectively) (Fig. 2; Extended Data Fig. 3a; Supplementary Information). Furthermore, the fraction of highly retained novel gene families (i.e, that are not lost in descendant lineages or novel core) is the highest in the LCAs of vertebrates, gnathostomes and cyclostomes (novel core genes: ++81, ++86 and ++98, respectively; Fig. 2; Extended Data Fig. 3a). These are notably larger than those observed in other major evolutionary episodes in metazoan evolution^31, 33^, suggesting that new gene families played important roles in the origin and diversification of early vertebrates. GO enrichment analyses demonstrate that the origin of vertebrates was characterised by the appearance of genes involved in signalling pathways, cell communication and transcriptional regulation, while novel core genes involved in immunity played an important role in the origin of gnathostomes (Supplementary Information). Consistently, gnathostomes and cyclostomes have evolved independent adaptive immune systems, based on immunoglobulins in the former, and in variable lymphocyte receptors (VLR) in cyclostomes^34^ (Supplementary Information). The largest fraction of gene losses occurs in the ancestral cyclostome lineage (Fig. 2), suggesting that a strong asymmetric reduction of gene complements accompanied the early evolution of the group. The hagfish genome lacks for instance several vision and circadian rhythm-related genes, probably associated with its degenerated eyes (Supplementary Information). Inferred rates of gene duplication across Metazoa identify widespread duplications associated with the vertebrate and teleost stem-lineages (Extended Data Fig. 2b, c), probably reflecting the 1R, 2R and 3R WGD events^35^. We also inferred high duplication rates in each of the lineages leading towards crown gnathostomes and lampreys (Extended Data Fig. 2b, c) which might suggest large-scale duplications associated with these groups, consistent with the WGD events proposed recently^5, 18^. This type of analysis, however, cannot discriminate between WGD and other large-scale gene duplication mechanisms.

## Conserved *Hox* cluster evolution in cyclostomes

The number of Hox clusters and ancestral WGD events are usually correlated and, thus, the former has been used as a genomic marker of the latter. The presence of six Hox clusters in lamprey genomes^17, 18^ has been interpreted to indicate the possibility that more than two WGD events occurred in this lineage^18^. We have here extended our previous observations^19^ and confirmed the presence of 40 Hox genes arranged in 6 complete Hox clusters in *E. burgeri* (Fig. 3). Two of the hagfish clusters are located in the same chromosome (cluster 3), separated by >80Mb, likely the result of chromosomal shuffling due to the intense reorganisation of the hagfish genome from ancestral chromosomes (see below). Phylogenetic analyses of Hox coding sequences have long proven inconclusive to determine the orthology relationship between lamprey and hagfish Hox counterparts^19^. We thus applied a microsynteny conservation approach using extended Hox loci, which, together with phylogenetic analyses of selected non-Hox syntenic genes, allowed us to establish clear one-to-one orthologous correspondences between hagfish and lamprey Hox clusters, named α to ζ after the lamprey clusters^18^ (Fig. 3; Extended Data Fig. 4a-d; Supplementary Fig. 8; Supplementary Tables 29, 30). This suggests that the crown-cyclostome already possessed six Hox clusters, distinct from the crown-gnathostome ancestor, which possessed four clusters (Supplementary Fig. 9), and thus lampreys and hagfish share a genome history exclusive from that of gnathostomes. This implies that there were likely more than two WGDs in early vertebrate evolution^6, 18^.

**Figure 3.**
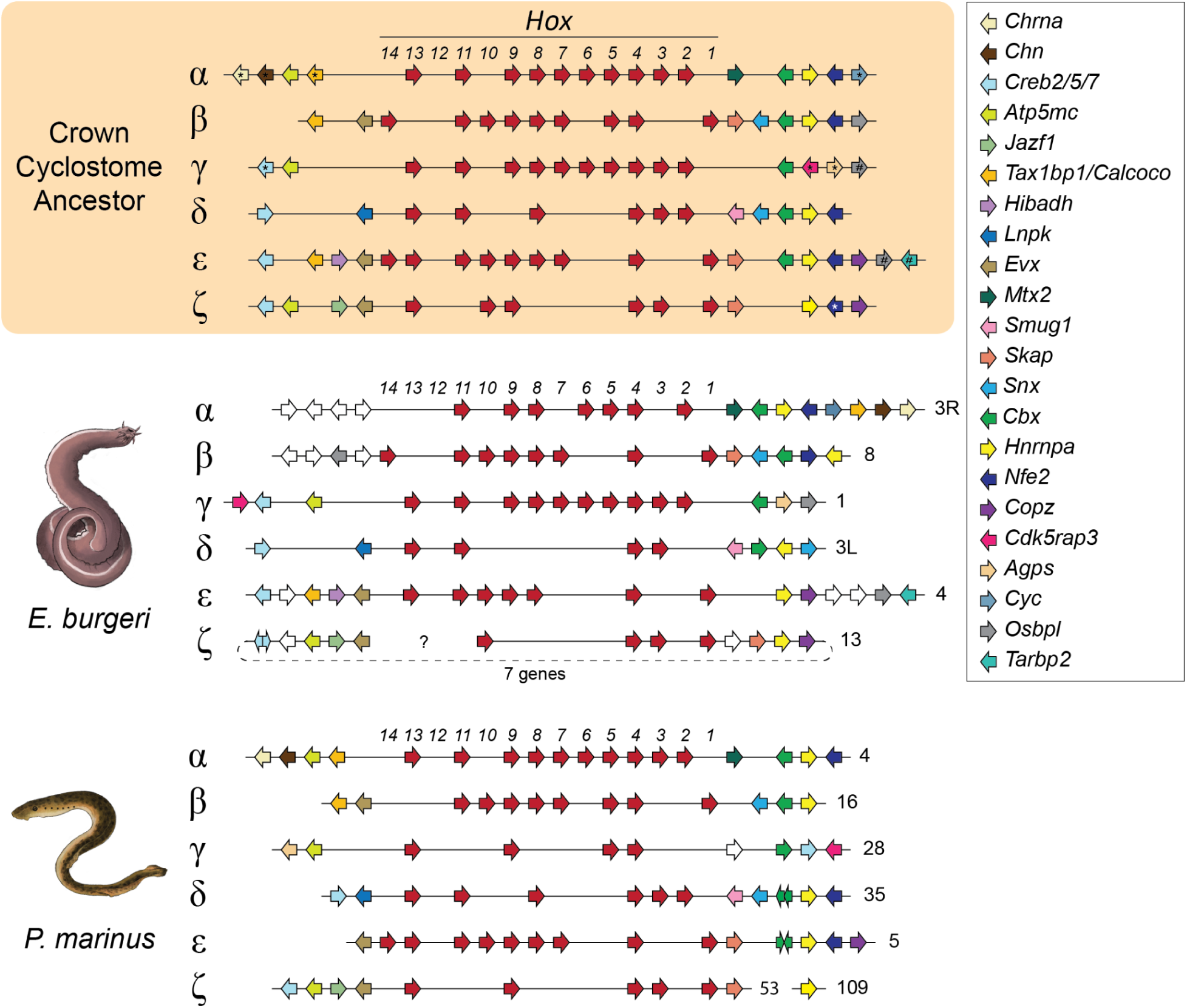
Reconstruction of the Hox complement of an ancestral cyclostome. Schematic representations of Hox clusters and syntenic genes of the inshore hagfish (*E. burgeri*; middle), the sea lamprey (*P. marinus*; bottom), and a reconstruction of the complement of the last common ancestor of hagfishes and lampreys (top). Genes are represented by coloured-coded arrows, whose direction marks the sense of transcription: Hox genes in red, non-Hox genes coloured by homology (legend to the right). The block between *Evx-ζ* to *Creb2/5/7-ζ* is assembled downstream to the *Hox-ζ* cluster, separated by 7 genes. This might be a missassembly, and their “natural” upstream position is marked by a dashed line. The sea lamprey genome scaffolds and hagfish Hi-C clusters in which Hox clusters are located are indicated to the right of each cluster, with cluster 3 separated into 3L (0-107.78 Mb) and 3R (107.78-194 Mb). Black asterisks mark genes placed at opposite sides of a cluster in the hagfish and the lamprey, but placed at one side of the ancestrally reconstructed cluster based on comparisons with gnathostomes (Supplementary Fig. 9); hashes denote genes present in lampreys in the same chromosome but at a long distance (see Supplementary Fig. 8); white asterisk: *Nfe2-ζ* is inferred due to its presence in the Arctic lamprey.

## Gnathostomes and cyclostomes share 1R but not 2R

The reconstruction of the pre-duplicative vertebrate protokaryotype by means of macrosynteny analysis is the most robust approach to test the 2R event and its phylogenetic position^36^. Previous reconstructions of the ancestral vertebrate karyotype differ greatly, with either 10-13 (REFs. 4,15) or 17 (REFs. 5,11,14) ancestral, pre-duplicative chromosomes, and also revealed a conflicting picture, in which lampreys diverged from gnathostomes before or after 2R^5, 6, 11, 18^. The lack of a hagfish genome assembly has precluded confirmation or rejection of the predictions based on lamprey genomes^37^. To shed new light on vertebrate genome evolution we performed chromosome-scale analysis of macrosynteny conservation between gnathostomes, cyclostomes and selected invertebrate deuterostomes. We first examined how 2R shaped chromosomal evolution of gnathostomes by comparing representative gnathostomes (chicken, spotted gar and elephant shark) with the genome of the sea cucumber *Apostichopus japonicus* (echinoderm)^38^ as a pre-duplicative outgroup species (Supplementary Methods). We inferred a proto-gnathostome karyotype of 17 ancestral chromosomes (ACs), largely consistent with previous studies^5, 11, 14^, but with minor differences (Supplementary Table 34, Supplementary Information). All ACs correspond to sets of four (11/17) or three (6/17) paralogous chromosomes in chicken or gar, confirming the 2R event^39^. Four of the 17 ACs correspond each to 2 or 3 linkage groups (putative chromosomes) of the sea cucumber genome (Supplementary Figure 15). Further comparison against amphioxus genome^5^ suggests that these ACs derive from pre-1R chromosome fusions in vertebrates after their split with cephalochordates (Supplementary Information). Similarly, we detected 8 fusions occurring between 1R and 2R (Extended Data Fig. 5b; Supplementary Information). We then reconstructed the gene contents of 16 of the 17 ACs (corresponding to 5,045 Belcher’s lancelet genes). With these in hand, we stringently selected 696 sets of orthologous genes (Methods and Supplementary Information), between the sea cucumber, elephant shark, chicken, gar and an AC (amphioxus) gene, and built robust chromosome-level phylogenies with a median of 38 concatenated gene sets across each of the ACs (Supplementary Table 36). The highly supported, clear-cut topologies depict the exact evolutionary trajectory from each AC to their modern chicken and gar descendants (Extended Data Fig. 5a), further supporting the existence of 2R in gnathostomes.

We next tested hypotheses of WGD timing relative to cyclostome divergence. We assessed the phylogenetic signal of hagfish and lamprey genes anchored to 656 orthologous gene sets, including elephant shark orthologs as a control for the 2R signal, and amphioxus genes as outgroups. 73.2%, 79.1% and ∼75.7% of trees including hagfish, lamprey or both hagfish and lamprey orthologous genes, respectively, are compatible with 1R (Fig. 4a). However, while 99.5% of elephant shark gene tree topologies are 2R-compatible, only 19.1% of hagfish, 10.6% of lamprey and 8.2% of cyclostome genes are compatible with a 2R history (Fig. 4a; Supplementary Files 5-8). Thus, we only find strong support for 1R as shared among cyclostomes and gnathostomes.

**Figure 4.**
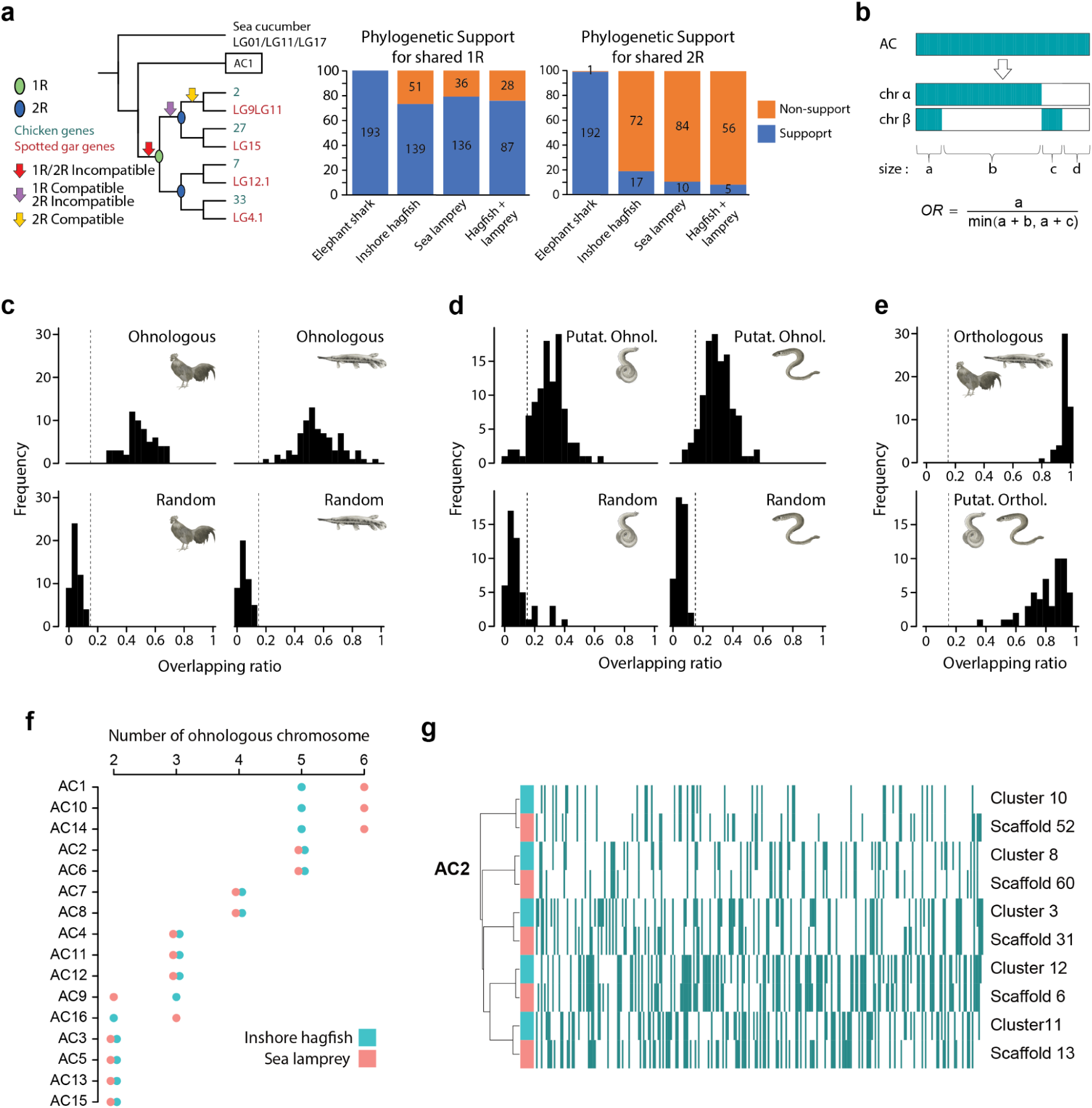
Hagfish and lamprey shared three rounds of WGDs. **a,** Phylogenetic support of gnathostome and cyclostome genes for 1R and 2R. Elephant shark, hagfish, lamprey or both cyclostomes genes (test genes) were analysed in the context of spotted gar and chicken gene phylogenies by AC (using amphioxus genes) and orthologous sea cucumber genes (outgroup). Left panel shows possible positions where test genes can group, supporting or not 1R or/and 2R (see legend). Middle and right panels show statistics of supporting (blue) or not supporting (orange) gene phylogenies from each species’ tested genes. All phylogenetic trees are available in Supplementary Files 5-8. **b**, Formula to calculate the overlapping ratio (OR) between two chromosomes. Dark cyan denotes genes from the AC, retained in modern chromosomes; white indicates gene loss. **c**, *OR* values distribution between WGD-generated paralogous (ohnologous) chromosomes in chicken (top left) and spotted gar (top right), and the artificially split chromosomes in chicken (bottom left) and spotted gar (bottom right). Dashed lines mark *OR* = 0.15. **d**, *OR* values distribution between putative ohnologous chromosomes in hagfish (top left) and lamprey (top right), and the artificially split chromosomes in hagfish (bottom left) and lamprey (bottom right). **e**, *OR* values distribution between chicken and spotted gar (top) and between hagfish and lamprey (bottom) orthologous chromosomes. **f**, Numbers of mutually ohnologous chromosomes in cyclostome genomes that correspond to each one of the 16 reconstructed ACs. **g**, Retention profile clustering analysis of cyclostome chromosomes deriving from AC2. Retained genes are denoted by dark cyan lines. Five putative orthologous chromosome pairs are defined.

To further confirm the timing of the 1R event, we investigated whether signals of the four pre-1R and eight post-1R fusion events are present in cyclostomes. When assessing how hagfish and lamprey chromosomes descended from the 17ACs, we found that the hagfish genome displays a large amount of rearrangement –at least 52 fusions detected—, making any signal of hypothetical shared events unreliable (Extended Data Fig. 6b). However, most lamprey chromosomes are descendants of single ACs^5^ (Extended Data Fig. 6a), making the lamprey a better model to investigate these rare genomic changes. We found that the sea lamprey genome bears signals of three of the four pre-1R fusions but none of the eight post-1R fusions (Supplementary Information; Supplementary Fig. 24), suggesting that the lamprey (and, thus, cyclostomes) diverged after the 1R but before all eight post-1R/pre-2R fusions detected in gnathostomes. The fourth pre-1R fusion corresponds to AC3, which corresponds to two separate chordate linkage groups (CLGs) as described in the model of Simakov et al.^5^ (termed CLGI and CLGQ in Simakov’s nomenclature), implying that independent pairwise fusions could have occurred after the 1R in gnathostomes and mimicking a single pre-1R fusion event (Supplementary Fig. 23). *In silico* simulations show that this would not be rare (30% of cases expected by chance; Supplementary Table 38; Methods). Thus, the number of ancestral chordate chromosomes could be 18, as recently noted^39^. Taken together, our comprehensive phylogenetic analysis and the constraints given by pre- and post-1R chromosomal fusions provide strong evidence in favour of a pan-vertebrate 1R event, but constraints 2R to the gnathostome lineage.

## Cyclostome-specific whole genome duplications

It has been suggested that the lamprey genome has been shaped by at least three WGD events^6, 18^ and the presence of six orthologous *Hox* clusters in lamprey and *E. burgeri* (Fig. 3) would imply that this is the ancestral condition for cyclostomes^6, 19^. Although we find that multiple chromosomes are direct descendants of each AC in both cyclostome groups, the extensive rearrangements observed in the hagfish and the large haploid number in the lamprey impedes chromosome-level macrosynteny conservation analysis in order to distinguish intraspecific ohnologous and interspecific orthologous relationships. To confidently infer karyotype evolution in cyclostomes we developed a new metric, the ‘overlapping ratio’ (*OR*), to measure the similarity of gene retention profiles of any two chromosomes hypothetically descending from a common AC (Fig. 4b, Supplementary Information). A retention profile is defined by a vector listing the presence or absence of genes on a modern vertebrate chromosome from their corresponding AC. Therefore, we expect the *OR* of chromosomes deriving from a duplication event to be significantly higher than that of chromosomes deriving from an ancestral fission followed by gene translocations. As proof of concept, we applied this metric to gnathostomes: knowing their genomes have been shaped by the 2R event, we found that the median *OR* of ohnologous chromosome pairs in chicken or spotted gar was 0.49 (interquartile range, IQR: 0.44-0.56) and 0.54 (IQR: 0.47-0.65), respectively (Fig. 4c), while *OR* values for simulated fission-derived chromosome pairs was never larger than 0.15, indicating that ohnologous chromosomes indeed share more retained genes (Fig. 4c, Methods).

We then applied the *OR* metric to the sea lamprey (assisted by a meiotic map of the Pacific lamprey *Entosphenus tridentatus*^17^; Supplementary Table 44) and hagfish, defining ohnologous chromosome pairs as those with *OR* > 0.15. We found that the median *OR* between putative ohnologous chromosomes was 0.30 (IQR: 0.23-0.36) and 0.29 (IQR: 0.23-0.37) for the lamprey and hagfish, respectively (Fig. 4d; Extended Data Fig. 7a,b; Supplementary File 9). Using this approach, we found that most ACs analysed (12/16 or 75%) have descended into at least three mutually ohnologous chromosomes in both the lamprey and hagfish (Fig. 4f), suggesting that, at least, a second WGD occurred in cyclostomes. In both genomes, at least five chromosomal regions are direct descendants of each of the same five ACs (1, 2, 6, 10 and 14) by duplication, with 3 ACs contributing to 6 chromosomes each in the lamprey (Fig. 4f). The distribution of multiplicity across ACs is highly correlated across the two species (Spearman ρ=0.92), suggesting that cyclostome genomes have been shaped by three shared WGD events. It is expected that *OR* will decrease with each WGD (Supplementary Information, Supplementary Fig. 25) and, thus, the lower value in cyclostomes is consistent with the occurrence of this extra WGD.

To further confirm that these proposed additional WGD events are shared by hagfish and lamprey, we extended the use of *OR* to detect putative orthologous chromosomes. During the process of diploidization after a WGD event, two descendant chromosomes diverge and fix their mutations independently and so it is expected that interspecific orthologous chromosomes will have more similar gene retention profiles than intraspecific ohnologous chromosomes. Accordingly, orthologous chromosomes of chicken and spotted gar have a median *OR*=0.96 (IQR 0.95-0.98; Fig. 4e; Supplementary Table 41) and clustering-based analysis based on gene retention profiles places chicken and gar orthologous chromosomes closer to each other, completely reflecting the phylogenetic signal (Extended Data Fig. 7c; Supplementary File 10; Supplementary Information). When we applied this approach to cyclostome genomes, we found the median *OR*=0.84 (IQR: 0.74-0.91; Fig. 4e) for 52 (∼87%) chromosome pairings between lamprey and hagfish that putatively represent 1:1 orthologs –higher than that of ohnologous chromosomes— and only 8 (∼13%) one-to-two or two-to-one ambiguous relationships, probably due to secondary independent chromosome losses in either group (Supplementary Information). Clustering analysis of retention profiles recovers orthologous relationships between lamprey and hagfish (Fig. 4g; Supplementary File 11). Overall, intra- and interspecific gene retention profile analyses indicate that two independent WGD events took place in the cyclostome stem-lineage; we refer to these as CR1 and CR2, to avoid confusion with the gnathostome-specific 2R event.

## Increase of developmental regulatory complexity

To investigate the immediate consequences of the independent CR1/CR2 events on cyclostome genome evolution, we first asked whether retained duplicates (ohnologues) are especially associated with developmental functions in the hagfish as in their gnathostome counterparts^14, 40^. Gene ontology enrichment analysis shows that hagfish gene ohnologues are also significantly enriched for functions associated with developmental processes (Extended Data Fig. 8a, b; Supplementary Information). Gnathostomes have increased their regulatory complexity (higher number of regulatory regions per gene), particularly of developmental ohnologues ^40^. We identified accessible chromatin regions (ACRs) as putative non-coding regulatory elements in the hagfish genome with an assay for transposase-accessible chromatin coupled to sequencing (ATAC-seq), using two embryos of *E. burgeri* at different stages^41^ (45 and 53; Supplementary Fig. 28). We found a significantly higher number of ACRs per gene (Fig. 5a) (similar to gnathostomes^40^ but unlike the cephalochordate amphioxus), particularly in distal regions from transcriptional start sites (Fig. 5b,c; Extended Data Fig. 8c). This pattern is especially evident in developmental genes (Fig. 5d; Extended Data Fig. 8d-f), implying that their higher retention after cyclostome WGD events is underlain by a more complex regulatory landscape of developmental genes, as in gnathostomes^40^.

**Figure 5.**
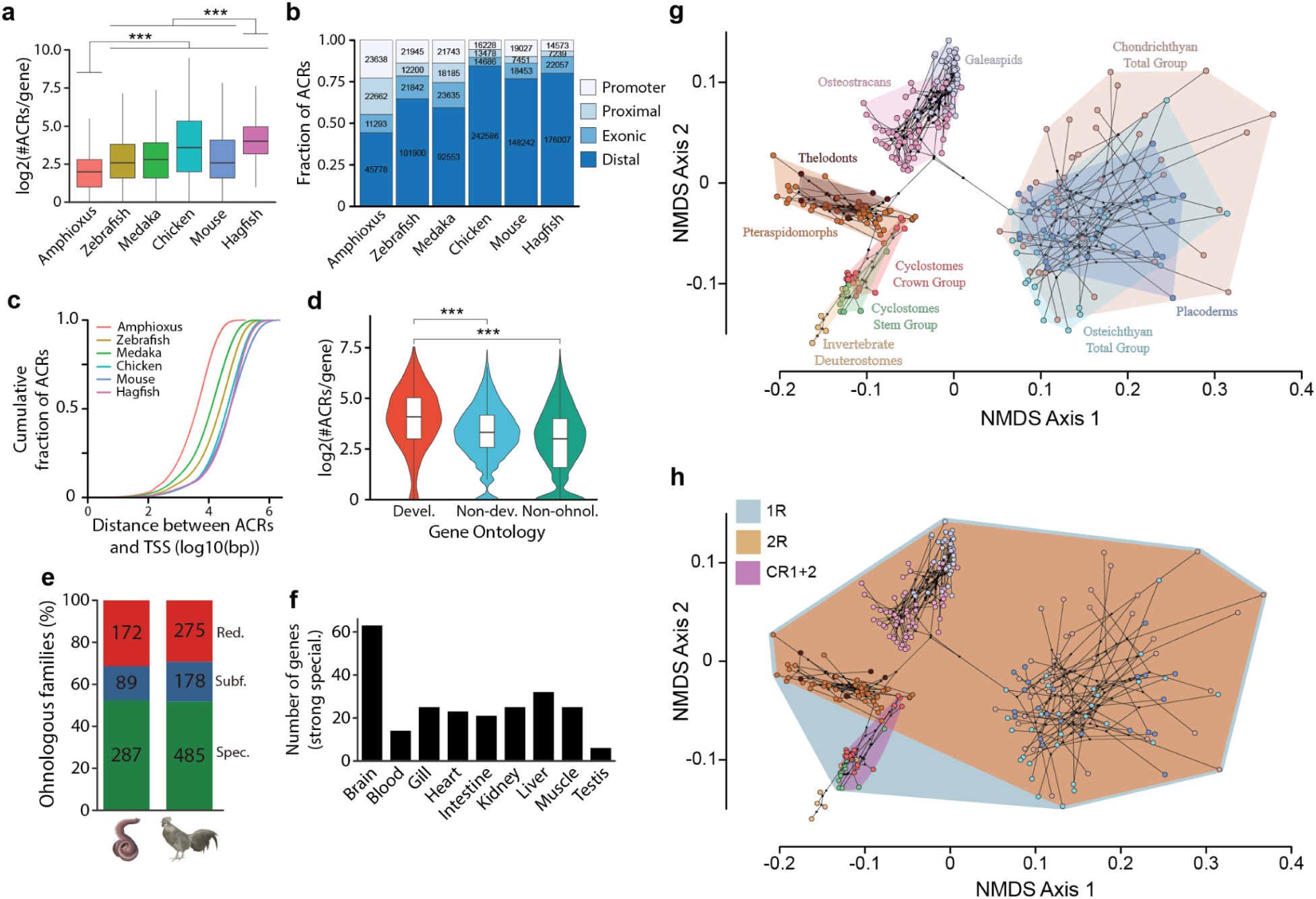
Impact of WGD events on the regulatory genome and morphological evolution of vertebrates. **a**, Distributions of the ACR numbers within the *cis-*regulatory regions of each gene (see Methods). **b**, Numbers and fractions of ACRs with respect to genomic annotations in each species. Promoters, between 1 kb upstream and 0.5 kb downstream of annotated TSSs; proximal, within 5 kb upstream and 1 kbp downstream of annotated TSSs, but not overlapping promoters; exonic, within exons of protein-coding genes but not overlapping proximal regions; distal, not in aforementioned locations. **c**, Cumulative proportion of the distance of ACRs from the closest TSSs in each species. Statistical information for panels **a**, **b**, and **c** are given in Supplementary Tables 46–50. **d**, The distribution of ACR number across different classes of genes, according to PANTHER Gene Ontology database (devel, developmental ohnologs; non-dev, non-developmental ohnologs; non-ohnolg, singletons). **e**, Distribution of fates of ohonologous families after WGD. **f**, Number of ohnologs with strong specialization expressed in hagfish tissues **g**, **h**, Morphological disparity across vertebrates. Non-metric ordinations are presented highlighting the morphological variance among (**g**) taxonomic lineages of extant and extinct vertebrates and (**h**) the descendants of 3 whole genome duplication events. Convex hulls have been fitted around groups. The underlying tree is derived from a consensus of relationships from the literature. In panels a and d, ***, *P* values < 2.2 × 10−16, two-sided Wilcoxon sum-rank tests.

In gnathostomes, retained duplicates can evolve via expressional specialization (reduction of expression domains of one of the ohnologs)^40^, likely coupled to neofunctionalization rather than subfunctionalization (differential erosion of enhancers)^42^. Taking advantage of adult transcriptome data across nine organs (see Methods), we next analysed the fate of hagfish ohnologues after CR1/CR2. Hagfish duplicates also tend to reduce their expressional domains: over 68% and 71% of gene families subfunctionalized or specialized in the hagfish and chicken, respectively (Fig. 5e). Hagfish ohnologues that have restricted their expression domains (subfunctionalization or specialization) are associated with a larger amount of regulatory elements and a higher sequence evolutionary rate than those that have maintained the ancestral patterns (Extended Data Fig. 8g,h), similar to gnathostomes (Extended Data Fig. 8i)^40^. Furthermore, the largest portion of ohnologues with strong specialization (one or two ancestral expression domains) are expressed in the brain (Fig. 5f), mirroring the pattern observed in gnathostomes^40^ (Extended Data Fig. 8j). In summary, our results indicate that cyclostomes and gnathostomes followed parallel evolutionary pathways after their independent WGD events. Genes gained a larger regulatory complexity, mostly on distal regions and especially in duplicates with developmental functions, which tend to be retained more often. Furthermore, specialization is a common fate of ohnologs associated with faster sequence evolution and the acquisition of novel regulatory elements that drive their tissue-specific expression. Alternatively, the possibility that a decrease in the number of regulatory elements took place in the amphioxus cannot be confidently ruled out.

## Impact of WGD events on vertebrate morphological diversity

Hypotheses on the role of WGD events in the origin and elaboration of the vertebrate body plan range from deterministic to permissive^7^. There can be no doubt that many vertebrate and gnathostome novelties are contingent on gene paralogues that are the product of the 1R and 2R events, though whether WGD played a causal role remains unclear. We employed two tests of a causal relationship: (i) absolute timing of the WGD events and the clades with which they are causally associated, and (ii) contrast in phenotypic diversity before and after the WGD events. Using a dataset of 177 genes and 33 fossil calibrations, our relaxed molecular clock analyses estimate the 1R event to have occurred 536-525 Ma (early Cambrian), 15-34 Myr before the divergence of crown-vertebrates (510-502 Ma; middle Cambrian) (Fig. 2); the CR1 and CR2 events are dated in a rapid succession to 505-494 (late Cambrian) and 491-473 Ma (latest Cambrian - earliest Ordovician), respectively, at least 10-39 Myr prior to the 463-452 Ma (Middle-Late Ordovician) divergence of crown-cyclostomes (Fig. 2); and the 2R event is dated to 498-486 Ma (late Cambrian), 46-53 Myr before the divergence of crown-gnathostomes (450-445 Ma; late Ordovician^43^) (Fig. 2). To characterise phenotypic disparity across WGD events we compiled a phenotype matrix composed of 577 traits for 278 living and fossil chordates, encompassing all aspects of morphology, which we ordinated using Non-Metric Multidimensional Scaling (Fig 5g). This shows that each duplication is followed by an increase in phenotypic disparity through occupation of novel regions of morphospace (Fig 5h), but the majority of chordate disparity (88-97% of the morphospace encompassed by a vertebrate convex hull) emerged subsequent to the 2R event (Fig. 5h). Thus, while all three of WGD events are of comparable antiquity, there is a stark contrast in terms of phenotypic evolution and species diversity between the descendents of 2R and the other WGD events. 2R occurred very early within the gnathostome stem-lineage, before the acquisition of gnathostome phenotypic characters. This is evidenced by the ‘ostracoderm’ fossil record^7, 44^ which shows that the accrual of phenotypic disparity following the 2R event was neither delayed nor explosive. Rather, phenotypic characters were acquired along the gnathostome stem during the 46-53 Myr between 2R and the divergence of crown-gnathostomes (cf. REF. 7).

## Discussion

The hagfish genome offers unique insights into the biology of early vertebrates. First, it reveals a robust and accurate history of WGD events in early vertebrates, demonstrating that cyclostomes diverged from gnathostomes after the 1R, but before the 2R. This is consistent with early studies on the matter^12, 14^ ending debate over the timing of 2R^37^ and evidencing two additional WGD events, CR1 and CR2, in stem-cyclostomes. Thus, key vertebrate innovations (e.g., elaborate tripartite brain, neural crest cell-derived tissues among other novelties^45^) originated in a tetraploid stem-vertebrate, before 2R. This basic vertebrate body plan was elaborated independently in cyclostomes and gnathostomes as a result of their lineage-specific genome duplications, for instance, facilitating the evolution of different adaptive immune systems (immunoglobulin-based in jawed vertebrates, VLR-based in cyclostomes^34^), or the appearance of key morphological innovations, such as the jaw and paired appendages in gnathostomes. Interestingly, these independent WGD events shaped their ancestral genomes in similar ways, by permitting an increase in regulatory complexity, especially of genes with roles in development. Duplicates of developmental genes are indeed more likely to be retained in both lineages, highlighting the crucial role of development in evolution of novel complex traits.

The contrasting phenotypic consequences of 2R versus the other WGD events might suggest that there is no direct causal relationship or that there should be no general expectation of macroevolutionary consequences from WGD events, despite their clear impact increasing regulatory potential of the genome. Another possibility is that the 2R event is different in nature from the 1R and CR events. Indeed, a number of recent studies have suggested that while 1R was likely an autopolyploidy event, 2R was an allopolyploidy^5, 6, 46^. This is significant since the macroevolutionary consequences of allopolyploidy are expected to be more immediate than autopolyploidy, resulting in chromosomal rearrangements, changes in chromatin structure, DNA methylation, gene expression and the activation of transposable elements^47–49^, extensive and immediate changes that promote species and ecological diversification^50, 51^ as well as evolutionary novelty^52–55^. This may go some way to explain why the evolutionary consequences of the 2R WGD are so much greater, leading to the profound diversification of gnathostome bodyplans that have dominated vertebrate communities since the early Palaeozoic.

### Online content

Any methods, additional references, Nature Portfolio reporting summaries, source data, extended data, supplementary information, acknowledgements, peer review information; details of author contributions and competing interests; and statements of data and code availability are available at https://doi.org/XX.XXXX/XXXXXXXXX.

## Supporting information

Supplementary Information, including Supplementary Figures 1-31

## METHODS

No statistical methods were used to predetermine sample size. The experiments were not randomized, and investigators were not blinded to allocation during experiments and outcome assessment.

### Animal sampling and experimentation

Adult inshore hagfish animals were captured off the coast of Shimane, Japan, as previously described^56^. Hagfish embryos (staged according to Oisi et al.^41^) used for ATAC-seq were obtained as previously described^19, 56^. The sampling and experiments were conducted according to the institutional and national guidelines for animal ethics, approved by the RIKEN Animal Experiments Committee (approvals H14-25-23 and H14-25-25).

### Genome sequencing and assembly

We sequenced a mix of short-insert paired end and long-insert mate pair libraries prepared from DNA extracted from the testis of a single, sexually mature, male individual of the inshore hagfish, *E. burgeri*, resulting in ∼240X of Illumina clean data (Supplementary Tables 2 and 3 and Supplementary Information). Hagfish species have large genome sizes, ranging between ∼2.2-4.5 Gb^23^. We estimated the genome of *E. burgeri* at 3.12 Gb based on k-mer frequency distribution (Extended Data Fig. 1a and Supplementary Information), in line with other hagfish species. We assembled the genome of *E. burgeri* following gradual steps using different strategies. First, we obtained a primary assembly using just the Illumina short-read data (version 2.0). To improve contiguity, this primary assembly was supra-scaffolded using Chicago *in vitro* proximity ligation at Dovetail Genomics^22^ significantly increasing the N50 from 0.44 to 2.69 Mb. This assembly was polished with all short-insert sequencing data using Pilon^57^ v1.22 and the resulting version (3.2 in our pipeline) was made publicly available in both public repositories (GenBank accession no. GCA_900186335.2) and Ensembl genome browser^58^ (release 93; https://www.ensembl.org/Eptatretus_burgeri/). We further sequenced over 2200X of raw Hi-C short read data from a second adult, male individual and obtained approximately 350X valid Hi-C contact data to improve scaffolding. Hi-C contacts have also been used to correct 280 likely misjoined scaffolds (Supplementary Information). After a process of parameter optimization, we have used LACHESIS^59^ to assemble 1,573 scaffolds into 19 Hi-C contact clusters. Detailed assembly procedures can be found in Supplementary Information.

### RNA-sequencing

Adult tissues were dissected from two adult male individuals of *E. burgeri* (brain, gills, liver, intestine, heart, skeletal muscle, kidney and testis from animal #20150825; blood from animal #20150917). Total RNA was extracted using a RNeasy Plus Universal Mini kit (QIAGEN) for the brain, heart, skeletal muscle, kidney and testis samples, and with ISOGEN (Nippon Gene Co.), a guanidinium thiocyanate-phenol-chloroform based extraction protocol, for the intestine, liver and gill samples. In all cases, DNA was removed including a DNAseI step. RNA-seq libraries were prepared with the TruSeq Stranded RNA Lib Prep Kit (Illumina, San Diego, CA) and quantified by qPCR using the KAPA Library Quantification Kit for Illumina Libraries (KapaBiosystems, Wilmington, MA) for all samples. Library profiles were assessed with an Agilent 2100 Bioanalyzer (Agilent Technologies). All libraries were sequenced at RIKEN BDR in an Illumina HiSeq 1500 platform, 127-bp paired-end mode.

### Genome annotation

Annotation of the hagfish genome assembly version 3.2 was created via the Ensembl gene annotation system^24^, assisted by RNA-seq data from 9 adult tissues (this study) and by developmental RNA-seq data from three embryos (Dean stages 35, 40 and 45) generated in a previous study^19^. Coordinates of annotated features were later converted to the final Hi-C assembly, version 4.0. Detailed methodology and annotation results can be found in Ensembl (http://www.ensembl.org/info/genome/genebuild/2018_06_eptatretus_burgeri_genebuild.pdf). In addition, before performing phylogenetic analyses corresponding to Fig. 4a, 1957 gene models were manually corrected with the numbers ranging between 120 (amphioxus) and 703 (sea lamprey).

### GC-content, codon usage and amino acid composition

Overall GC-content percentage was analysed for whole genomes of the inshore hagfish and 9 other chordate genomes (human, *Homo sapiens*; chicken, *Gallus gallus*; *Xenopus tropicalis*, zebrafish, *D. rerio*; spotted gar, *Lepisosteus oculatus*; elephant shark, *Callorhinchus mili*; sea lamprey, *Petromyzon. marinus*; sea squirt, *Ciona robusta*; and the Floridian lancelet, *Branchiostoma floridae*) (Supplementary Table 12). GC-content distribution (Extended Data Fig. 1d) was calculated from non-overlapping sliding 10-kb windows. To calculate codon type frequency, we categorised each codon into GC-0/1/2/3 based on the number of G or C bases in a codon. The summed frequency of usage for each category is the sum of the normalized frequency of codon usage for all codons included in each category. To plot the distribution of GC content per codon position, GC percentage of each codon position for each protein coding gene (with only longest coding sequence per gene) was calculated, as well as the GC content for each whole coding sequence (equivalent to GC content of all three codon positions). Correspondence analyses on RSCU (relative synonymous codon usage) and amino acid composition were done with software codonW according to Smith et al.^60^.

### Completeness evaluation of genome and annotation

We used BUSCO^61^ v. 5.2.2 to assess the completeness of genomes at both assembly and annotation levels of hagfish (Eptatretus burgeri), three lamprey species (Far Eastern brook lamprey, *Lethenteron reissneri*; sea lamprey, *Petromyzon marinus*; and Arctic lamprey, *Lethenteron camtschaticum*) and two jawed vertebrates (elephant shark, *Callorhinchus milii*; and chicken, *Gallus gallus*). The program was run in both ‘genome’ and ‘protein’ modes, with gene predictor ‘metaeuk’ against the core metazoan database embedded in BUSCO (metazoa_odb10 dataset, built on 2021/2/17, with 954 BUSCOs).

### Species tree inference

Orthogroups of protein coding genes previously used in the analysis of the spotted gar genome^62^ were extended using HaMSTR^63^(Ebersberger, et al. 2009). The spotted gar genes from Braasch et al.^62^ were used as bait sequences in HaMSTR, which sequentially added the best matching protein sequence for each species provided the bait sequence was in turn the best match in the spotted gar proteome (reciprocity was fulfilled). HaMSTR uses HMM profiles to assign similarity scores. Of the 242 alignments used in the spotted gar study, 190 remained single copy in all the taxa used here. These were used to reconstruct the topology of vertebrates. The 190 protein families were individually aligned using MAFFT^64^ version 7.402 with default settings, concatenated to form an alignment of 310,527 sites and trimmed with automatic method selection in trimAl^65^ version 1.2 (-automated1). This concatenated alignment containing 84,017 sites is available within the Supplementary File 13. This was used to infer a phylogeny (also provided within Supplementary File 13). We used PhyloBayes^66, 67^ version 4.1 with the CAT^68^ GTR^69^ model with 4 discrete gamma categories for site rates^70^. The analysis can be repeated in PhyloBayes with: phylobayes – pb -d alignment -cat -gtr -dgam 4. Convergence was analysed visually in Tracer^71^ and using bpcomp and tracecomp - part of the PhyloBayes suite. Six chains were run for between 12,991 and 13,836 cycles. After a burnin of 1500 cycles, bpcomp revealed that all bipartitions were present in the exactly the same frequencies (maxdiff and meandiff = 0). Tracecomp revealed effective sample sizes of parameters ranging from 522 to 11,491 with relative differences of 0.018 to 0.218. We deem that, at least for topology construction, these chains have converged sufficiently, therefore, recovered topologies reflect the true posterior distribution.

### Dating species divergences

For molecular clock analysis, we expanded the dataset to include several non-vertebrate outgroups because many of the calibrations have similar maximum bounds, meaning the effective time prior would be older than intended if we did not include the outgroups. HaMSTR was then used to extend the orthogroups to include the new taxa. Of the original 190 orthogroups, 172 were retained in single copy in all taxa; they were aligned and trimmed as before. This alignment (provided within Supplementary File 14 - partitioned_alignment.phy) was used as input to MCMCtree^70^ using approximate likelihood estimation^72^. The analysis was run on each gene under the simplest possible model. The temporary control files were then used as input to CODEML for each gene with the following modifications. The substitution model was changed to the one that was preferred by ProtTest^73^ from a subset of LG^74^, WAG^75^, JTT^76^, Dayhoff^77^ and BLOSUM62^78^. fix_alpha was set to 0, alpha was set to 0.5 and the number of gamma categories was set to 5. The Hessian matrices generated were concatenated to form the in.BV file which was used for the approximate likelihood estimation in the full analysis. The time prior was constructed by applying a uniform prior distribution with a hard minimum bound and a soft maximum bound (with 2.5% probability greater than the maximum) to nodes. We used the autocorrelated rates clock model with a gamma, prior distribution with shape = 2, scale = 4.53. This was constructed by dividing a typical distance between two tips whose most recent common ancestor was at the root of the tree under LG + F + G4 (inferred with IQ-TREE^79^ v1.6.3) by the expected time for the tree based on root prior. This was multiplied by the shape parameter of 2 (leading to a fairly flat gamma distribution, corresponding to a relatively uninformative prior). The variance prior (sigma2) had shape = 1, scale = 1, meaning variation in rates are not highly penalised in the posterior distribution. Rates across sites were modelled by a gamma distribution with shape = 1 and scale = 1 with 5 discrete categories. After a burn-in period of 10,000 generations, parameter values were saved every 20^th^ generation until 20,000 cycles were saved (400,000 generations total). Convergence was investigated in Tracer^71^, revealing convergence had been reached in the six chains run (the lowest effective sample size was 194 and posterior distributions in all 6 chains looked almost identical). The alignments, control files and tree are available from the Supplementary File 14.

### Estimation of gene duplication rates

Orthogroups were predicted using OrthoFinder^80, 81^ version 2.3.5; output from this analysis is available as Supplementary File 15. OrthoFinder includes a gene duplication prediction step as part of its pipeline. Gene duplication events presented here have greater than 50% support. The species tree was fixed to the topology inferred in this study.

### Rooting the vertebrate phylogeny

Orthogroups were predicted using OrthoFinder^80^ version 2.3.5 for only the vertebrate taxa (hagfish, lampreys and gnathostomes). For each gene family, sequences were aligned using MAFFT^64^ with default settings then trimmed using trimAl^65^ with heuristic choice of trimming parameters. IQ-TREE^79^ was then used to generate 1000 bootstrapped trees in a maximum likelihood framework with the model selected using ModelFinder^82^. These bootstrapped trees were used as the input to ALEobserve to create ALE objects. Two species trees were used as hypotheses; one with hagfish sister to all other vertebrates and one with monophyletic cyclostomes. ALEml_undated^83^ was used with each of these species tree hypotheses, with default settings apart from tau (the transfer rate) set to 0, meaning transfers could not be inferred. This estimates the pattern of gene duplication and loss for each gene family under the different species tree hypotheses, as well as a likelihood under the species tree. An approximately unbiased test^30^ was then performed on the likelihoods of each gene family under the two competing hypotheses, using the program CONSUL^84^.

### Ancestral gene family complements

A total of 45 animal genomes (Supplementary Table 1; Extended Data Fig. 3) were compared using a pipeline described previously^31–33^. Briefly, the proteomes were compared using a reciprocal blastp of all-vs-all sequences with DIAMOND^85^ (e-value threshold of x10-5). Markov Cluster Algorithm (MCL^86^) was used to infer homology groups (HGs) from the BLAST output, with default inflation parameter (I=2). GOs were assigned to the different HGs by analysing the human protein sequences in each HG with PANTHER GO^87^.

### Orthology relationships of *Hox* gene clusters

Hagfish Hox sequences were obtained from a previous study and used as queries with TBLASTN to find the location in the Hi-C assembly and Ensembl annotation. Information about Hox syntenic genes in lamprey, human and elephant shark, and the European amphioxus as outgroup, were obtained from previous studies^17–19, 88^, downloaded from Ensembl or NCBI GenBank and used as queries to find their presence in the hagfish Hi-C assembly and Ensembl annotation with TBLASTN. Location of Hox and their syntenic genes, as well as their Ensembl Gene IDs is provided in Supplementary Table 29. For phylogenetic analysis of Hnrnpa, Cbx, Gbx and Agap, amino acid sequences were aligned using MUSCLE^89^ as implemented in MEGAX^90^ v.10.2.4. The alignment was trimmed by trimAl^65^ v.1.2rev59 using the ‘-automated1’ option, and then formatted into a nexus file using readAl (bundled with the trimAl package). The Bayesian inference tree was constructed using MrBayes^91^ v.3.2.6, under the assumption of an LG + I + G evolutionary model, with two independent runs and four chains. The tree was considered to have reached convergence when the standard deviation stabilised under a value of less than 0.01. A burn-in of 25% of the trees was performed to generate consensus trees. Multisequence alignments with MrBayes parameters and number of generations for each tree are provided as supplementary files in Supplementary Files 16-19.

### Dating genome duplications in vertebrates

OrthoFinder inferred gene families were selected that showed a clear signal of both the 1R and 2R duplication events and were broadly congruent with current phylogenetic hypotheses. This resulted in 28 gene families, in which each gnathostome is represented up to four times and each cyclostome twice. Gene families containing a signal of both 1R, 2R, and the cyclostome duplication events (CR1, CR2) were rare and so to date the CR events, an additional dataset was assembled by extracting OrthoFinder derived gene families that contained at least 3 genes from the same set of AC-anchored ohnologs in either hagfish or lampreys and hence showed signal for both CR1 and CR2. The resulting dataset consisted of 10 gene families, in which each cyclostome species was represented by three to four gene copies (Supplementary Information).

For each analysis, taxon sampling towards the root of the tree was improved by including additional outgroup taxa *Nematostella vectensis* (Cnidaria)*, Trichoplax adhaerens* (Placozoa)*, Mnemiopsis leidyi* (Ctenophora) and *Hofstenia miamia* (Xenacoelomorpha); this served to remove the nodes of interest from the root of the tree and include additional relative and absolute calibration information for more universal clades. Individual gene families were aligned using MUSCLE^89^ and trimmed using the ‘-automated1’ option in trimAl^65^. The best fitting model for each gene family was determined using IQ-TREE^79^ and all gene families were concatenated into a single alignment.

The node age time priors were based on the posterior estimates from the associated species divergence times analysis (see ‘Dating species divergences’ above), using the span of the 95% highest density credibility intervals of node ages from that analysis to inform uniform time priors on the same species nodes in gene tree analysis, with a 1% probability tail that the maximum age could be exceeded. Calibrations within lineages that have undergone WGD were repeated across the duplicated clades with identical probability distributions. Molecular clock analyses were performed using the normal approximation method in MCMCtree^92^, with each gene treated as a separate partition. Four independent MCMC chains were run for 2 million generations each, with the first 20% discarded as burn-in. Convergence was determined using Tracer^71^ and by comparing congruence among all four runs. The alignments, MCMCtree control files and calibrations used are available within Supplementary File 20.

### Reconstruction of vertebrate ancestral chromosomes

Based on reciprocal best BLASTp (e-value threshold of 10e-6) search and a Chi-squared test (multiple test correction with FDR, *q* value threshold of 0.05), we identified homologous chromosomes within either chicken or spotted gar genome that possessed significantly more between-chromosome homologous genes. Homologous chromosomes between either chicken or spotted gar genome and sea cucumber chromosomes were also inferred, except that the best BLASTP search is unidirectional wherein the sea cucumber genes are the reference. From inferred within-species and between species homologies, all chicken and spotted gar chromosomes could be grouped into seventeen groups, which represent seventeen predicted ancestral chromosomes that contribute to extant gnathostome karyotype. The gene content of the ancestral chromosomes were complemented with Belcher’s lancelet genes. In definition, each vertebrate ancestral chromosome has its homologous relationship to specific sea cucumber, chicken and spotted gar chromosomes. A Belcher’s lancelet gene was distributed to one vertebrate ancestral chromosome if either (1) the scaffold this gene is located on is homologous to the specific sea cucumber chromosome and this gene is homologous to chicken and spotted gar genes; or (2) this gene is homologous to at least five different chicken and spotted gar genes.

### Phylogenetic support around 1R/2R

A homologous gene set is a group of genes that share the same best BLASTP hit AC gene. Multiple sequence alignments were obtained with PRANK^93^ v150803. ModelFinder^82^ (embedded in IQ-TREE^79^ v1.6.12) with BIC criteria and ‘-mtree’ parameter was used to find the best-fitting model. RAxML-ng v0.9.0 and IQ-TREE were repeatedly run for 10 times with different seed numbers. For the 20 obtained maximum likelihood trees, we used RAxML-ng to re-evaluate their likelihoods and chose the best tree as the final tree for each homologous gene set (Supplementary Files 5-8).

### Definition and calculation of overlapping ratio

For a reference chromosome and all genes on it, the existence or absence of homolog on a query chromosome is denoted as binary mode 1 or 0. We defined it as the gene retention profile. Mathematically, it is a vector with values of either 1 or 0 and with fixed length that corresponds to the number of genes on the query chromosome. One notable property towards gene retention profile is that gene order within the query chromosome does not alter the gene retention profile itself. Overlapping ratio is calculated between two gene retention profiles that correspond to one same reference chromosome. It is equal to the number of shared homologs divided by the smaller one of two total numbers of homologs. Overlapping ratio has a value range between 0 and 1. One notable property of overlapping ratio is that it is insensitive to the size difference between two query chromosomes.

### Hierarchical clustering based on gene retention profile

For multiple chromosomes homologous to one same vertebrate ancestral chromosome, we inferred their gene retention profiles and calculated all pairwise overlapping ratios. We used one minus overlapping ratio as a measure of pairwise distance and performed hierarchical clustering with the ‘Ward.D’ method provided by the R platform.

### GO enrichment analysis of ohnologs

We mapped hagfish and chicken ohnologs to human genes and performed GO enrichment analysis with human orthologs. Functional enrichment was examined with the Metascape^94^ online tool. We used the 959 human orthologs of hagfish ohnologs and randomly sampled 2,999 genes (as Metascape has a limit of 3,000) from a total of 3,595 chicken orthologs. GO (Biological Process) enrichment analysis was performed against all genes of the two species. Genes were annotated as either developmental ohnologs, non-developmental ohnologs or non-ohnologous genes according to their GO terms annotated by PANTHER^95^.

### Chromatin accessibility profiling

ATAC-seq experiments in two hagfish embryos, at stages Dean 45 (collected in 2018) and 53 (collected in 2017) (Supplementary Figure 28), were performed following previous descriptions^88, 96^ with slight variations (details are provided in Supplementary Information). Embryos were divided and processed into two halves to gain positional information to be used in a future project. Approximately 50,000 nuclei per replicate (∼200,000 nuclei per embryo) were processed for tagmentation using Tn5 from the Illumina Nextera DNA Library Prep Kit. Libraries were multiplexed and sequenced at the Beijing Genomic Institute in 4 lanes (2 per embryo) in an Illumina HiSeq 4000 platform.

To identify ACRs as putative gene *cis*-regulatory regions, we collected ATAC-seq data of hagfish and other chordate embryos (amphioxus, zebrafish, medaka, chicken, and mouse; GSE106428^40^ and DRA006971^97^ with two replicates each. For each data, ATAC-seq paired-end reads were aligned to the reference genome using Bowtie2^98^. After extracting nucleosome-free read pairs (the insert shorter than 120 bp), we performed peak-calling by using MACS2^99^. Finally, based on the replicate information, reproducible peaks were identified as ACRs by using the IDR framework^100^.

### Fate of ohnologs after WGD

After quantile normalization, TPM (transcripts per million) > 5 was used as a threshold to consider a gene to have either a ‘expressed’ or a ‘not expressed’ state. Only ohnolog pairs in which both genes are expressed in at least one tissue were analysed. Fates of ohnologs were classified according to their expressional patterns in the tissues assayed in this study for the hagfish, or from a previous study in the case of chicken^101^. Fates were defined following a different strategy than in Marlétaz et al.^40^ due to the lack of information from homologous tissues of the amphioxus to the ones assayed here for the hagfish. After WGD, ohnologs can follow one of the following fates: (i) redundancy, if the two ohnologs are expressed in the same set of tissues; subfunctionalization, if both ohnologs are each expressed in a tissue not shared with the other. In other words, each of them have tissue-specific expression domains; (iii) specializatoin, if one ohnolog has a reduced set of expression domains contained in a larger set of expressional tissues of the other ohnolog. Ohnolog families with a ‘specialization’ fate can be further defined as having a ‘strong specialization’ if the number of tissues in which one ohnolog is expressed is less than forty percent than that of the other ohnolog.

### Phenotypic disparity analyses

A character matrix of 578 characters and 278 taxa was assembled as follows. Characters were collected from direct observations and multiple literature sources^102–113^. Previous literature sources were modified to ensure that duplicated character states were removed and that overlapping characters from different sources were combined into single characters, or subdivided into multiple characters, so as to encompass all variation across vertebrates. We ensured that all characters were coded for as many taxa as possible. Missing data are coded as ‘?’; inapplicable characters are coded as ‘-’. Character state observations were coded using primary observations and through the literature. Characters were coded using hierarchical contingencies^114–116^. The character matrix and descriptions are available within Supplementary File 21 (Vertebrate_disparity_matrix.nex).

The phenotype character matrix was transformed prior to disparity analyses such that characters coded ‘not applicable’ were scored as ‘0’, and each subsequent character state was increased by 1. Ancestral character states were estimated along a tree representative of current phylogenetic hypotheses using stochastic character mapping^117^, with 1000 simulations per character; the tre is available in the Supplementary File 21 (Disparity.tre). Distances between each taxon and reconstructed internal nodes were estimated using Gower’s dissimilarity metric^118^, and these distances were ordinated using non-metric multidimensional scaling (NMDS), a method that seeks to reduce dimensionality while preserving distances between taxa. A pre-ordination phylomorphospace was plotted using the inferred ancestral states, NMDS scores and the representative phylogeny. Convex hulls were fitted around taxonomic lineages and groups that have undergone successive rounds of WGD. All stem gnathostomes were adjudged to have undergone the 2R WGD because they postdate the timing of 2R inferred from the gene tree based molecular clock analysis (see ‘Dating genome duplications in vertebrates’ above). Disparity metrics were estimated using dispRity with 1000 bootstrap replicates^119^.

### Reporting summary

Further information on research design is available in the Nature Portfolio Reporting Summary linked to this article.

### Data availability

Raw genome sequencing data and the genome assembly together with adult RNA-seq data have been deposited in the European Nucleotide Archive (ENA) at EMBL-EBI under accession number PRJEB21290. Supplementary files are available at FigShare (https://figshare.com/s/ca9e9b23cc36cfb59a44). A mirror of the UCSC Genome Browser containing hagfish assembly and annotations is available at http://ucsc.crg.eu/.

## Acknowledgements

This research was supported by grants from the Ministry of Science and Innovation of Spain (PID2021-125078NA-I00) and a KAKENHI Grant-in-aid for Scientific Research C from the Japan Society for the Promotion of Science (19K06798) to J.P.-A.; from the National Key R&D Program of China (2019YFA0802600), the Chinese Academy of Sciences (ZDBS-LY-SM005), the National Natural Science Foundation of China (31970565) and the Open Research Program of the Chinese Institute for Brain Research (Beijing) to Y.E.Z.; the Strategic Priority Research Program of the Chinese Academy of Sciences (XDB13000000) to Y.E.Z and Wen W.; the Wellcome Trust (to F.J.M., Grant number 108749/Z/15/Z); the Leverhulme Trust (RF-2022-167 to P.C.J.D); the Biotechnology and Biological Sciences Research Council (BB/T012773/1 to P.C.J.D); the Natural Environment Research Council (NERC) grant (NE/P013678/1 to P.C.J.D. and D.P.), part of the Biosphere Evolution, Transitions and Resilience (BETR) programme, which is co-funded by the Natural Science Foundation of China (NSFC); the John Templeton Foundation (Grant 62220 to P.C.J.D. and D.P.; the opinions expressed in this publication are those of the author(s) and do not necessarily reflect the views of the John Templeton Foundation); Wellcome Trust Seed Award (210101/Z/18/Z to J.P.). The computing was jointly supported by the HPC Platform of BIG and that of the Scientific Information Centre of IOZ. We thank Capt. Osamu Kakitani and the members of the Fishery Association in Gotsu City, Shimane Prefecture, Japan, for his assistance in collecting hagfish; the technical support staff of the Laboratory for Phyloinformatics in RIKEN Kobe for sequencing data production, and RIKEN aquarium staff. We thank Tamara de Dios Fernández for animal illustrations.For the purpose of open access, the authors have applied a CC BY public copyright licence to any Author Accepted Manuscript version arising from this submission.

## Author contributions

D.Y., Y.R., J.S., Y.L., W. Wan, Wen W., Y.E.Z. and J.P.A sequenced, assembled and evaluated the hagfish genome. F.J.M. generated and coordinated annotation process at Ensembl. M.M., C.C., M.P. and W.A. generated Ensembl Compara data. A.d.M., D.Y., A.J.S.B. and J.W.C. generated orthogroups. R.F., J.F.F., F.S., S.D’A., C.B., C.A., G.S., B.F., K.J.P., S.D., M.H., J.P.R., M.D.C., contributed to gene family curation. A.J.S.B., J.W.C., D.P., and P.C.J.D. performed phylogenomics and dating analyses. J.P. performed analysis of gene family evolution. D.Y., J.S. and Y.E.Z performed macrosynteny analysis. I.M. and M.I. advised on macrosynteny and ohnolog evolution analyses and discussed the data. J.P.-A, I.S., F.S. and S.K. obtained hagfish embryonic and adult material. J.P.-A performed ATAC-seq and RNA-seq experiments and ensured sequencing project management with BGI and RIKEN. M.U. and J.S. carried out regulatory genome profiling and ohnolog fate evolution analyses. A.J.S.B., J.W.C., J.N.K., E.M.C., R.P.D., S.G., E.R., R.S., D.P., and P.C.J.D. obtained fossil data, calibration data and performed morphological disparity analysis. Wen W., S.K., N.I. and J.P.-A. conceived the project. S.K., N.I., P.C.J.D., Wen W., Y.E.Z and J.P.-A contributed to project design. J.P.-A. coordinated the project. Y.E.Z., P.C.J.D and J.P.-A. wrote the manuscript, with inputs from all authors. All authors revised and approved the manuscript.

## Competing interests

The authors declare no competing interests.

## Additional information

**Supplementary information** The online version contains supplementary material available at https://doi.org/xx.xxxx/sxxxxx-xxx-xxxxx-xx.

**Correspondence and requests for materials** should be addressed to Juan Pascual-Anaya.

**Peer review information** TBD

**Reprints and permission information** is available at http://www.nature.com/reprints

---

**Extended Data Figure 1.**
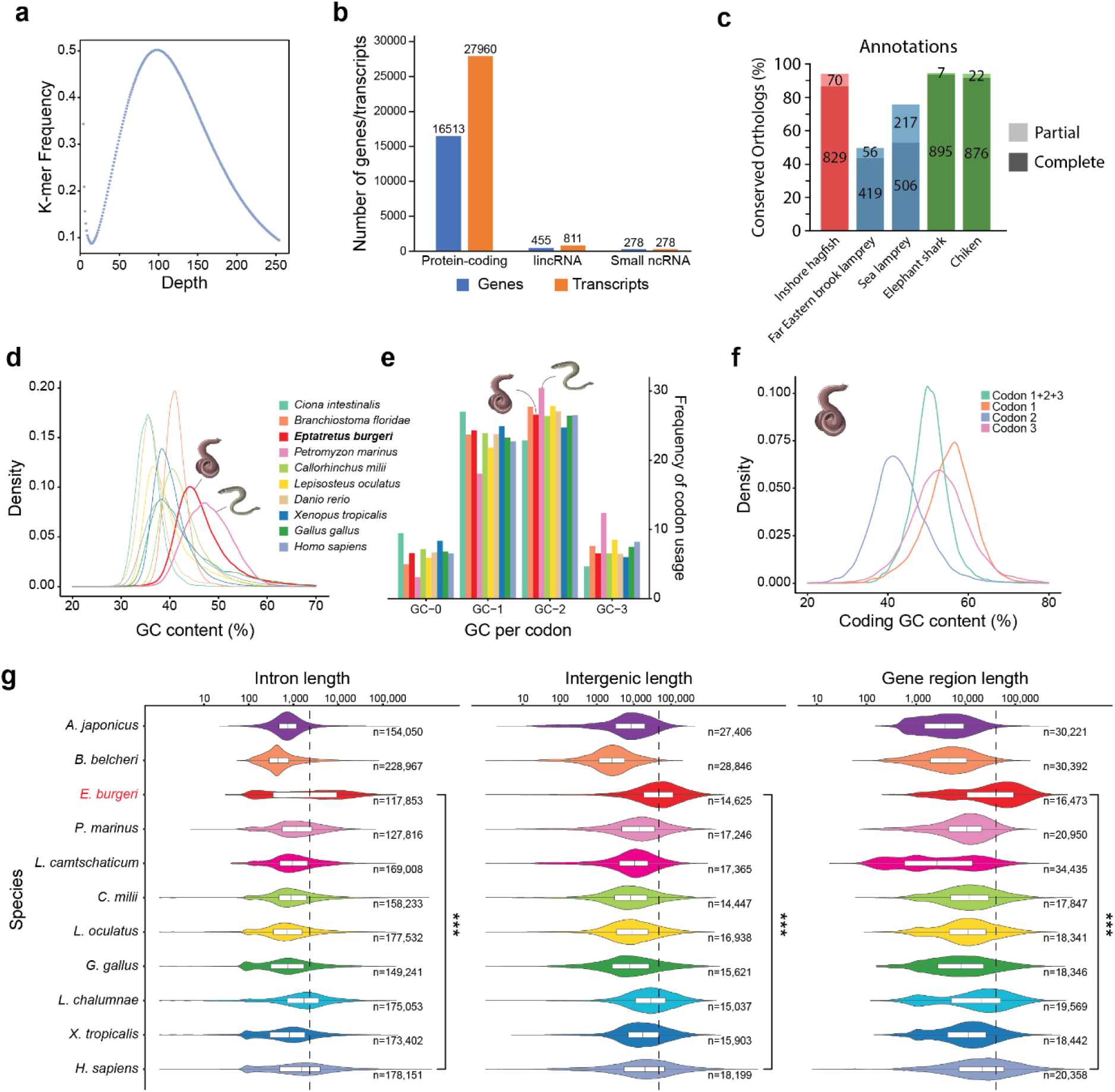
Features of the hagfish genome. **a**, 17-mer distribution for inshore hagfish genome size estimation using all raw reads from short insert-size libraries. **b**, Counts for major classes of genes and transcripts from Ensembl annotation. **c**, Completeness assessment of the annotation of the inshore hagfish *E. burgeri* genome (red), three lamprey species (blue) and two jawed vertebrates (green). Numbers of conserved metazoan orthologs (metazoa_odb10 dataset, n =954) are indicated for each case. **d**, GC-content distribution of the hagfish genome and other chordates calculated from 10-kb non-overlapping windows. **e**, All codon type frequency in given chordate genomes according to GC-content. GC-0/1/2/3 indicates the number of G or C bases in a codon. **f**, Distribution of GC content at each codon position or at all codon positions (Codon1+2+3). For each protein coding gene, we only kept the longest coding sequence. For each coding sequence, we calculated the GC content at separate codon positions. We also calculated the GC content for each coding sequence, which is equal to the GC content of all three codon positions. **g**, Violin plots of size distribution of intron (left), intergenic (middle) and gene body (right) lengths over a logarithmic scale of the hagfish (*E. burgeri*), two lamprey species (sea lamprey, *P. marinus*; Arctic lamprey, *L. camtschaticum*), six gnathostome vertebrates (human, *H. sapiens*; frog, *X. tropicalis*; coelacanth, *L. chalumnae*; chicken, *G. gallus*; spotted gar, *L. oculatus*; and the elephant shark, *C. milii*) and two invertebrate deuterostomes (sea cucumber, *A. japonicus*; amphioxus, *B. belcheri*). For each genomic feature and each species, the median and IQR (interquartile range) length statistics are indicated with a white rectangle. Dashed vertical line indicates median size of *E. burgeri* features; *** *P* < 2.2 × 10^−16^, two-sided Wilcoxon sum-rank test.

**Extended Data Figure 2.**
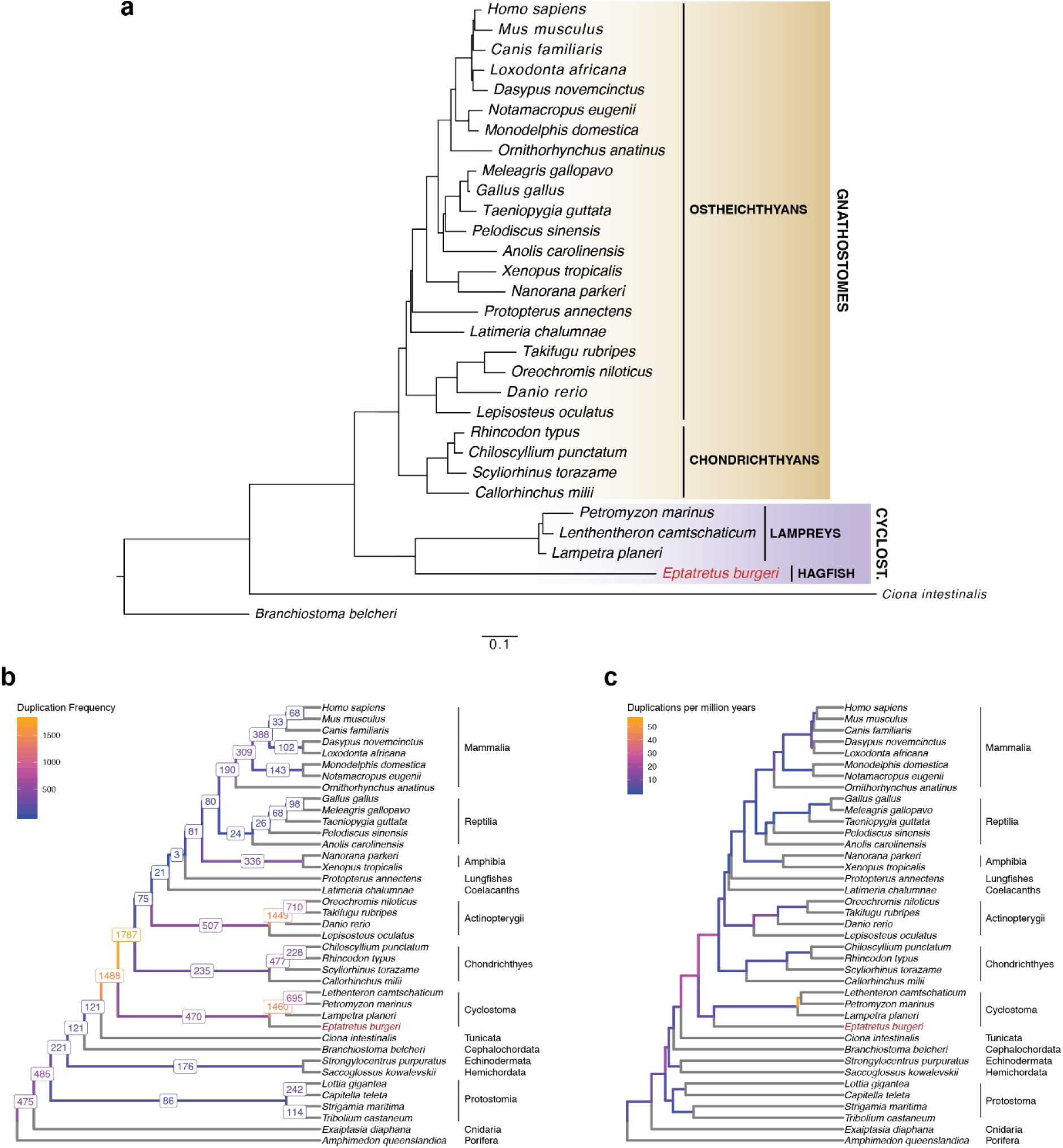
Phylogenomic analysis of chordates and gene duplication rates across metazoan evolution. **a,** Bayesian Inference tree of 31 chordate species was built using a protein alignment (see Methods). All nodes were recovered with a posterior probability of 1. Cyclostome monophyly was unequivocally supported. Scale bar indicates 0.1 substitutions per site. **b, c,** The exact number of gene duplication events inferred using OrthoFinder2 with greater than 50% support (**b**) and the number of events per million years per branch (**c**) are shown. In each case, the colour of the branch represents the value according to the key on the upper-left of each pane. Hagfish is highlighted in red font in all panes.

**Extended Data Figure 3.**
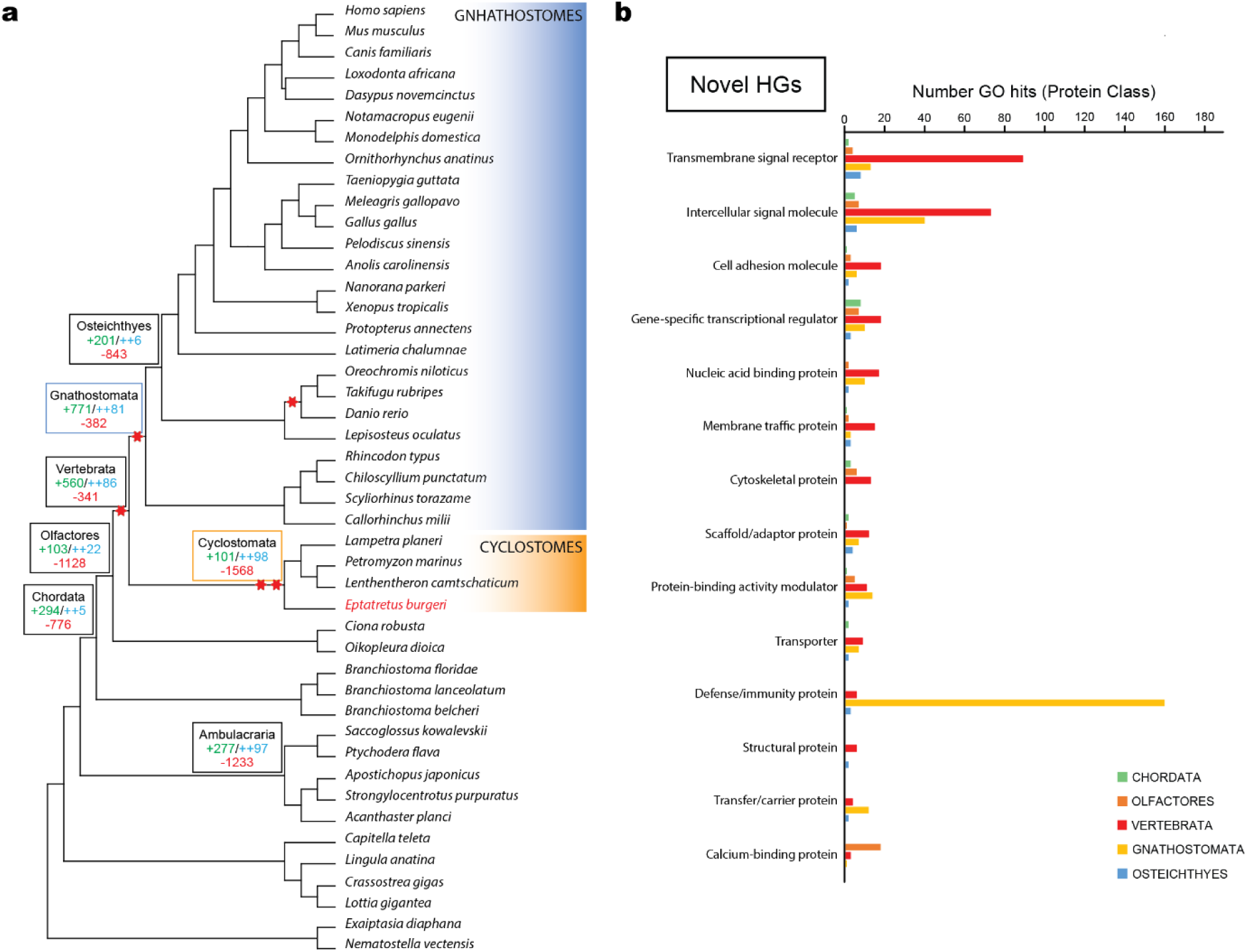
Reconstruction of ancestral gene content. **a**, Cladogram showing the phylogenetic relationships of 45 species representatives of all major eumetazoan taxa with species of gnathostomes and cyclostomes highlighted in blue and orange, respectively. Gene family gains and losses are indicated in selected nodes: green, novel homology groups (HG); blue, novel core HGs; red, lost HGs. **b**, Top 14 Protein Class GO hits for novel homology groups (HG) gained across different nodes of chordates, color coded by taxa (legend at the bottom right) and sorted by the Vertebrata node. The largest GO enriched terms are ‘transmembrante signal receptor’ and ‘intercellular signal molecule’ in vertebrates, and ‘defense/immunity protein’ in gnathostomes.

**Extended Data Figure 4.**
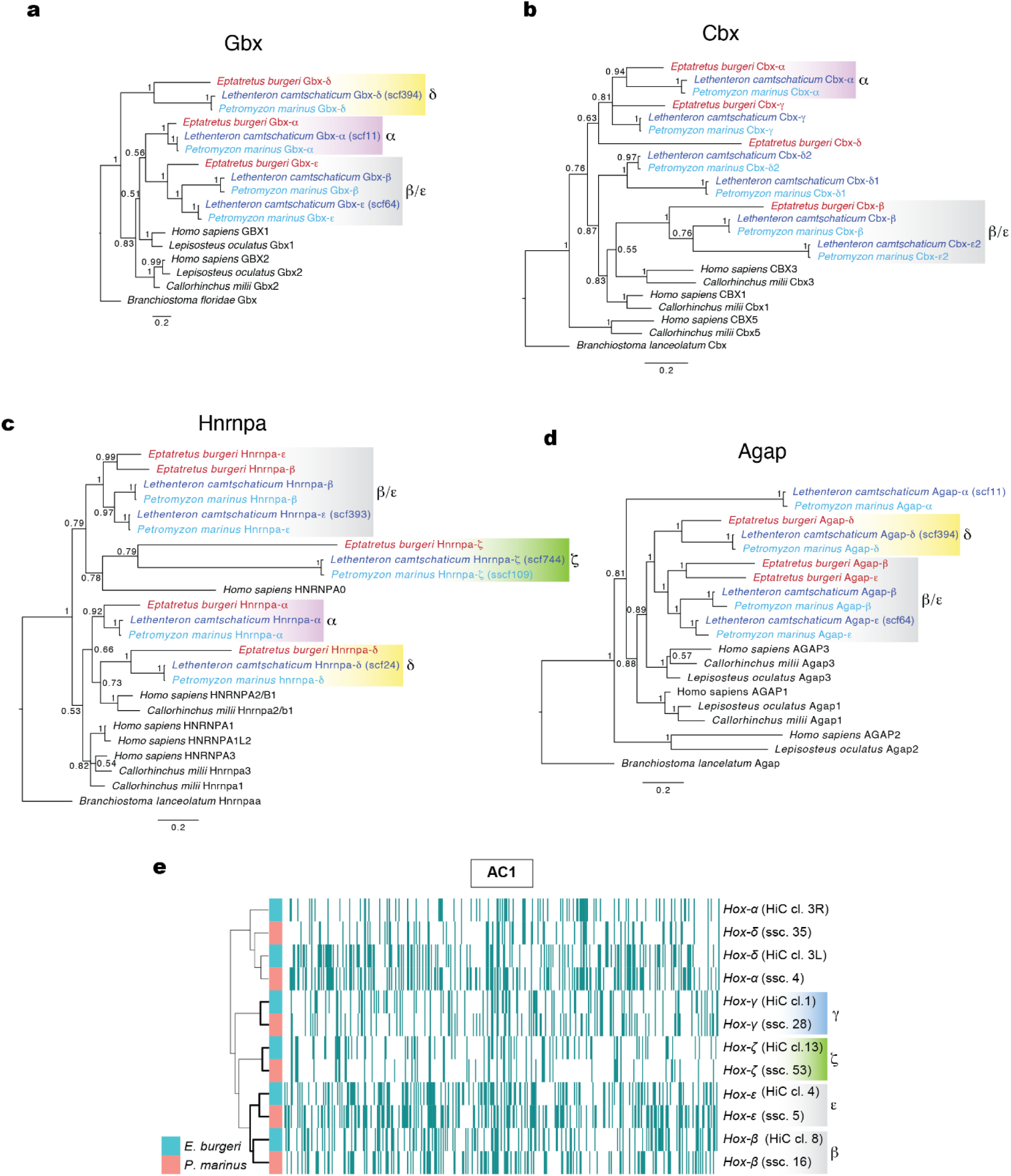
Phylogenetic and retention profile clustering analyses of Hox syntenic regions. **a-d,** Bayesian inference phylogenetic trees of amino acid sequences of 4 non-*Hox* syntenic genes to *Hox* clusters, Gbx (**a**), Cbx (**b**), Hnrnpa (**c**) and Agap (**d**), of the inshore hagfish (in red), the sea lamprey (in light blue), the Arctic lamprey (in dark blue) and selected gnathostomes. Orthologs from the European amphioxus *Branchiostoma lanceolatum* were used as outgroup to root the trees. Posterior probability is indicated in each node. Scales indicate number of substitutions per site. Phylogenetic analyses of *Hox* genes generally fail to determine orthology due their high conservation and short alignments. The phylogenetic trees of these non-*Hox* linked genes clearly support the orthology of *Hox-α* (Gbx, Cbx and Hnrnpa), *Hox-δ* (Gbx, Hnrnpa and Agap), and *Hox-ζ* (Hnrnpa) clusters, while *β* and *ε* genes always group together, as previously observed for the lamprey^17^. The alignments used to build the trees, together with the MrBayes parameters and number of generations used to build each tree are provided as Supplementary Files 16-19. **e**, clustering analysis of retention profiles (see main text) resolved the orthology relationships of *Hox-β*, *Hox-ε*, *Hox-γ*, as well as *Hox-ζ* clusters. Supported orthologies in each analysis are marked with color-coded rectangles. The location of each cluster is indicated in parenthesis in e (ssc, super scaffold; HiC cl, Hi-C contact cluster, or chromosome). For the clustering analysis of AC1-derived chromosomes in the lamprey and hagfish, we split Hi-C cluster 3 into two halves, each containing one Hox cluster: 3L (coordinates 0-107.78 Mb), 3R (107.78-194 Mb).

**Extended Data Figure 5.**
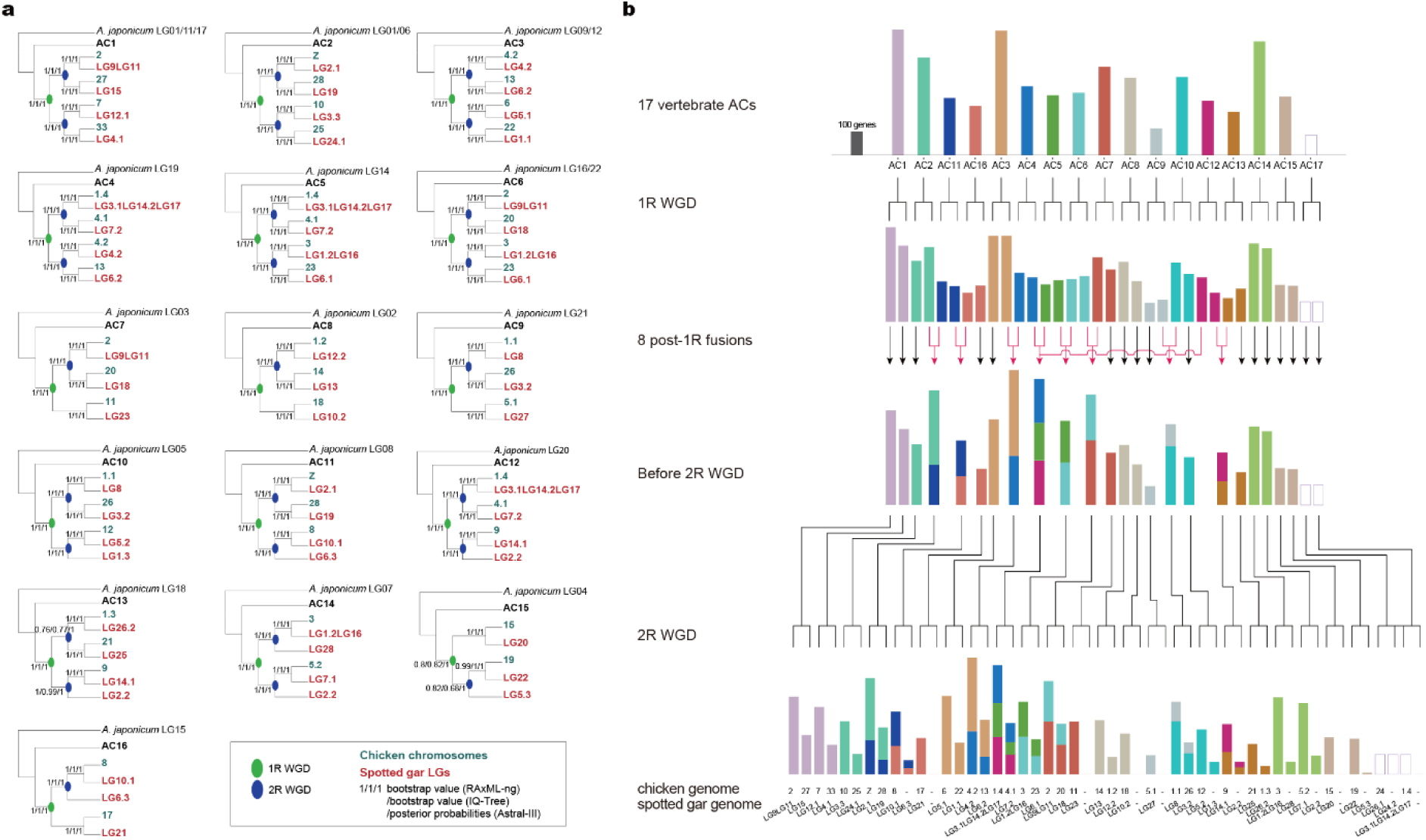
Emergence of gnathostome karyotypes via 2R. **a**, Chromosomal level phylogenetic trees demonstrate the occurrence of 2R WGD. The two rounds of WGD are color-coded. Cyan and red denote genes encoded by corresponding chicken chromosomes and gar LGs, respectively. Bootstrap support and posterior probability values of three methods are marked on branches. **b**, Pre- and post-1R chromosomal fusions contributed to the evolution of ancestral gnathostome karyotypes. The size of each ancestral chromosome, except for AC17 contributed chromosomes (see Supplementry Information), is proportional to the number of retained genes from ACs. The pre-1R fusion leading to AC3 is drawn as a dotted line due to its uncertainty.

**Extended Data Fig. 6.**
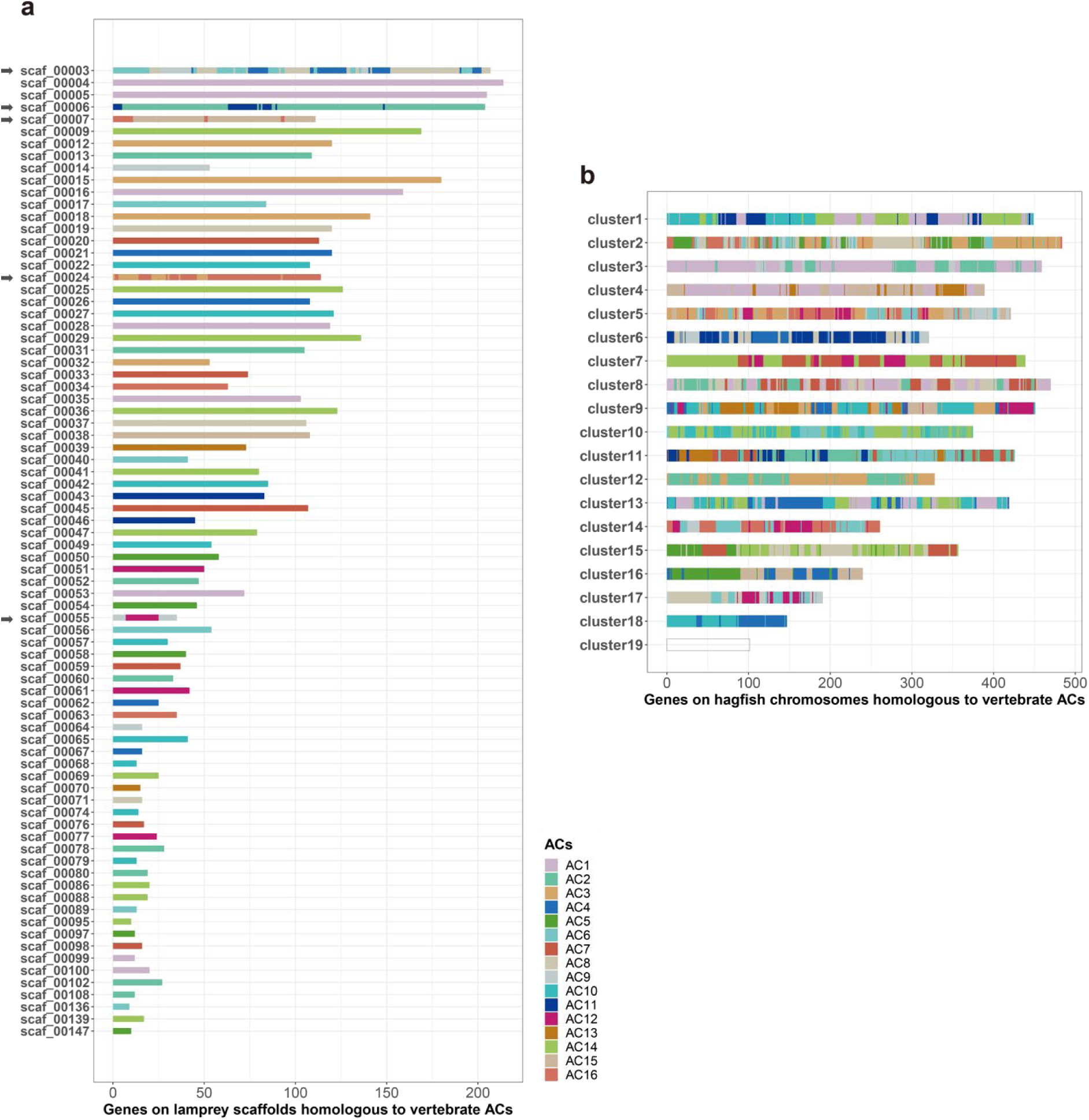
Contributions of vertebrate ACs to the genomes of hagfish and lamprey. **a**, Sea Lamprey scaffolds are generally homologous to one single AC except five scaffolds labeled with an arrow. Sea lamprey scaffolds including scaf_00001, scaf_00002, scaf_00008, scaf_00010, scaf_00011 and scaf_00023, confounded by missassembly, are not presented here. **b**, The distribution of homologous genes to ACs on 19 hagfish cluster. Only significant homologous relationships between hagfish clusters and ACs are shown. Because hagfish cluster 19 is not homologous to any AC, it is presented as a blank block. Genes are color-coded according to its homologous AC.

**Extended Data Figure 7.**
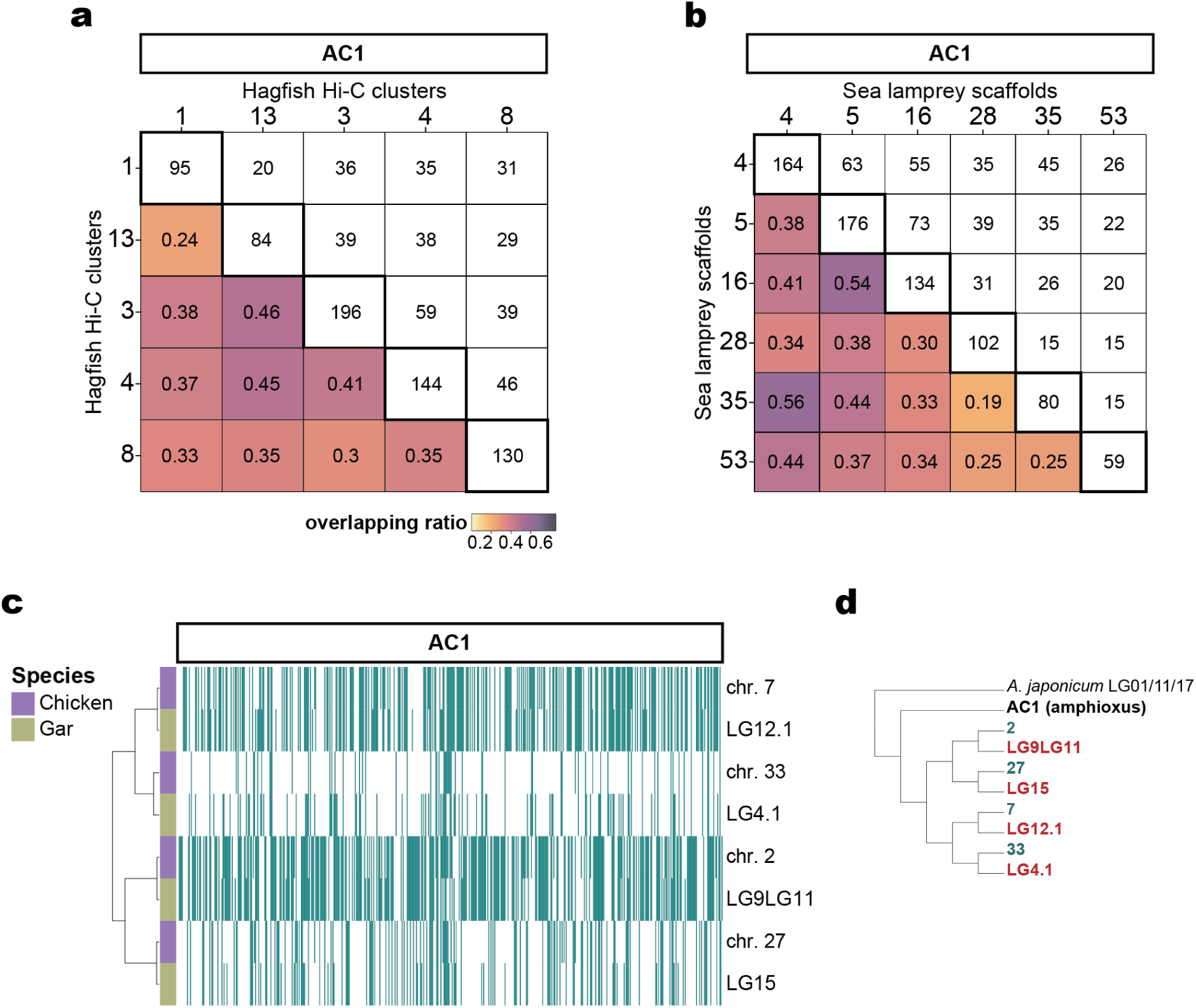
Overlapping ratio and clustering analysis of ohnologous and orthologous chromosomes. **a**,**b**, AC1 corresponds to five and six mutually paralogous chromosomes in hagfish (**a**) and sea lamprey (**b**) genomes, respectively. Numbers in colour-coded cells (bottom left triangle) indicate the *OR* between two chromosomes. Numbers in white cells (top right triangle) indicate the number of shared retained genes between two chromosomes. Numbers on the diagonal line from top left to bottom right (thick-lined cells) indicate the total number of retained genes of a chromosome. Overlapping rations corresponding to all hagfish chromosomes and lamprey scaffolds are provided in Supplementary File 9. **c**, Retention profile clustering analysis of gnathostome orthologous chromosomes deriving from AC1. Retained genes are denoted by dark cyan lines. Four orthologous chromosome pairs are defined. **d**, The clustering found in (**c**) is the same as that found in the phylogenetic analysis (Extended Data Fig. 5a), demonstrating the reliability of the *OR* approach.

**Extended Data Figure 8.**
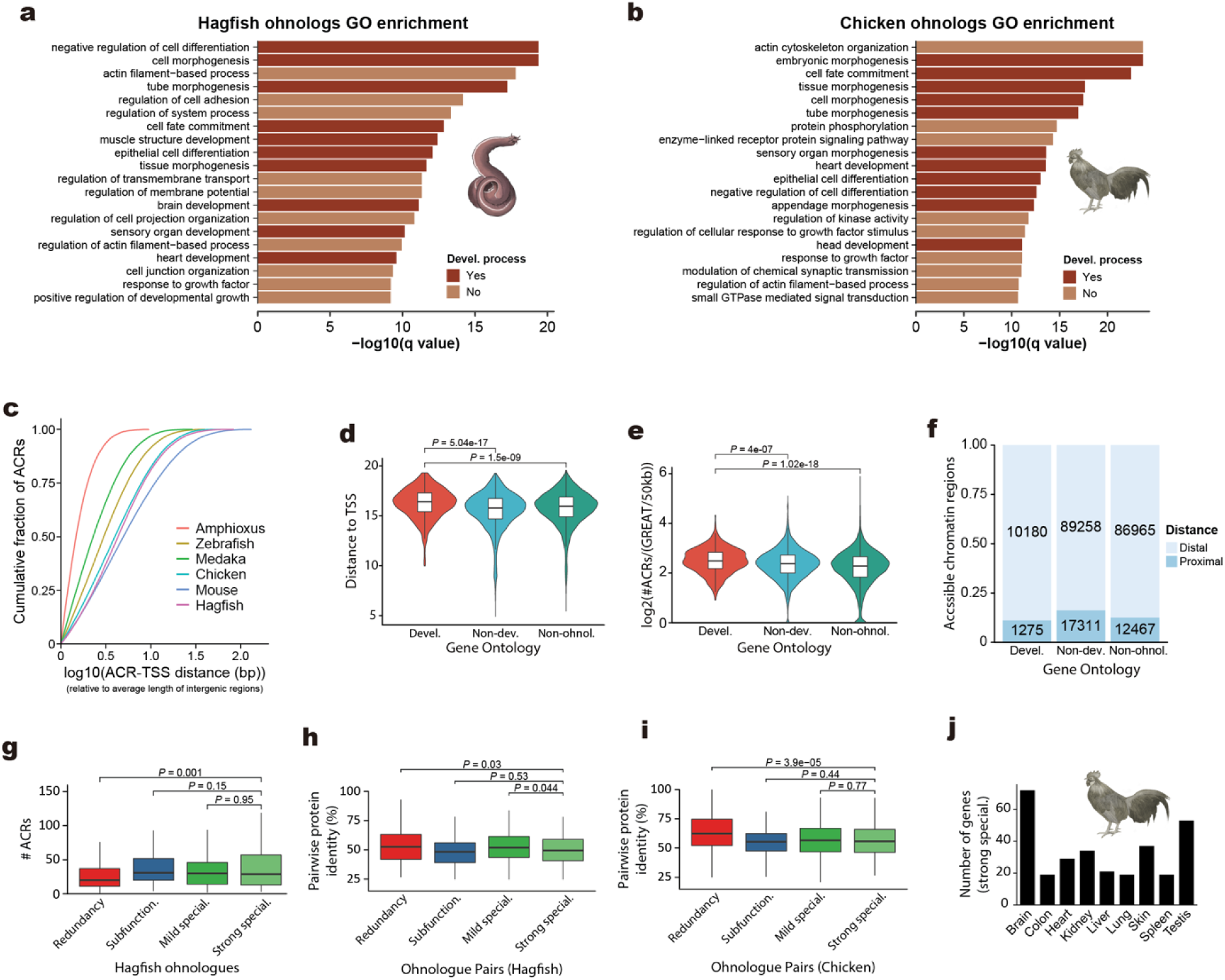
Fate and *cis*-regulatory evolution of ohnologs after WGD. **a**, **b**, Gene Ontology enrichment analysis of ohnologs in the hagfish (**a**) and the chicken (**b**). Top 20 terms are shown, majority of which are related with development. **c**, Cumulative distribution of distance of ACRs from the closest TSSs normalized by the average length of intergenic regions in each genome. **d**, Distribution of the distance from ACRs to closest TSS of developmental ohnologs (Devel.), non-developmental ohnologs (Non-dev.) and non-ohnologous (Non-ohnol.) genes. **e**, Distribution of the number of ACRs per gene, normalized by the GREAT region length, for developmental ohnologs (Devel.), non-developmental ohnologs (Non-dev.) and non-ohnologous (Non-ohnol.) genes. **f**, Proportion of distal ACRs across different gene functional categories. Within a GREAT-defined region, proximal regulatory sequences were defined as those from 5 kb upstream to 1 kb downstream of a TSS, and the rest of the region was treated as distal. **g**, Distribution of the number of ACRs of hagfish ohnologs for each category (special., specialization). **h**, Distribution of pairwise protein identity for hagfish ohnologous pairs for each category. **i**, Distribution of pairwise protein identity for chicken ohnologous pairs for each category. **j**, Number of ohnologues with strong specialization in chicken expressed in each tissue. Only the gene in a pair with narrower expression breadth is analyzed. *P* values in **d**, **e**, **g**, **h**, **i** correspond to two-sided Wilcoxon sum-rank test between the indicated groups.

## Notes

### Competing Interest Statement

The authors have declared no competing interest.

