## Supplementary Information, including Supplementary Figures 1-31 for "Hagfish genome illuminates vertebrate whole genome duplications and their evolutionary consequences"

#### 1. ASSEMBLY AND ANNOTATION OF THE HAGFISH GENOME

##### 1.1) Hagfish genome assembly.

**1.1.1) Genome sequencing.** We extracted high-molecular-weight genomic DNA from the testis of a single, sexually mature individual of the inshore hagfish, *Eptatretus burgeri,* following a standard phenol-chloroform and Proteinase K extraction protocol for high-molecular-weight DNA (Chapter 6, Protocol 1 from Sambrook and Russell[1]) and was used to prepare a total of six paired-end libraries (IDs MM1-MM5 and PE250, with insert size means of 174, 234, 242, 279, 612 and 300 base pairs (bp), respectively) and five long-insert, mate-pair libraries (IDs MM6-MM10, with insert size means of 5, 6, 8.5, 12.5 and 17.5 kilobases (kb)) (Supplementary Table 2). These libraries were sequenced using Illumina HiSeq 2500 (libraries MM1-MM5) and Illumina MiSeq (library PE250) platforms, generating over 753 Gb (Gigabase) of clean data after some filtering (see below), representing ~240X of sequencing coverage (based on an estimated size of 3.12 Gb, see below) (Supplementary Table 2).

**1.1.2) Filtering and correcting raw reads.** Low quality reads from paired-end (MM1-MM5) and mate-pair (MM6-MM10) data were removed with an in-house Perl script. For mate-pair data, reads were shortened 5 bp at the 5' end, and 20 bp at the 3' end. Then, read pairs with more than 30 low-quality bases (Illumina quality score < 7) or more than 20% of unknown bases in at least one read were discarded. For paired-end data, reads were shortened 2 bp at both 5' and 3' end. Read pairs with more than 30 low-quality bases or more than 5% unknown bases in at least one read were discarded. Then, all reads were filtered for duplications and adapter sequences were discarded. Finally, short-insert-size library reads data (MM1-MM5) were corrected with SOAPec v. 2.03

(Luo et al., 2012), with the following parameters: `-k 21 -i 500000000 -g 120000000000`. The basic sequencing, filtering and correction stats are shown in Supplementary Tables 2, 3. For the PE250 library with the highest sequencing length (paired end 250 bp) but moderate mean insertion size (300 bp), we applied FLASH software with default parameters to merge a pair of reads into one[2]. As a result, a median length of 385 bp single end library was obtained (Supplementary Table 2).

**1.1.3) Genome size estimation.** We estimated the genome size of *E. burgeri* to be 3.12 Gb using a k-mer distribution of all 17-mers using sequencing data from the five short-insert paired-end libraries (libraries MM1 to MM5; Extended Data Figure 1a, Supplementary Table 4) following previously reported methods[3,4]. Briefly, we calculated the estimated genome size by dividing the total base pairs used for the analysis (filtered reads) by the real sequencing depth (N). N is calculated using the formula $N = (M * L) / (L – k + 1)$ (REF. 3), where L is the average read length, k the k-mer size and M the k-mer coverage at the maximum frequency of the 17-mer distribution (Extended Data Figure 1a). Our size estimation was in line with previous studies[5] and within the range of other hagfish species[6].

**1.1.4) Primary genome assembly and Chicago (Dovetail) scaffolding.** For whole genome sequence assembly, filtered and corrected reads (MM1-MM5, PE250) were firstly assembled into contigs using ABySS[7] (v1.9.055) with a k-mer size of 79. Contigs were assembled into scaffolds with SOAPdenovo2 (REF. 8) (v2.04-r24154, parameters '`-K 41 -d 1 -M 2 -F`') after remapping reads from libraries MM6-MM10 to contigs. Finally, we retrieved the read pairs wherein one end was uniquely mapped to one contig and the other end was located in a gap region, and performed a local assembly for these collected reads to fill the gaps. This gap filling process was performed with GapCloser v1.12-r6, from the SOAPdenovo2 package[8]. To further improve the large-scale structure of this assembly, long-range ligation data from three Chicago libraries were used to correct and improve the scaffolding of the assembly, using the HiRise pipeline at Dovetail Genomics[9] (Supplementary Table 5). This step increased the contiguity of our assembly by 7-fold, increasing the scaffold N50 from 439 kb to 2.69 Mb, and reducing the number of scaffolds >1kb from 21,115 to 5,465 (Supplementary Table 6). We assembled ~2.60 Gb of the 3.12-Gb estimated size, probably missing hnighly repetitive, difficult-to-sequence microchromosomes[10,11] (see below). This assembly was polished using Pilon[12] (version 1.22) using all data from short-insert libraries and mapped to the genome with Bowtie2 (REF. 13). The resulting corrected genome assembly, version 3.2 in our pipeline, was first publicly opened in Ensembl (http://www.ensembl.org/Eptatretus_burgeri), release 93, in July 2018 (https://www.ensembl.info/2018/07/17/ensembl-93-has-been-released/), with the goal of providing the scientific community with early access to the genome[14].

**1.1.5) Generation of a Hi-C guided hagfish assembly.** We next decided to generate a chromosome-level assembly to enable macrosynteny comparative analysis, crucial to resolve the timing of whole genome duplication (WGD) events in early vertebrate evolution. We generated a Hi-C library from DNA extracted from the testis of a second male individual following a previously described method[15]. We sequenced 2,380 million Hi-C read pairs with an Illumina HiSeq X Ten platform and mapped them to the draft assembly with the HiCUP (version 0.7.2) pipeline[16]. We retained 363 million (15% of the library) valid read pairs, which corresponds to approximately 350X genomic coverage. In a previous study, for a draft human genome assembly with N50 of 437 kb, 240 million retained Hi-C read pairs lead to a new assembly with the clustering error rate below 0.2% and the ordering or orienting error below 2% (REF. 17). Since the *E. burgeri* genome has a similar size than the human genome (3.12 Gb vs 3.1 Gb, respectively) and our

Dovetail assembly already reached a N50 of 2.69 Mb, we expected our Hi-C data to be enough to achieve a high-quality chromosome-level assembly for *E. burgeri*. Since small scaffolds cause serious computing load and are difficult to cluster due to limited Hi-C read pairs mapped to them, we only analyzed 1,635 scaffolds longer than 50 kb, which constitute 99.1% (2,583,606,940/2,608,383,542) of the version 3.2 genome assembly (Dovetail). After loading read pairs mapped to these 1,635 scaffolds into the Hi-C assembly software LACHESIS[17] (compiled on Apr 19, 2019), we tried different parameters in order to optimize the coverage toward the Dovetail assembly. We first used `CLUSTER_N = 18` and `CLUSTER_N = 26`, which represent somatic and germline haploid chromosome number, respectively[18]. Note that although we used a testis sample in order to get germline DNA, it actually represents a mixture of both somatic and germline cells (from the presence of blood vessels and connective tissue of testis). The Hi-C read counts were normalised by a square root vanilla coverage normalisation method[19]. Then it was used as input to draw a heatmap plot in R via the ggplot2 package. With this heatmap, we found that one cluster obviously consists of two separate Hi-C contact blocks in case of `CLUSTER_N = 18`. For `CLUSTER_N = 26`, multiple clusters show extensive between-cluster Hi-C contacts. So, we adjusted the parameters to ‘`CLUSTER_N = 19 CLUSTER_MIN_RE_SITES = 200 CLUSTER_MAX_LINK_DENSITY = 3.6 CLUSTER_NONINFORMATIVE_RATIO = 6 ORDER_MIN_N_RES_IN_TRUNK = 100 ORDER_MIN_N_RES_IN_SHREDS = 50`’ and assigned 96.2% (1573/1635) scaffolds longer than 50 kb into 19 clusters (preliminary Hi-C assembly; Supplementary Figure 1).

Next, by integrating different strategies to identify chromosomal rearrangements or misassemblies[17,20,21], and to polish likely Dovetail missassemblies (Supplementary Figure 1), we divided each scaffold into 200 kb sized segments and treated scaffolds smaller than 200 kb as a single segment, and then developed a hybrid strategy to detect and correct assembly errors based on three types of Hi-C contact information (see Supplementary Figure 2 for an example). First, we examined how a segment interacted with its original cluster and other clusters. To do so, we calculated the read pair count between any segment pair. For a segment of interest, we calculated the cluster level interaction intensity as the median of this segment relative to all segments harboured by one cluster of interest. For any 200 kb-segment, if its highest Hi-C contact occurs in another cluster not associated with its scaffold (Supplementary Figure 2), the corresponding scaffold could be misassembled. Second, for each candidate misassembled scaffold, the read pair densities between this scaffold and two clusters (only the cluster the scaffold belongs to and the other cluster that show highest Hi-C contact with segments of the scaffold) would be partitioned where different segments interact with two cluster differentially (Supplementary Figure 2). Third, for each candidate misassembled scaffold, the read pair densities within this scaffold would be partitioned too where more than one Hi-C interaction block would emerge (Supplementary Figure 2). With these three lines of information, we identified 280 scaffolds associated with 364 misassembly boundaries since a single scaffold could harbour two or more misjoinings (Supplementary Table 7; Supplementary File 1). Given the boundary coordinates, we split these scaffolds into smaller scaffolds and performed a second round of Hi-C based assembly with the same parameters.

Given Hi-C interaction heatmaps, we examined orders of scaffolds within a cluster and performed adjustment in 10 clusters, including cluster1, cluster2, cluster3, cluster5, cluster6, cluster7, cluster8, cluster9, cluster10 and cluster11. We assigned 98.7% (1873/1999) of all corrected scaffolds longer than 50 kb to 19 clusters (main Figure 1d; Supplementary Tables 8, 9). These scaffolds constitute 98.4% (2,578,071,452/2,608,383,542) of the Dovetail hagfish draft assembly in size and include 99.1% (16363/16513) of the annotated protein coding genes. Accordingly, we translated the gene coordinates from the Dovetail to the Hi-C assembly and excluded 41 genes that overlapped with boundaries of misjoins (Supplementary Table 10). This corrected assembly (v4.0 in our pipeline) served as the final assembly used in the main text.

**1.1.6) Investigation on hagfish Hi-C cluster 19.** The Hi-C contact cluster 19 is the smallest cluster in size (~32 Mb; Figure 1d, Supplementary Table 9). Initially, since the diploid number of chromosomes of *E. burgeri* somatic cells is 2n=36 (haploid n=18) after the elimination of germline-specific 8 pairs of chromosomes (2n=16) (REF. 10), we speculated that cluster 19 and one other Hi-C contact cluster may represent a pair of sex chromosomes, suggesting the presence of XY sexual chromosomes in *E. burgeri*'s male sex. The sex determination system of hagfishes is mostly unknown (reviewed by Stöck et al.[22]). If two of the 19 hagfish Hi-C clusters represent in fact sexual chromosomes, their absolute DNA content should be each half of that from an autosomal chromosome. Accordingly, read coverage of the sexual chromosomes is thus expected to be approximately half of the remaining chromosomes. In order to calculate read coverage across Hi-C clusters, we randomly selected raw data from the genome sequencing (MM-1 lane 1, MM-2 lane 2, MM-5 lane 4 and MM-5 lane 4_2, Supplementary Table 3) and measured the read depth of uniquely mapped reads to each cluster of the Hi-C assembly. DNA sequencing reads were mapped to the hagfish Hi-C genome assembly using BWA[23], version 0.7.2-r351. Only reads with mapping quality exceeding 30 are reported. Base level read coverage was calculated with Bedtools genomecov tool[24], version 2.25.0. As a result, we found that all 19 clusters have a comparable read depth (Supplementary Table 9), making it unlikely that two of the 19 clusters are a pair of XY sex chromosomes. Compared with the remaining clusters, Hi-C cluster 19 depicts the most uneven distribution of read coverage along its base pairs (Supplementary Figure 3), probably because it contains the highest proportion of N's in its sequence (73% v.s. 42-45%; Supplementary Table 9), indicating that it might consist mostly of difficult-to-assemble, highly repetitive sequences, especially when using only short read data. In conclusion, Hi-C cluster 19 most likely represents a germline-specific, difficult-to-assemble microchromosome[10,11].

**1.2) Ensembl.**

**1.2.1) Ensembl annotation (Vertebrate Annotation team).** The detailed process of annotation of the hagfish genome can be found in Ensembl site: http://www.ensembl.org/info/genome/genebuild/2018_06_eptatretus_burgeri_genebuild.pdf.

***1.2.1.1) Methods.*** A set of potential transcripts was generated using two major techniques: primarily through alignment of short read RNA-seq data and also through gap filling with protein-to-genome alignments of a subset of vertebrate proteins with experimental evidence from UniProt (The UniProt Consortium, 2019) vertebrate proteins. The short-read RNA-seq data were sourced from samples generated as part of this study (uploaded to ENA under Study accession number PRJEB21290) from blood, brain, gills, heart, intestine, kidneys, liver, testes, skeletal muscle. Data from a second study (PRJNA371391) covering development stages at days 30, 35 and 40 were also used[25]. The UniProt vertebrate proteins had experimental evidence for existence at the protein or transcript level, termed protein existence (PE) level 1 and 2 proteins. Due to the high evolutionary distance from most species to hagfish, all vertebrate PE1/2 proteins were downloaded and aligned in order to increase sensitivity for detecting coding exons. At each locus, low quality transcript models were removed, and the data were collapsed and consolidated into a final gene model plus its associated non-redundant transcript set. When collapsing the data, priority was given to models derived from transcriptomic data. For each putative transcript, the coverage of the longest open reading frame was assessed in relation to known vertebrate proteins, to help differentiate between true isoforms and fragments. In loci where the RNA-seq data were fragmented or missing, homology data took precedence, with preference given to longer transcripts that had strong intron support from the short read data. Gene models were classified, based on the alignment quality of their supporting evidence, into three main types: protein-coding, pseudogene, and long non-coding RNA. Models with hits to known proteins, and few structural abnormalities (i.e. they had canonical splice sites, introns passing a minimum size threshold, low level of repeat coverage) were classified as protein-coding. Models with hits to known protein, but having multiple issues in their underlying structure, were classified as pseudogenes. Single-exon models with a corresponding multi-exon copy elsewhere in the genome were classified as processed pseudogenes. Models with transcriptomic support, but no evidence of a protein-coding ORF were listed as potential lncRNAs. Potential lncRNAs were further filtered to remove transcripts that had overlap with a protein coding gene (regardless of strand) or if the transcript had a signature of a protein domain. Single exon lncRNAs were filtered out due to lack of evidence to separate them from background transcriptional noise. A separate pipeline was run to annotate small non-coding genes. miRNAs were annotated via a BLAST[26] of miRbase[27] against the genome, before passing the results into RNAfold[28]. Poor quality and repeat-ridden alignments were discarded. Other types of small non-coding genes were annotated by scanning Rfam[29] against the genome and passing the results into Infernal[30].

***1.2.1.2) Results.*** *Building tissue- and development stage-specific transcript models*: A breakdown of the number of transcript structures identified across the nine tissues and three development stages, in addition to the merged sample that contained all mapped reads can be seen in Supplementary Table 11. The number of structures identified per sample ranged from 21,748 from skeletal muscle to 40,619 in the merged sample. Samples with relatively low (blood, skeletal muscle, liver) or high (brain, testis, embryo) counts of annotated transcript structures were in line with observations from other species annotated with the Ensembl gene annotation system.

*Final gene set*: Counts genes and transcripts of the finalised gene set can be seen in Extended Data Figure 1b. The gene annotation of hagfish was challenging due to the large evolutionary distance between it and other annotated vertebrates. A lot of the data available for vertebrates are biased towards mammals, with human and mouse in particular having the most well annotated genomes. As such, the annotation relied heavily on the transcriptomic data over homology-based structures wherever possible. The protein-coding gene count of 16,513 is likely to be an underestimate of the true number of coding genes and more representative of highly conserved or highly expressed genes. The annotation of hagfish was made available as part of Ensembl version 93 (July 2018).

**1.2.2) Ensembl comparative analyses (Compara team).** Hagfish (*E. burgeri*, Eburgeri_3.2) was included in the Ensembl comparative analyses from its version 93. The analyses include phylogenetic trees (on both coding and non-coding genes), orthology and paralogy relationships, as well as whole-genome alignments with some model organisms.

***1.2.2.1) Phylogenetic tree – whole genomes.*** We started with building a phylogenetic tree of all Ensembl species using the pairwise Mash[31] distances (computed on the whole, unmasked, genomes). In this analysis, hagfish is closer to the Cyclostomata ancestral node than lamprey (*Petromyzon marinus*, Pmarinus_7.0) with a distance of about 0.13 vs. 0.17.

- Download links:

https://ftp.ensembl.org/pub/release-93/compara/species_trees/ensembl_species-tree_Ensembl.nh

***1.2.2.2) Whole-genome alignments.*** The hagfish genome was aligned using LastZ[32] (version 1.04) against lamprey, zebrafish (*Danio rerio*, GRCz11), and human (*Homo sapiens*, GRCh38). For all three alignments, we used our standard parameters for vertebrates (Supplementary Table 13). In short, the pipeline[33] does an all-vs-all comparison of two genomes using LastZ, and then smoothens the alignment blocks by building *chains* and *nets*[34]. Only about 0.5% of the hagfish genome (between 12 and 13 Mb) could be aligned to either of the three species. As in other long-distance alignments, most of the alignment (49.4% of hagfish-lamprey, 59.9% of hagfish-zebrafish, 63.3% of hagfish-human) is concentrated on the coding exons, covering respectively 26.4%, 32.2%, and 31.7% of these. The hagfish alignments cover more of the zebrafish and human genomes (resp. 21.6 Mb and 22.0 Mb) than the lamprey alignments (resp. 19.3 Mb and 17.0 Mb).

- Download links:

https://ftp.ensembl.org/pub/release-93/ensembl-compara/pairwise_alignments/pmar_pmarinus_7.0.v.ebur_eburgeri_3.2.lastz_net/

https://ftp.ensembl.org/pub/release-93/ensembl-compara/pairwise_alignments/drer_grcz11.v.ebur_eburgeri_3.2.lastz_net/

https://ftp.ensembl.org/pub/release-93/ensembl-compara/pairwise_alignments/hsap_grch38.v.ebur_eburgeri_3.2.lastz_net/

***1.2.2.3) Phylogenetic trees and orthologies***. We computed phylogenetic trees on the genes from a set comprising hagfish and 100 other genomes in Ensembl, mostly vertebrates, on coding and non-coding genes, using the methods described in REFs. 33, 35. From those phylogenetic trees, orthology and paralogy relationships were derived. The first stage of the pipeline is to cluster the genes together into families, from blast-derived distances. Statistics about the resulting clusters are presented in Supplementary Table 14. In short, hagfish's non-coding gene set is too small to pursue meaningful statistics. Hagfish's coding set is about half larger than lamprey's, making the comparison of the respective statistics of both species hard. Then, for each cluster, the pipeline pulls and aligns the protein sequences, then builds a tree on the back-translated alignments with TreeBest, which itself makes a consensus of trees built with different methods and substitution models. The trees are reconciled with the species tree, and pairwise relationships (incl. orthologies) are derived. Overall, hagfish has about 7,500 orthologues with lamprey, and 11,000 with zebrafish and human, as seen in Supplementary Table 15. In all three cases, more than 5,000 of them are in a single-copy in both species. In contrast (Supplementary Table 16), lamprey has about 8,000 orthologues with zebrafish and human, about 4,000 of them being in single-copy, but lamprey's gene annotation is far smaller than hagfish's. For each pair of orthologues, we compare whether the loci were aligned by LastZ, and flag the orthologues that have sufficient LastZ coverage and sequence identity [https://jul2018.archive.ensembl.org/info/genome/compara/Ortholog_qc_manual.html] as "high confidence". Hagfish has about 4,500 high-confidence orthologues with lamprey, and 7,100 with zebrafish and human. We also systematically compare certain gene features across species, and flag genes that are split across several models, and genes that are significantly shorter or longer than their counterpart in other species. However, these comparisons are less reliable when there are few closely related species. Figures of hagfish and lamprey (Supplementary Table 17) must be interpreted with caution.

- Download links:

https://ftp.ensembl.org/pub/release-93/emf/ensembl-compara/homologies/Compara.93.protein_default.cds.fasta.gz https://ftp.ensembl.org/pub/release-93/emf/ensembl-compara/homologies/Compara.93.protein_default.nhx.emf.gz

https://ftp.ensembl.org/pub/release-93/emf/ensembl-compara/homologies/Compara.93.ncrna_default.nt.fasta.gz

https://ftp.ensembl.org/pub/release-93/emf/ensembl-compara/homologies/Compara.93.ncrna_default.nhx.emf.gz

https://ftp.ensembl.org/pub/release-93/tsv/ensembl-compara/homologies/eptatretus_burgeri/Compara.93.protein_default.homologies.tsv.gz

https://ftp.ensembl.org/pub/release-93/tsv/ensembl-compara/homologies/eptatretus_burgeri/Compara.93.ncrna_default.homologies.tsv.gz

https://ftp.ensembl.org/pub/release-93/mysql/ensembl_compara_93/gene_member_qc.txt.gz

#### 2. TIME-CALIBRATED PHYLOGENOMICS.

Whether hagfish form a clade with lampreys (Cyclostomata) or represent the sister to all other vertebrates (including lampreys) has, for a long time, depended on whether molecular or morphological evidence are considered. Morphological studies have tended to prefer Cyclostome paraphyly[36–42]. However, Heimberg et al.[43] demonstrated that this has been a consequence of the uncritical recycling of out-dated phenotype datasets and, after revision, they cannot discriminate statistically between cyclostome monophyly and paraphyly. Subsequent analyses of morphological data have recovered cyclostome monophyly[44,45] though it is not clear that they can reject cyclostome paraphyly[36,38]. Phylogenies inferred from molecular evidence have almost exclusively recovered cyclostome monophyly[43,46–51]. Further confusion arises from a paucity of genomic data from hagfish, addressed in this publication.

##### 2.1) The phylogeny of vertebrates.

The time calibrated, reconstructed phylogeny of vertebrates with outgroup rooting (main text Figure 2 and Extended Data Figure 2b) supports monophyletic cyclostomes. All nodes were recovered with a posterior probability of 1 and topology of the rest of the tree reflects our current understanding of relationships of vertebrates.

We analysed the gene duplication and loss histories of all gene families in order to test different hypotheses of the root of vertebrates. An approximately unbiased test[52] strongly rejected hagfish sister (log likelihood difference = 7947.7, AU = 0.996, multiscale bootstrap probability < 0.001). Usual bootstrap probability, Bayesian posterior probability, a Kishino-Hasegawa test[53], a Shimodaira-Hasegawa test[54], a weighted Kishino-Hasegawa test and a weighted Shimodaira-Hasegawa test all gave a probability of monophyletic cyclostomes > 0.999.

Duplication events inferred from OrthoFinder, with greater than 50% support, are strongly enriched in the vertebrate, gnathostome, lamprey and teleost stem branches. Within vertebrates, actinopterygians and cyclostomes are also inferred to have many gene duplication events present (Extended Data Figure 2b).

By computing the likelihood of the gene gain and loss histories of gene families in vertebrates, we show that they are significantly more likely under monophyletic cyclostomes, providing another line of evidence for the monophyly of Cyclostomes. This corroborates other molecular studies, and one recent morphological study[45]. We suggest that the position of hagfishes in the tree of life has now been fully resolved.

**2.2) Calibrations.**

The calibrations are described with respect to node labels in Supplementary Figure 5. New calibrations are fully justified; where we follow previously justified calibrations, we cite the source and provide a summary justification.

**Node 1: Crown Metazoa | 574-609 Ma.**

**Fossil taxon and specimen:** *Charnia masoni* (OUM ÁT.429/p) from level DRK-10 within the Drook Formation (sensu Matthews et al.[55]) at Mistaken Point, Newfoundland[56].

**Minimum justification:** *Charnia masoni* has been interpreted as a stem eumetazoan by Dunn et al.[57]. DRK-10 within the Drook Formation at Mistaken Point has been directly dated to 574.17 Ma ± 2.8 Myr through Zircon U/Pb dating 573.4 Ma ± 0.28 Myr using Pb/Pb dating[55]. However, Matthews et al.[55] demonstrate and Yang et al.[58] argue that rangeomorphs have been found in the rock record from 574 Ma.

**Soft maximum justification:** Lantian biota - this biota has extensive macrofossils but nothing that can definitively be classified as metazoans. Age from Yang et al.[58]. This age allows for the possibility of *Eoandromeda* being a crown metazoan.

**Node 2: Crown Eumetazoa | 561.1-590.8 Ma.**

**Fossil taxon and specimen:** *Auroralumina attenboroughii* (GSM 106119; British Geological Survey, Nottingham, UK) from Bed B, Bradgate Formation, Maplewell Group, Charnian Supergroup, Leicestershire, UK[59].

**Minimum justification:** *Auroralumina attenboroughii* has been shown to be a crown-group member based on tetraradial periderm with corner sulci, allying it with medusozoans[59]. It is from Bed B, Bradgate Formation, Charnian Supergroup, which has a minimum age of 563 ± 1.9 Ma, or 561.1 Ma[60]. Newer U-Pb data suggest the minimum age of the Bradgate Formation is 556.6 ± 6.4, however this large age uncertainty entirely encompasses the maximum age for the clade. For this reason, the age of 561.1 Ma, based on Wilby et al.[60], will be used.

**Soft maximum justification:** Weng'an biota[58] which may contain total group metazoans[61], but there is no convincing evidence of crown metazoans.

**Node 3: Crown Bilateria | 532-590.8 Ma.**

Our minimum constraint on crown-Bilateria follows the justification for crown-Mollusca in Benton et al.[62].

**Minimum justification:** *Aldanella janjiahensis*, or *Aldanella attleborensis*[63] is a widely accepted stem group gastropod dated to 532 Ma

**Soft maximum justification:** Weng'an biota[58] which may contain total group metazoans[61], but there is no convincing evidence of crown metazoans.

**Node 4: Crown Protostomia | 532-590.8 Ma.**

Our minimum constraint on crown-Bilateria follows the justification for crown-Mollusca in Benton et al.[62].

**Minimum justification:** *Aldanella janjiahensis*, or *Aldanella attleborensis* is a widely accepted stem group gastropod[64], the oldest record of which is dated to 532 Ma[63].

**Soft maximum justification:** Weng'an biota[58] which may contain total group metazoans[61], but there is no convincing evidence of crown metazoans.

**Node 5: Mollusca-Annelida | 532-590.8 Ma.**

Our minimum constraint on crown-Bilateria follows the justification for crown-Mollusca in Benton et al.[62].

**Minimum justification:** *Aldanella janjiahensis*, or *Aldanella attleborensis* is a widely accepted stem group gastropod[64], the oldest record of which is dated to 532 Ma[63].

**Soft maximum justification:** Weng'an biota[58] which may contain total group metazoans[61], but there is no convincing evidence of crown metazoans.

**Node 6: Myriapoda-Pancrustacea: Mandibulata | 514-543 Ma.**

Our minimum constraint on crown-Mandibulata follows the justification in Benton et al.[62] and Wolfe et al.[65].

**Minimum justification:** *Yicaris dianensis* and *Wijicaris muelleri* both have limb characteristics that place them within crown Crustacea[62,65].

**Soft maximum justification:** *Rusophycus* trace fossils are widely accepted to be made by arthropod-grade organisms with bilateral symmetry and evidence of segmented limbs (age from REFs. 62, 65, 66. They evidence the existence of arthropods, but there is no evidence of mandibulates at this time.

**Node 7: Crown Deuterostomia | 517.32-590.8 Ma.**

Our minimum constraint on crown-Deuterostomia follows the justification in Benton et al.[62].

**Minimum justification:** *Haikouichthys ercaicunensis* from the Chengjiang Biota have been identified as a crown chordate on the basis of branchial structures, myomeres, and a notochord[67]. The age of the Chengjiang Biota was established by Yang et al.[68].

**Soft maximum justification:** Weng'an biota[58] which may contain total group metazoans[61], but there is no convincing evidence of crown metazoans.

**Node 8: Crown Ambulacraria | 515.15-590.8 Ma.**

Our minimum constraint on crown-Ambulacraria follows the justification in Benton et al.[62].

**Minimum justification:** Isolated pelmatozoan columnals that have characteristic morphology and stereom structure of the echinoderm total group[69,70].

**Soft maximum justification:** Weng'an biota[58] which may contain total group metazoans[61], but there is no convincing evidence of crown metazoans.

**Node 9: crown-Chordata | 517.32-590.8 Ma.**

Our minimum constraint on crown-Chordata follows the justification in Benton et al.[62].

**Minimum justification:** *Haikouichthys ercaicunensis* from the Chengjiang Biota have been identified as a crown chordate on the basis of branchial structures, myomeres, and a notochord[67]. The age of the Chengjiang Biota was established by Yang et al.[68].

**Soft maximum justification:** Weng'an biota[58] which may contain total group metazoans[61], but there is no convincing evidence of crown metazoans.

**Node 10: crown-Olfactores | 517.32-590.8 Ma.**

Our minimum constraint on crown-Olfactores follows the justification in Benton et al.[62].

**Minimum justification:** *Haikouichthys ercaicunensis* from the Chengjiang Biota have been identified as a crown chordate on the basis of branchial structures, myomeres, and a notochord[67]. The age of the Chengjiang Biota was established by Yang et al.[68].

**Soft maximum justification:** Weng'an biota[58] which may contain total group metazoans[61], but there is no convincing evidence of crown metazoans.

**Node 11: crown-Vertebrates | 497-590.8 Ma.**

**Fossil taxon and specimen**: *Furnishina bigeminata* (GMPKU2589; Geological Museum of Peking University, Beijing) from the middle Cambrian Huaqiao Formation, Wangcun section[71].

**Minimum justification:** Conodonts are tooth-like 'elements' that have been interpreted as coming from vertebrates, possibly stem gnathostomes[72]. Soft tissue fossils show notochords, segmental trunk musculature, a caudal fin, paired sensory organs, and gill pouches, all of which demonstrate a vertebrate affinity. The oldest conodont is a *Furnishina*, a paraconodont in the euconodont lineage[73] and the earliest records of *Furnishina* are from the middle Cambrian Huaqiao Formation at Wangcun section, dated to 497 Ma[71].

**Soft maximum justification:** Weng'an biota[58] which may contain total group metazoans[61], but there is no convincing evidence of crown metazoans.

**Node 12: crown-Gnathostomata | 420.7-468.4 Ma.**

Our minimum constraint on crown-Gnathostomata follows the justification in Benton et al.[62].

**Minimum justification:** *Guiyu oneiros* has lobe-finned fish synapomorphies which identify it as a stem sarcopterygian and therefore crown Osteichthyes and crown Gnathostomata[62,74].

**Soft maximum justification:** Stem group gnathostomes ('ostracoderms') are present in the Ordovician, but there are no crown representatives in any of the well preserved fossil localities. This suggests the crown group was not yet present[62].

**Node 13: crown-Osteichthyes | 420.7-444.9 Ma.**

Our minimum constraint on crown-Osteichthyes follows the justification in Benton et al.[62].

**Minimum justification:** *Guiyu oneiros* has lobe-finned fish synapomorphies which identify it as a stem sarcopterygian and therefore crown Osteichthyes and crown Gnathostomata[62,74].

**Soft maximum justification:** Early Silurian rocks have diverse jawless fishes, but there is nothing that is clearly crown-osteichthyan. The Llandovery can therefore provide the maximum age[62].

**Node 14: crown-Sarcopterygii | 408-427.9 Ma.**

Our minimum constraint on crown-Sarcopterygii follows the justification in Benton et al.[62], which was based on *Youngolepis* sp. from the lower Xishacun Formation, Quijing, Yunnan, China (IVPP V10519.1, Institute of Vertebrate Palaeontology and Palaeoanthropology, Beijing). We defer to Benton et al.[62] for the phylogenetic and age justification of the soft maximum constraint.

**Node 15: crown-Rhipidistia | 408-427.9 Ma.**

Our minimum constraint on crown-Rhipidistia follows the justification in Benton et al.[62], which was based on *Youngolepis* sp. from the lower Xishacun Formation, Quijing, Yunnan, China (IVPP V10519.1, Insitute of Vertebrate Palaeontology and Palaeoanthropology, Beijing). We defer to Benton et al.[62] for the phylogenetic and age justification of the soft maximum constraint.

**Node 16: crown Tetrapoda: 337-351 Ma.**

Our minimum constraint on crown-Tetrapoda follows the justification in Benton et al.[62].

**Minimum justification:** *Lethiscus stocki* is either a Lepospondyli (within Batrachomorpha) or a Reptiliomorpha, but since both are crown Tetrapoda it can be safely interpreted as a crown tetrapod[62].

**Soft maximum justification:** Although both baphetids and colosteids are nearer the crown of Tetrapoda, both are found in much younger deposits. Whatcheerids are more distantly related, but are found in older deposits, so these are used as the maximum age constraint - crown tetrapods are not found alongside them[62].

**Node 17: crown Amniota: 318-332 Ma[62].**

Our minimum constraint on crown-Amniota follows the justification in Benton et al.[62].

**Minimum justification:** The eureptilian *Hylonomus lyelli* is a stem diapsid, making it a crown amniote[62].

**Soft maximum justification:** The East Kirkton locality has diverse batrachomorphs and reptiliomorphs, but no diapsids or synaposids, and older sites show the same thing[62].

**Node 18: crown-Reptilia: 255.9-295.9 Ma.**

Our minimum constraint on crown-Reptilia follows the justification in Benton et al.[62] which is based on *Protorosaurus spenceri* Meyer, 1832 (Royal College of Surgeons, RCSHC/Fossil Reptiles 308), holotype of the oldest archosauromorph, from the Kupferschiefer of Germany and the Marl Slate of NE England[75]. We defer to Benton et al.[62] for the phylogenetic and age justification of the soft maximum constraint.

**Node 19: crown-Archelosauria | 250.6-295.9 Ma.**

Our minimum constraint on crown-Reptilia follows the justification in Benton et al.[62] which is based on *Proterochersis robusta* which is the oldest unequivocal member of total-group Testudines. Though its phylogenetic placement within crown or stem[76,77] Testudines is contested, its membership of total-group Testudines is not. We defer to Benton et al.[62] for the phylogenetic and age justification of the soft maximum constraint.

**Node 20: crown Neognathae | 66.0-86.8 Ma.**

Our minimum constraint on crown-Neognathae follows the justification in Benton et al.[62] which is based on *Vegavis iaai* (holotype, Museo de la Plata, MLP 93-I-3-1), a partial skeleton from lithostratigraphic unit K3 of Vega Island, Antarctica. We defer to Benton et al.[62] for the phylogenetic and age justification of the soft maximum constraint.

**Node 22: crown-Mammalia | 176.03-252.2 Ma.**

**Fossil taxon and specimens:** *Henosferus molus* (MPEF 2353; Museo Paleontológico Egidio Feruglio, Trelew, Patagonia, Argentina) from strata belonging to the lowermost Cañadón Asfalto Formation (or beds transitional with the underlying Lonco Trapial Formation) at Queso Rallado[78,79].

**Minimum justification:** Rougier et al.(2007) identify *Henosferus molus* as an early diverging australosphenidan (pan-monotreme) based on a formal morphology-based phylogenetic analysis.

**Age justification**: Cúneo et al.[78] established a minimum age for the Cañadón Asfalto Formation of 176.15 ± 0.12 Ma, thus 176.03 Ma.

**Soft maximum justification:** Some stem-monotremes may extend into the Triassic, and to include the possibility of the haramiyids being crown mammals the Permian-Triassic boundary can be used as the soft maximum[80].

**Node 23: crown-Theria | 121.56-169.6 Ma.**

**Node Calibrated:** The clade comprised of Marsupialia and Placentalia.

**Fossil Taxon and Specimens:** *Sinodelphys szalayi* (Chinese Academy of Geological Sciences CAGS00-IG03, articulated holotype specimen).

**Phylogenetic Justification:** *Sinodelphys szalayi* was resolved on stem to Marsupialia, within Theria, by Luo et al.[81,82], Yuan et al.[83] and Bi et al.[84].

**Age Justification:** Fossils from the Jehol biota of northeast China come from Barremian to Aptian-age deposits, although there is some ambiguity about their age[85]. Here we take the youngest age estimate from tuffs overlying the main fossil-bearing layers, dated to 121.96 Ma ± 0.5 Myr[86], thus 121.56 Ma. We conservatively use the age of the Barremian to define the ages of Jehol specimens, which has an upper margin of 125.0 Ma. Given the possibility that southern, tribosphenic mammals such as *Ambondro* are therian (cf. REF. 87), we set the soft maximum age for Theria in the Bathonian, 168.3 Ma ± 1.3 Myr so 169.6 Ma[86].

**Node 24: crown-Placentalia | 61.6-164.6 Ma.**

Our minimum constraint on crown-Placentalia follows the justification in Benton et al.[62] which is based on the carnivoran *Ravenictis krausei* (UALVP 31175) from the Ravenscrag Formation, Saskatchewan[88,89]. We defer to Benton et al.[62] for the phylogenetic and age justification of the soft maximum constraint.

**Node 25: crown-Boreoeutheria | 61.6-164.6 Ma.**

Our minimum constraint on crown-Boreoeutheria follows the justification in Benton et al.[62] which is based on the carnivoran *Ravenictis krausei* (UALVP 31175) from the Ravenscrag Formation, Saskatchewan[88,89]. We defer to Benton et al.[62] for the phylogenetic and age justification of the soft maximum constraint.

**Node 26: crown-Euarchontoglires | 61.6-164.6 Ma.**

Our minimum constraint on crown-Euarchontoglires follows the justification in Benton et al.[62] which is based on Torrejonian occurences of extinct primate sister taxa such as plesiadapids from north-eastern Montana, e.g., *Paromomys farrandi* (UCMP 189520)[90]. We defer to Benton et al.[62] for the phylogenetic and age justification of the soft maximum constraint.

**Node 27: crown-Atlantogenata | 56.0-164.6 Ma.**

Our minimum constraint on crown-Atlantogenata follows the justification in Benton et al.[62] which is based on the extinct proboscidean *Eritherium azzouzorum* (MNHN PM69, Paris) from the Sidi Chennane quarries, phosphate bed IIa, lower bone-bed, Ouled Abdoun Basin of Morocco, regarded as upper Paleocene (early Thanetian)[91]. We defer to Benton et al.[62] for the phylogenetic and age justification of the soft maximum constraint.

**Node 28: crown-Marsupialia | 47.6-131.3 Ma.**

Our minimum constraint on crown-Marsupialia follows the justification in Benton et al.[62] which is based on *Djarthia murgonensis* (Queensland Museum, QM F52748)[92], from the early Eocene of Murgon, Australia. Phylogenetic Justification. *Djarthia* possesses a continuous lower ankle joint, a synapomorphy shared with the marsupial clade Australidelphia[92]. The status of Djarthia as a crown marsupial is stronger than that for Khasia[92,93], and thus it serves as a more definitive record of the latest point by which crown marsupials evolved. We defer to Benton et al.[62] for the phylogenetic and age justification of the soft maximum constraint.

**Node 29: crown-Pipoidea | 148.1-201.5 Ma.**

Our minimum constraint on crown-Pipoidea follows that of Cannatella[94] which we reiterate here:

**Node Calibrated:** The divergence between *Xenopus-Nanorana* which equates to crown-Pipoidea[94].

**Fossil Taxon and Specimens:** *Rhadinosteus parvus* (holotype DINO 14693, Dinosaur National Monument, Colorado, USA, partial skeleton, articulated to loosely articulated).

**Phylogenetic Justification:** *Rhadinosteus parvus* is the fossil representative of crown-Pipoidea, identified as a stem-rhinophrynid, more closely related to *Rhinophrynus* than Pipidae[95,96].

**Age Justification:** *Rhadinosteus parvus* was recovered from the Brushy Basin Member of the Morrison Formation of Utah[96] which has been dated to 148.1-150.3 Ma[97], thus our minimum constraint on crown-Pipoidea is 148.1 Ma. Our soft maximum constraint on the *Xenopus-Nanorana* divergence follows that for crown-Anura in Benton et al.[62] which was based on the Triassic-Jurassic boundary, allowing for finds of stem anurans and other small tetrapods from several continents, but as yet no crown representatives of these clades. The Triassic-Jurassic boundary is dated to 201.3 Ma ± 0.2 Myr[86], and so 201.5 Ma.

**Node 30: crown-Neopterygii | 250.0-331.1 Ma.**

Our minimum constraint on crown-Neopterygii follows the justification in Benton et al.[62], which was based on *Watsonulus eugnathoides*[98] from the Middle Sakamena Formation, Sakamena Group, Ambilombe Bay, Madagascar (syntype MNHN MAE 33a, b, Muséum national d'Histoire naturelle, Paris). We defer to Benton et al.[62] for the phylogenetic and age justification of the soft maximum constraint.

**Node 31: crown-Clupeocephala | 150.94-235 Ma.**

Our minimum constraint on crown-Clupeocephala follows the justification in Benton et al.[62] which is based on *Leptolepides haerteisi(Arratia 1997)* from the Solnhofen Formation, Zandt Member, Zandt, Bavaria, Germany (holotype JM SOS 2473, Jura Museum, Eichstätt, Germany). We defer to Benton et al.[62] for the phylogenetic and age justification of the soft maximum constraint.

**Node 32: Ovalentaria-Tetraodontiformes | 69.71-130.8 Ma.**

Our minimum constraint on the divergence between Ovalentaria and Tetraodontiformes follows the justification in Benton et al.[62] which is based on *Cretatriacanthus guidottii* from the 'Calcari Melissano' (of historical usage) at Canale near Nardò, Italy (holotype MCSNV 1377, Museo Civico di Storia Naturale, Verona, Italy). We defer to Benton et al.[62] for the phylogenetic and age justification of the soft maximum constraint.

**Node 33: crown-Chondrichthyes | 333.56-422.4 Ma.**

Our minimum constraint on crown-Chondrichthyes follows the justification in Benton et al.[62], which was based on *Chondrenchelys problematicus* from Mumbie Quarry, Glencartholm Volcanic Group, Upper Border Group of the Calcifererous Sandstone, Glencartholm, Scotland (NMS

1998.35.1, National. We defer to Benton et al.[62] for the phylogenetic and age justification. We defer to Benton et al.[62] for the phylogenetic and age justification of the soft maximum constraint.

**Node 36: crown-Cyclostomata | 358.5-614 Ma.**

Our minimum constraint on crown-Cyclostomata follows the justification in Benton et al.[62], which was based on *Priscomyzon riniensis* (Albany Museum, Grahamstown, Eastern Cape, South Africa, catalogue number AM5750), holotype, consisting of a whole dorso-ventrally compressed organism. We defer to Benton et al.[62] for the phylogenetic and age justification. Our soft maximum constraint is based on the Ediacaran Weng'an Biota which preserves a diversity of multicellular eukaryotes preserved in subcellular fidelity, but which has yielded no evidence of chordates[99]. The Weng'an Biota has been dated to 609 Ma ± 5 Myr[100], hence we use it to evidence a 614 Ma soft maximum constraint on the age of crown-Chordata.

### 3. GENE FAMILY EVOLUTION

#### 3.1) Gene family gains and losses across vertebrate evolution.

We used comparative genomic analyses (see Methods) to infer the gene complements of different ancestors within deuterostomes. The ancestral vertebrate genome was composed of at least 9952 gene families, from which ~17,000 modern human genes descend. The most abundant biological functions in the ancestral vertebrate genome are related to metabolite interconversion, regulation of transcription, and transmembrane proteins involved in signal reception and transport. The most predominant pathways are related to Wnt, inflammation, gonadotropin-releasing hormones, integrins, and angiogenesis (Supplementary Table 23).

Gnathostomata and Vertebrata are among the nodes with the largest amount of gene gains and lowest number of losses (respectively Novel Homology Groups (HGs) and Lost HGs, Figure 2, Extended Data Figure 3a and Supplementary Table 24), supporting a major role for genomic novelty and conservation in the origins of those lineages (see Novel Core HG below). On the other hand, Cyclostomata and Olfactores are among the clades showing the largest number of gene losses (Figure 2, Extended Data Figure 3a). The gene gains and loss values observed here are similar to the numbers observed in other major metazoan transitions in previous studies[101,102].

The evolution of gene novelties and their functions across the different nodes (Supplementary Table 24) shows that the Novel HG of the LCA of vertebrates are dominated by genes involved in signalling, cell adhesion, and regulation of transcription (e.g., ‘gene-specific transcriptional regulator’, ‘nucleic acid binding protein’). The Novel HGs of the LCA of gnathostomes display many novel genes related to the immune system (Supplementary Table 24).

An interesting subset of novel genes is those highly retained after their emergence (Novel Core HG), as they may contain genes of high functional relevance. Cyclostomata, Vertebrata, and Gnathostomata show the highest numbers of conserved gene novelties (Novel Core HG, Figure 2, Extended Data Figure 3a). This is consistent with the patterns of WGD described elsewhere in this study, and hints at the importance of novelty over loss in these three clades. These numbers here are indeed larger than the equivalent values for other metazoan transitions; for example, here Cyclostomata, Vertebrata, and Gnathostomata respectively have ++ 98, ++ 86, and ++ 81 Novel Core HG, compared to ++ 25 in Metazoa, ++ 0 in Bilateria, ++ 1 / ++ 5 in the three bilaterian superclades[101,102].

The list of highly retained new HG (Novel Core HG) shows 5 Novel Core HG in Chordates, including the ZHX homeobox genes family, a keratin gene (KRT), or MARCH7 which is an ubiquitin involved in immune response. Vertebrates retained many novel genes involved in signalling like the Novel HG (Supplementary Table 25, Extended Data Figure 3b); thus, signalling genes not only were predominant among the novel genes, but they were also highly retained later across all vertebrate lineages. Similarly, gnathostome Novel Core HG (Supp show that over 71% of conserved novel genes are immunity genes, also seen among the Novel HG (Supplementary Table 26, Extended Data Figure 3b). In contrast to vertebrates, gnathostomes retained genes involved in gene regulation (gene-specific transcriptional regulators, nucleic acid binding protein) compared to their Novel HGs.

**3.2) Analysis of hagfish gene families of interest.**

**3.2.1) Immune system related genes.** Due to their pivotal phylogenetic position, the agnathans serve as model organisms for understanding the origin and early evolution of the vertebrate adaptive immune system[103]. Both hagfish and lamprey possess B-like and T-like lymphocytes, but unlike the jawed vertebrates which use immunoglobulin domain-based anticipatory receptors, they use leucine-rich repeat (LRR)-based variable lymphocyte receptors (VLRA, VLRB and VLRC)[103]. We carried out similarity searches against the *E. burgeri* genome using available assembled hagfish *VLR* cDNAs. In the *E. burgeri* genome sequence, the majority of *VLR* donor cassettes and three germline *VLR* genes are found on three large scaffolds (Supplementary Figure 6). The *VLRB* and *VLRC* germline genes are located next to 94 and 19 donor cassettes, respectively, on assembly cluster15 (Supplementary Figure 6a), while the *VLRA* germline gene flanks 14 donor cassettes on cluster12 (Supplementary Figure 6b). Two additional clusters containing 16 and 36 *VLRA*-like donor *LRRV* cassettes are separated by 4.6 Mbp on cluster1 (Supplementary Figure 6b). Shared *LRR* cassette usage for *VLRA* and *VLRC*[104] suggests that a conserved functional relationship could exist between the VLRA- and VLRC-producing T-cell lineages in hagfish (Supplementary Figure 6c). The analysis of the hagfish genome assembly reveals that it harbours gene homologues of important genes that function in adaptive and innate immunity in jawed vertebrates, although orthology is complicated by the divergent genomic history of the agnathans (e.g., *GATA2/3* and *TLR13/21*) (Supplementary Table 27). Our results indicate that despite the differences in the anticipatory receptors in jawed and jawless vertebrates, several fundamental components of lymphocyte development and regulation were present in the common ancestor of living vertebrates.

***Methodology***: Variable lymphocyte receptor BLAST searches of the *E. burgeri* genome sequence assembly were carried out using sequence fragments taken from available hagfish *VLR* cDNA sequence. Accession numbers are as follows: *E. burgeri VLRC* (initially labeled as VLRA): AY964719.1-AY964729.1, AY964731.1-AY964762.1, AY964764.1-AY964782.1,

AY964786.1-AY964789.1; *E. burgeri VLRB*: AB519991.1-AY965525.1, AY965527.1-AY965534.1; *E. stoutii VLRA* (third VLR): KF314046.1-KF314109.1; *E. stoutii VLRC*: AY964837.1, AY964783.1-AY964785.1, AY964790.1-AY964809.1, AY964811.1-AY964822.1,AY964825.1-AY964836.1, AY964838.1-AY964872.1, AY964874.1-AY964881.1, AY964883.1-AY964906.1, AY964908.1-AY964918.1 and AY964920.1-AY964931.1. Fragments used in the searches were as follows: LRRNT, LRR1, LRRV, LRRVe-CP, LRRCT, germline signal peptide and germline C-terminal regions. Both BLASTN and TBLASTN (using translated fragment sequences) were carried out using expect values of e ≤ 10-6, although higher expect values were sometimes used to detect short sequence matches to connecting peptide (CP) and LRR1 sequences. All matches were individually inspected.

**3.2.2) Hagfish vision related genes.** The vertebrate visual pathway includes a diversity of proteins conserved across the entire clade. However, how this pathway was assembled in Deuterostomia remains unclear, and multiple lineage-specific gene-losses across Vertebrata confuse its subsequent evolution. While lamprey genomes have been available for a while, the absence of a hagfish genome, together with some uncertainty on the evolutionary relationships between the hagfishes –with their simpler and perhaps secondarily degenerated eyes[105]—, the lampreys and the gnathostomes, hampered progress in our understanding of the evolution of vision at the root of the vertebrate tree.

We investigated patterns of presence and absence in 49 vision-associated proteins across 39 deuterostomes, including the hagfish *E. burgeri*. Our results show that while a subsample of these genes (average = 13.2) is shared across all deuterostomes, cyclostomes have approximately twice as many visual genes (Average = 30.5), and *E. burgeri* has 27. The average number of genes associated with vision further increase in the gnathostomes, with an average of 42.5 visual genes.

Three visual pigments (Opsins) were found in the genome of *E. burgeri*: Melanopsin (Opn4), Peropsin (RRH) and Rhodopsin 1 (RH1 or RHO) (Supplementary Figure 7). In vertebrates, Melanopsin is expressed in ipRGCs cells or their homologues[105] and this opsin is not involved in processes of image-forming vision. Differently, RH1 is known to be expressed in rod cells and is generally involved in dim-light (scotopic) image forming vision[105–107]. Peropsin[108,109] is also not directly involved in the process of image-forming vision. However, Peropsin plays a key role (together with the RGR Opsin – that we did not detect in the hagfish) in re-isomerising (recycling) *all-trans* retinal to *11-cis* retinal[110]. In the process of vertebrate vision, *11-cis* retinal, which is attached to opsins, like the RH1 opsin, through the retinal binding domain, transforms into *all-trans* retinal after being hit by a photon of light. *All-trans* retinal is then detached from its opsin (a process known as opsin bleaching). *All-trans* retinal is re-isomerised to *11-cis* retinal and reattached to visual opsins so that they can be re-used in the process of image-forming vision. Peropsin (RRH), that we found in the hagfish genome, is one of the key proteins in the process of retinal re-isomerisation[110], and its presence in the hagfish genome is evidence that RH1 is functional in the hagfish. None of the other vertebrate opsins used for image-forming vision (LWS, SWS1, SWS2 and RH2) are found in the genome of *E. burgeri*. However, these genes have previously been identified in lampreys, with the pouched lamprey *Geotria australis*, for example, possessing the full complement of vertebrate visual pigments[107]. These observations suggest that cones and rods, as well as cone-mediated bright-light colour vision and rod-mediated dim-light vision, most likely emerged in the stem vertebrate lineage[107], and that *E. burgeri* secondarily lost all its visual opsins with the exclusion of RH1. The hagfish eye has been suggested to be degenerate, and its retina only seems to include one type of photoreceptor cell. While the morphology of these photoreceptors cannot discriminate whether the hagfish retina is composed of modified cones or rods, Lamb[105] has pointed out that "hagfish photoreceptors exhibit some rod-like properties". Our results agree with Lamb's assertion.

The emergence of both cones and rods in the stem vertebrate lineage is confirmed by the analysis of visual pathway genes (Supplementary Figure 7). Six of the 11 candidate genes that are known to be involved in both the cone and rod opsin pathways (PDC, GPSM2, GNB5, GUCY2F, RGS9 and RGS9BP) were present in the last common ancestor of all Deuterostomia, and predate the origin of cones, rods and of the cone- and rod-specific opsins. The remaining 5 (RCVRN, GRK1, GUCA1B, GUCY2D, and GUCA1A) are present in both Cyclostomata and Gnathostomata, but are not found outside Vertebrata; suggesting that these genes emerged in the stem vertebrate lineage. In Cyclostomata, *E. burgeri* secondarily lost GUCA1A and GUCY2D, while the brook lamprey (*Lampetra planeri*) lacks GNB5, RGS9 and RGS9BP. *L. planeri* and *E. burgeri* also shared the absence of RCVRN and GRK1, while *L. planeri* and a second lamprey (Arctic lamprey, *Lethenteron camtschaticum*) miss GUCA1A (see Supplementary Figure 7).

Ten genes that are exclusively expressed in cones were considered. Three of these genes (CNGB3, SLC24A2, CNGA3) predate the emergence of the cone opsins, being present across all Deuterostomia. Six genes (ARR3, GRK7, GNAT2, GNGT2, PDE6H and PDE6C) emerge in the stem vertebrate lineage, while GNB3 is gnathostome-specific despite having been lost in some Chondrichthyes (see Supplementary Figure 7). Within Cyclostomata, GNAT2, GNGT2 and PDE6H were each only present in one of the studied species: *L. planeri*, *L. camtschaticum* and the sea lamprey *Petromyzon marinus* respectively. The genomes of the brook and Arctic lampreys also lack PDE6C and CNGB3. While the hagfish has the three genes used by cones that are found across all deuterostomes, only two of the vertebrate-specific cone-expressed genes are present

(ARR3 and GRK7). This pattern of losses suggests a loss of cone-associated visual processes in the hagfish (see Supplementary Figure 7).

All rod-opsin pathway specific genes are present in at least one cyclostome, suggesting that this pathway was present in their last common ancestor. As these genes are also broadly distributed across gnathostomes, we can conclude that the rod-opsin pathway also evolved in the stem vertebrate lineage (as suggested by the presence/absence of opsin genes). Three rod-expressed genes CNGB1, CNGA1 and SLC24A1 are broadly distributed in Deuterostomia, indicating a longer evolutionary history, similar to their cone-expressed paralogs (CNGB3, CNGA3, SLC24A2), and a broader, non-rod specific, functional role. Five rod-specific genes (GNGT1, PDE6B, PDA6A, CNGB1 and CNGA1) could not be found in *E. burgeri.* As these genes are broadly distributed across Deuterostomia, it is possible that some of these genes might not have been sequenced or they might have been incorrectly annotated. GNGT1, on the other hand, is absent from all cyclostomes and seems gnathostome-specific (see Supplementary Figure 7).

The level of conservation of genes involved in the visual cascade seems much more variable. Six of the 12 genes we considered are present across Deuterostomia (RDH8, RDH12, RPE65, RGR, RDH10 and RDH11). However, RDH11 appears to have been independently lost within Hemichordata and Cephalochordata, while RDH8 appears to have been independently lost in Cephalochordata and Urochordata. RBP3 is present in the amphioxus *Branchiostoma belcheri* but could not be found in any other Cephalochordate or Urochordate considered. If the presence of this gene (which is broadly distributed across vertebrates) in *B. belcheri* is not the result of some form of contamination, it would suggest that RBP3 emerged in the stem chordate lineage and was independently lost multiple times in Cephalochordata and Urochordata. Differently, RLBP1 appears to be specific to the Olfactores (tunicates and vertebrates); LRAT, RBP1 and ABCA4 emerge as vertebrate-specific and RDH4 is Gnathostome-specific. Within Cyclostomata, *E. burgeri* and *L. plnaeri* share the absence of RP3 and ABCA4, whilst *L. planeri* and *L. camtschaticum* both lack RDH8. Further independent absences are also observable, such as that of RBP1 in *L. planeri*, and of RDH11 and RGR in *E. burgeri*, respectively.

Given the monophyly of Cyclostomata, our results suggest that the cone and rod opsin specific pathways were established in the stem vertebrate lineage, and that the last common vertebrate ancestor was potentially capable of both bright-light colour vision and dim-light vision. The last common ancestor of the cyclostomes possessed all known visual opsins[107], as well as cone- and rod-specific opsin pathways genes. Hagfishes have eyes with a simplified retina and photoreceptors of unknown type. The presence of only one visual pigment (the rod-specific RH1) in *E. burgeri* and the loss of most cone-specific genes in this taxon, as well as several rod-specific genes, indicates that the hagfish have secondarily simplified eyes, that they lost the ability to see in colour, and that their photoreceptors may be modified rods.

All visual-related genes found in this analysis are provided as a multifasta file in the Supplementary File 2.

***Methods.*** 53 vision-associated and photosensitive proteins were selected based on Emerling and Springer[111], with the addition of the opsin sequences Peropsin and Melanopsin. These genes include those involved in both the cone and rod phototransduction pathways, alongside visual pathway genes and genes associated with visual disorders, and genes involved in light detection that are not directly used for image forming vision (Melanopsin and Peropsin). These genes have previously been used to establish secondary adaptation to low photic environments in mammals[111].

Independent BLAST searches for each of the 53 candidate genes (see Supplementary Figure 7 for a list) were conducted across 39 high coverage genomes selected from across Deuterostomia. Significant hits (e-value < e-20) were subjected to a BLAST search against the nr database in NCBI to confirm identity. Sequences that were positively confirmed as belonging to the candidate gene family were marked as present in Supplementary Figure 7. Those that could not be positively confirmed after these two steps were marked as absent.

**3.2.3) Circadian clock genes.** Cyclostomes are representatives of an ancient vertebrate lineage. The study of the circadian clock and of the non-visual photoreception in this taxon is useful to understand the evolution of the circadian timekeeping system and of photic entrainment (synchronization) in vertebrates.

In gnathostomes, the suprachiasmatic nucleus (SCN) of the hypothalamus, the retina and the pineal gland have been reported to be important sites of circadian oscillations. Hagfishes show daily rhythms of locomotor activity in light-dark cycles with a marked nocturnal pattern[112,113]. The rhythm is maintained in constant darkness demonstrating the existence of a functional circadian clock. Interestingly, the hagfish lack of a pineal gland and experiments of ablation showed that the central pacemaker generating behavioural rhythmicity in hagfish is located in the preoptic nuclei (PON) of the hypothalamus[114]. The light signal is perceived by the retina of the eyes and sent to the PON via the pretectum[115]. Hagfish eyes do not allow an image-forming vision, but they are able to sense light and drive behavioural responses, such as shadow detection and photophobic response. A previous investigation showed the expression of Melanopsin in the ganglion cells of the retina, which indicate that these cells could mediate the photic entrainment[116]. Furthermore, as in jawed vertebrates, extraocular photoreception has been demonstrated in the hagfish: it has a direct behavioural reaction to external light stimuli via the photoreceptor of the skin[117].

Interestingly, the photic signal through skin does not entrain behavioural circadian rhythms. However, it would be expected that light also penetrates deep into the brain and tissues of hagfishes, thereby participating in photic entrainment. Until now the molecular circadian clock of hagfish has not been characterised.

We surveyed the genome of *E. burgheri* for clock genes and clock-controlled genes, and found a primordial vertebrate circadian system, with fewer orthologous genes, in respect to gnathostomes, that presumably cover all basic functions. In particular, we found the following *bona fide* clock genes: 2 Clock, 3 Bmal, 1 Timeless (Tim), 2 Period (Per), 1 Cryptochrome (Cry), Nfil3-6 (E4bp4) and Tef (Supplementary Table 28). Interestingly, the opsin repertoire in hagfish (see previous section) is constituted exclusively by a Rhodopsin (RH1 or RHO), the light-sensitive receptor protein involved in phototransduction, a Peropsin (RRH) that recycle *all-trans* retinal to *11-cis* retinal, and a Melanopsin (Opsin 4), necessary for the photic entrainment. It would be very interesting to study in the future their expression profile in the retina, brain, gonads and skin.

Genomic and transcriptomic searches failed, nevertheless, in finding the Melatonin synthesis (Aanat) and Melatonin receptor (MtnR) genes. This is probably correlated with the documented anatomical absence of the pineal gland in hagfish. Conversely, lampreys have the pineal gland, which plays a central role in the generation of circadian locomotor rhythms[118]. We also failed to identify genes involved in the DNA repair, such as 6-4 photolyase (Cry5), DASH and CPD photolyase (Cpdp). As previously shown in cavefish, *Phreatichthys andruzzii*[119], the absence of these genes could be correlated with the adaptation to extreme photic environments.

#### 4. MACROSYNTENY ANALYSES OF VERTEBRATE GENOMES

##### 4.1) Source of genomic data.

All genome assemblies and protein annotations used for these analyses are provided in Supplementary Table 1.

##### 4.2 ) Defining homologous genes.

We conducted phylogeny-aware BLASTP genome-wide searches to identify between-species homologs. That is, for analysis between chicken, gar and elephant shark, or between hagfish and sea lamprey, or between sea cucumber and lancelet, we used reciprocal best hit gene pairs as homologs (Supplementary File 3). By contrast, for the alignment between vertebrate species (chicken, gar, elephant shark, lamprey and hagfish) and non-vertebrate species (sea cucumber, amphioxus), we used genes of later species as reference and searched for the one-way best hit gene from the former species as homologs (Supplementary File 3). In this way, we identified multiple homologs in vertebrates corresponding to one gene in non-vertebrate species, which is free from WGD. Since we reconstructed the gene content of 16 ACs by taking amphioxus genes as surrogates, the homology relationship between vertebrate species and ACs is directly proxied from that between vertebrate species and amphioxus. For chicken or gar, we also performed all against all BLASTP searches to identify reciprocal interchromosomal best hit pairs as within-species homologs. That is, BLASTP matched pairs from the same chromosomes or scaffolds were excluded to control for small scale duplication events.

#### 4.3) Definition of homologous chromosomes.

Given the gene pair level between- and within-species homology relationships, we tested whether two chromosomes harbour an excess of homologous pairs via Chi-Square test. Specifically, we counted that there are m genes on a single reference chromosome B. There are in total M genes on all reference chromosomes. We then identified n homologous genes between query chromosome A and reference chromosome B. And there are in total N homologous genes harboured by chromosome A relative to all reference chromosomes. We used these four numbers (n, N-n, m, M-m) to conduct the Chi-Square test followed by FDR correction. Homologous gene pair numbers over 5 and FDR q value less than 0.05 are used as the cut-off. This strategy is used across the whole manuscript.

#### 4.4) Macrosynteny between chicken and spotted gar.

Although chicken chromosomes and gar LGs largely form a one-to-one relationship in terms of distribution of homologous pairs (Supplementary Figure 10), there are a few one-to-many or many-to-one relationships, suggesting chicken- or gar-specific chromosomal fusion or fission events. We polarised such events by taking the shark genome as the outgroup. For example, gar LG5 is simultaneously homologous to chicken chr6 and chr12. We examined whether chr6 and chr12 of chicken correspond to different sets of shark scaffolds. If true, one gar-specific fusion would be inferred. Otherwise, one chicken-specific fission event would be more plausible. To increase the certainty, we required at least two shark scaffolds to support one scenario. As a result, we identified three gar-specific fissions (LG9 and LG11, LG1 and LG16, LG3, LG17 and LG14), ten gar-specific fusions (LG1, LG3, LG4, LG5, LG6, LG7, LG10, LG12, LG14, and LG26), three chicken-specific fusions (chr1, 4 and 5). Note, chicken chr1 seems to arise from a fusion of four ancestral chromosomes, while only two chromosomes were involved in emergence of chr4 or chr5. In addition, we identified one ambiguous case where both gar LG2 and LG4 are co-homologous to chicken chr4 and Z, while shark scaffolds support both the linkage of gar LG2 and LG4, and the linkage of chicken chr4 and Z. That is, all three species differ in their karyotype, and we thus left this case unclarified.

After polarization, we artificially back-mutated chicken and gar chromosomes to approximate the ancestral karyotype. In other words, for chicken- or gar-specific fusions, we manually split these chromosomes. The boundary was inferred based on the correspondence between chicken chromosomes and spotted gar LGs. If necessary, shark scaffolds were also used to confirm. Taking chicken chr5 as an example, genes from 1.0 Mb to 24.2 Mb (chr5:1.0-24.2 Mb) are defined as chr5.1, which corresponds to gar LG27 (Supplementary Figure 11). The remaining two parts of chr5:0–1.0 Mb and chr5:24.2 – 59.6 Mb are homologous to gar LG7 and to the same set of shark scaffolds (Supplementary Figure 11). Thus, we grouped them as chr5.2. The situation can be more complex. Taking chicken chr1 subjected to three fusions as an example, genes from chr1:0 Mb-75.7 Mb but excluding a middle region chr1:48.0-52.2 Mb are defined as chr1.1. We grouped chr1:0-48.0 Mb and chr1:52.2-75.7 Mb together since both of them correspond to gar LG8 or the same set of shark scaffolds (Supplementary Figure 12). Then, chr1:48.0-52.2 Mb, chr1:75.8-78.0 Mb and chr1: 78.1-196.2 Mb are defined as chr1.2, chr1.3 and chr1.4, respectively. By following the same practice, we back-mutated chicken chr4 and 10 gar-specific fusions (Supplementary Figures 13, 14). Analogously, for three gar-specific fissions (LG9 and LG11, LG1 and LG16, LG3, LG14 and LG17), we manually fused the related chromosomes. Since LG1, LG3, and LG14 have been already back-mutated due to occurrence of ancestral fusions, we re-evaluated the homologous relationships between these LGs and shark scaffolds and then fused the corresponding chromosome groups, which were later referred as LG9LG11, LG1.2LG16, and LG3.1LG14.2LG17.

During the analysis of macrosynteny, we identified six spotted gar-specific chromosomal regions (LG28, LG6.3, LG1.3, LG2.2., LG5.3 and LG24.2) sharing no homology to any chicken chromosome (Supplementary Figure 14). First, gar LG28 does not share an excess of homologs with any chicken chromosome. Second, when back-mutating gar genome we found spotted gar LG1.3, LG5.3 and LG6.3 are not homologous to any chicken chromosomes. Third, to increase the resolution, we performed a sliding-window (window size of 50 genes) analysis by examining whether less than five out of 50 genes could find homologs between chicken and gar. We further examined whether homologous regions are present (more than 10 out of 50 genes with homolog present) in shark and whether the region is not full (25 out of 50 genes) of small-scale duplicates since these duplicates would confound homolog mapping process. With this fine-scale analysis, we additionally found two spotted gar-specific segments: LG2:56.8-68.6 Mb (LG2.2) harbouring 425 genes and LG24:0-4.5 Mb (LG24.2) harbouring 174 genes. For them, only 26 and 13 genes harbour homologs scattered across multiple chicken chromosomes. Accordingly, we named the remaining chromosomal regions as LG2.1 and LG24.1, respectively. Notably, only LG2.2 and LG6.3 could find homologous scaffolds in elephant shark genome, and thus the corresponding regions get lost in chicken.

After back-mutation and identification of gar-specific chromosomal segments, almost all chicken and gar chromosomal segments form one-to-one homology (Supplementary Table 31) except the aforementioned ambiguous case where gar LG2 and LG4 are co-homologous to chicken chr4 and Z, and this co-homology relationship could not be detected in shark scaffolds. With introduction of back-mutation, this case becomes LG2.1 and LG4.2 co-homologous to chicken chr4.2 and Z (Supplementary Table 31).

We reasoned that there should be two separate ancient chromosomes before the split, with each mainly contributing to chicken chrZ (gar LG2.1) and chicken chr4.2 (gar LG4.2) respectively based on the following two observations: 1) chicken chrZ harboured more (355 vs. 56) homologs with gar LG2.1 and chicken chr4.2 harboured more (354 vs. 67) homologs with gar LG4.2 (Supplementary Table 31); 2) both chicken chrZ and gar LG2.1 are homologous to sea cucumber LG01/LG06 and LG08, while chicken chr4.2 and gar LG4.2 are homologous to sea cucumber LG09/LG12 and LG19 (Supplementary Table 31, Supplementary Figure 15). Therefore, we used chicken chrZ (gar LG2.1) and chr4.2 (gar LG4.2) to represent two ancestral gnathostome chromosomes. Their co-homology could be due to recurrent gene shuffling between this pair of chromosomes in both chicken and gar lineages.

**4.5) Macrosynteny between sea cucumber, chicken and spotted gar.**

Consistent with the prediction of WGD, in either chicken or gar genome, almost all individual chromosomes homologous to one sea cucumber LG shares a significant number of homologs between each other (Supplementary Figure 15). For example, chicken chr15 and 19 (homolog of sea cucumber LG04) shared 70 best-to-best paralogs, which was significant and could not happen by chance. And these genes are scattered across the whole chromosome pair (Supplementary Figure 16). More dramatically, the chromosome-level homology is not unique to chr15 and chr19 but applies for the whole chicken genome (Supplementary Figure 16). Thus, we corroborated the previous conclusion that WGD events rather than multiple independent segmental duplications accounts for these observations.

Notably, in four cases, two or three sea cucumber LGs are simultaneously homologous to the same set of three or four chromosomes in either chicken or gar genome (Supplementary Figure 15). They are LG01/LG11/LG17 (chicken: chr2, chr7, chr27, chr33), LG01/LG06 (chicken: chrZ, chr10, chr28, chr25), LG09/LG12 (chicken: chr6, chr4.2, chr13, chr22), and LG16/LG22 (chicken: chr2, chr20, chr3). We proposed that these four groups of sea cucumber LGs are very likely to be fused as one ancestral chromosome before gnathostome WGDs. Consistently, we built a phylogenetic tree covering chicken and gar homologous chromosome sets and sea cucumber LGs (like LG01, LG11 and LG17). We concatenated proteins from the same chromosome and observed the same tree topology for LG01, LG11 and LG17 (Supplementary Figure 17). Sea cucumber LG01/LG06, LG09/LG12 and LG16/LG22 situate in a similar case (Supplementary Figures 18-20). Therefore, these four cases should represent four pre-WGD fused chromosomes.

Given the homologous relationships between sea cucumber LGs and gnathostome chromosomes, the corresponding sea cucumber LGs and gnathostome chromosomes are divided into 17 separate groups (Supplementary Figure 15, Supplementary Table 32) or 17 vertebrate ancestral chromosomes (ACs). These ACs contributed to gnathostome karyotype by 2R WGDs. Of 17 ACs, 11 ACs correspond to four chicken or gar chromosomes and 6 ACs, including AC7, AC8, AC9, AC15, AC16 and AC17 correspond to three (Supplementary Table 32).

**4.6) Reconstruction and evaluation of 17 vertebrate ancestral chromosomes (AC).**

We reconstructed the gene content of 17 ACs based on annotated genes of Belcher's lancelet (Supplementary Table 33) by combining two strategies (Supplementary Figure 21). First, we identified Belcher's lancelet scaffolds homologous to sea cucumber LGs, and then mapped lancelet scaffolds to ACs based on homologous relationships between sea cucumber LGs and ACs. We grouped a lancelet gene into one specific AC if it was located on a scaffold corresponding to this AC , and if there was at least one chicken gene and at least one gar gene, both harboured by chromosomes corresponding to the same AC, have this lancelet gene as their best hit homologous gene. Second, we directly utilised the homology relationship between chicken or gar genes and Belcher's lancelet genes. If at least five chicken and gar genes located on five separate chromosomes, all of which correspond to the same AC, are having the same lancelet gene as their best hit homologous gene. This lancelet gene would be also grouped into the specific AC. Notably, that sea cucumber LG10, which corresponds to AC17, does not have homologous lancelet scaffolds. Together with the fact that most genes of ACs were inferred based on the first strategy, our reconstructed ACs only consists of AC1 to AC16. This is why we left AC17 out in subsequent analyses unless otherwise specified. All in all, 5,045 lancelet genes were anchored to 17 ACs (Supplementary Table 33).

We evaluated the quality of 17 ACs by comparing them to other reconstructed pre-WGD ancestral chromosomes, including 17 tetrads[120], 17 CLGs[121] and 18 PVCs[122]. First, the gene content of 17 tetrads were provided based on human gene annotation, i.e., post-WGD genes[120], we identified their one-way best hit in our AC harboured genes and then inferred correspondence between tetrads and ACs (Supplementary Table 34). Second, 17 CLGs were constructed based on the Florida lancelet genome. We mapped AC harboured genes to Florida lancelet genome release by performing LASTZ based whole genome nucleotide level alignment and inferred the correspondence between ACs and Florida lancelet chromosomes (Supplementary Table 35). Since 17 CLGs were also mapped to chicken and gar chromosomes from its original paper[121], we considered an AC corresponding to one CLG only if they co-correspond to the same Florida lancelet chromosome and if they share the same set of corresponding chicken and gar chromosomes. The only exception is AC17 and its correspondence to CLGs was inferred only based on a shared set of corresponding chicken and gar chromosomes (Supplementary Table 34). Third, the correspondence between PVCs and either CLGs or Tetrads were reported from its original paper[122], we utilised this information to infer the correspondence between ACs and PVCs (Supplementary Table 34).

**4.7) Resolving how 2R WGD leads to the extant chromosomal phylogeny of gnathostomes.**

**4.7.1) Chromosomal phylogeny of Gnathostomes.** With the homology map between sea cucumber LGs, ACs (proxied with genes of Belcher's lancelet), and gnathostomes chromosome set (approximated by one-to-one gar LGs and chicken chromosomes), we attempted to dissect the chromosomal phylogeny of gnathostomes generated by 2R WGDs. Specifically, we defined homologous gene sets as homologs harboured by chicken, spotted gar and amphioxus (Belcher's lancelet unless specified) with the following requirements: (1) for homologs of chicken and gar in one gene sets, their corresponding chromosome needs to map to a single AC (Supplementary Table 36); (2) at least a total of five different chromosomes (chicken) or LGs (spotted gar) needs to be covered, which has a maximum value of eight (four in chicken, and four in spotted gar) under the 2R WGD model; (3) gnathostome genes included in one set have the highest BLASTP scores with the amphioxus gene in the same set compared to remaining genes harboured by chicken or gar genomes. These criteria were applied in all phylogeny related analysis unless otherwise specified. To increase the number of homologous gene sets, we additionally supplemented amphioxus genes from Florida lancelet besides Belcher's lancelet. Among all 696 homologous gene sets (Supplementary Table 36), genes of Florida lancelet constitute only 19 homologous gene sets, and these 19 genes are likely unannotated in Belcher's lancelet.

As aforementioned, the spotted gar LG2.1 and LG4.2 are co-homologous to chicken chr4.2 and Z. In subsequent analysis, we took advantage of co-homology between chrZ (gar LG2.1) and chicken chr4.2 (gar LG4.2) to include more applicable genes. That is, homologous genes on chicken chr4.2 (gar LG4.2) were used to complement homologous genes on chrZ (LG2.1) in AC2 and AC11 related analysis given the correspondence between chrZ (LG2.1) and AC2 or AC11. And homologous genes on chicken chrZ (gar LG2.1) were used to complement homologous genes on chr4.2 (LG4.2) in AC3 and AC4 related analysis. For example, 71 chr4.2 genes and three chrZ genes were added into AC3 homologous gene sets (Supplementary Table 36).

If applicable, we added homologous sea cucumber genes into homologous gene sets to supplement phylogenetic information. Different from chicken and gar, we herein did not strictly follow the correspondence between sea cucumber LGs and ACs (Supplementary Table 36). For example, sea cucumber LG01/LG11/LG17 corresponds to AC1. But in homologous gene sets related to AC1, sea cucumber genes could be harboured by LG02 (1 gene) and LG09 (1 gene) in addition to LG01, LG11 and LG17 (a total of 48 homologous gene sets). The rationale is that translocations may occur due to the remote relationship between sea cucumber and gnathostomes and possible misassembly in sea cucumber may also confound the chromosomal linkage. Therefore, although we used a single sea cucumber linkage group name to denote a sea cucumber outgroup LG in the phylogenetic analyses, like "LG01/LG11/LG17" in (Extended Data Figure 5a), we clarify that the sea cucumber genes from other LGs make a small contribution too (Supplementary Table 36).

With homologous gene sets, we built and polished multiple sequence alignments (MSAs). Specifically, Prank v150803 was used to generate MSA with the parameter '-F' to imply permanent insertions[123]. Since annotation error is pervasive in non-model species[124] (e.g. lamprey), we visualised every MSA in MEGA7 (REF. 125) v7.0.18 and perform a series of manual curation: 1) for tandemly positioned genes in one species but annotated as one gene model in all other species, we interpreted them as artificial gene fission and corrected them back; 2) for alignments consisting of a highly diverged region from one protein, we viewed them as annotation error due to erroneous gene structure and thus discarded this region; 3) for a gene harbouring an alignment gap relative to other homologs, we interpreted it as a missing exon, analysed such region with BLAT and UCSC genome browser[125–127] given the surrounding alignable region of this gene and performed the correction according to annotation tracks of UCSC, NCBI orf-finder or TBLASTN gateways and the latest Ensembl annotation track of the same or closely related species. In total we corrected 1,957 sequences from all these species and attached the correct sequences in Supplementary File 4. After correction, gaps and poorly aligned regions were automatically filtered using Gblocks v0.91b with parameters as ‘’-b3=10 -b4=5 -b5=h” specifying a loose criteria, which is generally more superior than the default stringent settings when searching for ML tree based on high divergence alignments[128]. By contrast, for gene level tree analyses with much limited information, we did not implement Gblocks since filtering alignments sometimes leads to a worse result and the more aggressive filtering strategy is, the less reliable trees would be finally obtained[129].

We then performed phylogenetic tree reconstruction. For any MSA, we used ModelFinder with BIC criteria to find the best-fitting evolutionary model and with the parameter ‘-mtree’ to enable an exhaustive search[130]. To infer chromosome level phylogeny, MSAs corresponding to the same AC are concatenated to form a super matrix. Specifically, outgroup genes from sea cucumber or amphioxus were separately concatenated, vertebrate genes from the same chicken or gar chromosomes were concatenated. Each MSA within the super matrix was defined as a separate partition with dedicated evolutionary models[131]. We concurrently ran RAxML-ng[132] v0.9.0 and IQ-TREE[133] v1.6.12 to infer chromosome level maximum-likelihood (ML) trees. For RAxML-ng, bootstraps were automatically implemented and bootstrap supporting values were calculated with the TBE method[134]. For IQ-TREE, 1000 ultrafast bootstraps were conducted. Finally, by applying a coalescence approach to generate a chromosomal level consensus tree, all bootstrapped MSAs corresponding to the same AC were grouped as the input to Astral-III[135], version 5.6.3.

**4.7.2) Chromosomal fusion and karyotype evolution before, between and after 2R WGDs.** By following the parsimony rule, we reconstructed how gnathostome karyotype evolved around 2R WGDs (Extended Data Figure 5b). After the first round of WGD, 34 chromosomes were derived from 17 ACs. The correspondence between ACs and back-mutated chicken/gar chromosomes indicates eight post-1R fusions, in which one fusion event involved three chromosomes (Supplementary Figure 22). Thus, 34 chromosomes became 25 chromosomes. After the second round of WGD, 50 chromosomes emerged. Six chromosomes, including one copy derived from AC7, AC8, AC9, AC15, AC16 and AC17, respectively, appeared to be lost before the split of chicken and gar because these ACs correspond to 3 modern chromosomal sections, instead of four (Extended Data Figure 5). Therefore, only 44 chromosomes were retained. Four post-2R fusions occurred before the split of chicken and gar lineage, leading to chicken chr2 (gar LG9LG11), chicken chr3 (gar LG1.2LG16), chicken chr1.4 (gar LG3.1LG14.2LG17) and gar LG2.2 separately. Therefore, 40 chromosomes were present before the split of chicken and gar lineages. Six chromosomes were then lost specifically in chicken resulting in the aforementioned six gar-specific chromosomal segments.

There are two notable issues: (1) we only showed the evolutionary process from 17 ACs to 50 chromosomes after the second rounds of WGD in Extended Data Figure 5b, since post-2R events were irrelevant with our major focus; (2) except for AC17 (not reconstructed) derived chromosomes, the sizes of the other chromosomes in Extended Data Figure 5b were predicted as the total count of retained genes from chicken and gar orthologous chromosomes for post-2R chromosomes, and the total count of retained genes from two gnathostome chromosomes (reconstructed based on chicken and gar) for post-1R chromosomes.

**4.7.3) Comparing the gnathostome chromosomal fusions in our model to previous models.** We evaluated how our pre-1R and post-1R chromosomal fusion events differ from previous literatures. Pre-1R fusions were not formally discussed in the 17-tetrad model[120] or the 17-CLG model[121]. Post-1R fusions are provided as tetrad pairs, CLG pairs, or PVCs pairs from the original papers. Since we already mapped 17 ACs to either 17 tetrads, 17 CLGs or 18 PVCs (Supplementary Table 34), the correspondence of post-1R fusions is directly available. All post-1R fusions identified in previous models are reconciled in our model, though minor differences exist when it is related to AC3 (Supplementary Table 37).

**4.7.4) Simulating pairwise post-1R fusions.** Under our model of 17 ACs, eight post-1R fusion events occurred. However, Under the pairwise "post-1R fusion" model, AC3 did not represent a pre-WGD fused chromosome but instead represented two separate pre-WGD chromosomes that happened to have fused twice after the first round of WGD (Supplementary Figure 23). In this scenario, 10 post-1R fusion events occurred, which include one post-1R pairwise fusion. To estimate the likelihood of a pairwise fusion model, we simulated by introducing 10 random post-1R fusions and recording the number of pairwise fusions. We performed three technical replicates with each replicate including 10,000 independent simulations (Supplementary Table 38).

**4.8) Phylogenetic analyses of cyclostomes and gnathostomes.**

To decipher whether cyclostomes and gnathostomes share WGD(s), we anchored cyclostome genes into the aforementioned phylogenetic trees of gnathostomes. That is, we incorporated genes from shark, hagfish and lamprey based on homologous amphioxus genes, built MSAs and reconstructed maximum likelihood (ML) trees (Supplementary Table 39). To increase phylogenetic information, we also added homologous Florida lancelet genes of all vertebrate genes in a homologous gene set. For each single MSA, we separately and repeatedly run RAxML-ng and IQ-TREE for 10 times using different seed numbers to increase the power of ML tree search[136]. For the 20 obtained ML trees for each MSA, we used RAxML-ng to re-evaluate their likelihoods and chose the best tree as the final tree for a MSA.

To increase the number of applicable trees, we performed three separate analyses by only incorporating hagfish genes, lamprey genes, or genes of hagfish and lamprey, respectively (Supplementary Files 5-8). For each separate trial, shark genes were included as a positive control given their stable topological positions.

To choose high-quality phylogenetic trees, we implemented the ETE toolkit[137] to require that all bootstrap values in a single gene tree should be greater than 70%. The only exception is the monophyletic clade consisting of a shark gene, a gar gene and a chicken gene, for which the cutoff was relaxed to 60%. The reason is due to possible incomplete lineage sorting wherein shark genes may cluster more closely with chicken or gar genes. However, the internal topology of this clade did not affect the position of cyclostome genes.

We finally manually examined all trees to require that the topology of chicken and gar genes were consistent with the chromosome level tree. We required that the phylogenetic relationships of chicken genes and gar genes exactly recapitulated the chromosome level trees depicted in aforementioned gnathostome chromosome level tree (Extended Data Figure 5a), since only these trees provide information to evaluate the positions of cyclostome genes relative to two rounds of WGDs.

**4.9) Investigating whether lamprey lineage shared gnathostome pre-1R and post-1R fusions.**

As aforementioned, we predicted four pre-1R fusions, e.g., AC1 emerged after pre-1R fusion of sea cucumber LG01/LG11/LG17 (Supplementary Figure 17). We thus analysed homologous relationships between sea cucumber LGs and sea lamprey scaffolds. For example, if any one lamprey scaffold is homologous to all LG01, LG11 and LG17 of the sea cucumber assembly, we deemed that a pre-1R fusion of sea cucumber LG01/LG11/LG17 is shared by sea lamprey genome. We found evidence in the sea lamprey for 3 out of 4 of the pre-1R fusions (Supplementary Figure 24) As aforementioned, we have also predicted eight post-1R fusions (Supplementary Figure 22). The strategy to judge sharing of post-1R fusion follows the same rule as pre-1R fusion.

**4.10) Gene retention profile-based analyses.**

**4.10.1) Definition of gene retention profile and overlapping ratio.** To enable phylogenetic analyses between cyclostomes and ACs, we implemented a novel summary statistic, i.e., gene retention profile. It was defined as a vector listing the presence or absence of homologous genes on a vertebrate chromosome for all genes of the corresponding AC. Since an AC contains genes harboured by different Belcher's lancelet scaffolds, the order information of genes on an AC is unknown. So, when drawing the gene retention profile of a chromosome, the genes of one AC were firstly randomly ranked by the lancelet scaffold it originally belongs to and then by the real gene order within that scaffold (Supplementary Files 10, 11). Certainly, the order is irrelevant for all phylogenetic analyses, which were only based on presence or absence information. For two chromosomes corresponding to the same AC, the overlapping ratio was calculated based on two gene retention profiles or vectors (Figure 4b). To enhance robustness, we required at least 20 retained genes of an AC on descendent chromosomes before calculation of the overlapping ratio.

To make overlapping ratio comparable between gnathostomes and cyclostomes, we corrected a bias inherited during reconstruction of the gene content of ACs based on amphioxus genes: genes encoded by an AC must share homologs in corresponding homologous chromosomes of both chicken and gar, while hagfish and lamprey genes were not considered. Therefore, the genes on ACs should be biassed towards gnathostome-specifically retained genes. We corrected the bias by requiring the genes on ACs to share homologous genes in corresponding chromosomes of both hagfish and lamprey. Before the correction, 4,325 out of 5,045 genes on all ACs were retained in either hagfish or sea lamprey. After the correction, 3,081 genes were retained and used in cyclostome related analyses. Note, the cut-off (0.15 in Figure 4c-e) was derived based on all 5,045 genes on ACs. If only 3,081 genes were used, the cut-off (0.15) remained robust.

**4.10.2) Threshold of overlapping ratio to discriminate homologous chromosomes.** We generated the random expectation of overlapping regions by artificially splitting a chromosome into two parts and calculating the ratios. In other words, for a vertebrate chromosome corresponding to one AC, we ordered the homologous genes by their actual positions on the chromosome and manually split these homologous gene into two parts given the 20%, 30%, 40%, 50%, 60%, 70% and 80% quantile rank position. Then, if each part harbours at least 20 retained genes, we calculated their overlapping ratios. Therefore, for each chromosome and the corresponding AC, there would be up to seven ratios, the maximum value of which were specified as the final ratio for this chromosome. Two rationales were invoked in this simulation: 1) we could not randomly sample two chromosomes since they may be duplicates derived from one AC; 2) we chose the maximum rather than median value of seven ratios to be conservative, i.e., to generate an upper-bound estimation of overlapping ratios of two random chromosomal segments. Finally, given each chromosome and the corresponding AC, there are a total of 49, 45, 49 and 46 simulated split results in chicken, gar, hagfish and sea lamprey genome, respectively (Supplementary Table 40). By comparing the distribution of overlapping ratios for artificially split chromosome pairs and bona fide paralogous chromosome pairs in gnathostomes, we reasoned 0.15 should serve as a good threshold.

**4.10.3) Application of overlapping ratio.** We performed a series of analyses based on overlapping ratios: (1) we identified two chromosomes as paralogous chromosomes when their overlapping ratio exceeds the cut-off of 0.15, which was specified based on the comparison of *OR* values between randomly split chromosome pairs and that of real paralogous chromosome pairs generated by WGD from chicken and gar data (Figure 4c, c); (2) we identified the orthologous cyclostomes chromosomes as a pair of hagfish and lamprey chromosomes that formed a single clade in the clustering of chromosomes derived from the same AC based on gene retention profile (Supplementary Files 10, 11). For (2), the distance was equal to one minus overlapping ratio and the clustering was then conducted based on the distance matrix of these gene retention profiles with a "ward.D2" method embedded in the R "pheatmap" package (Supplementary Tables 41, 42).

**4.10.4) Splitting three hagfish chromosomes identified further cyclostome orthology.** We found that hagfish cluster9 (corresponding to AC10), cluster14 (AC10) and cluster16 (AC16) show peculiar overlapping ratio patterns after manual split and examined why. Specifically, during the previous chromosomal split analyses in hagfish and sea lamprey, we noticed that two parts in these three hagfish clusters show overlapping ratios higher than 0.15 (Supplementary Table 40), suggesting chromosomal fusion of two paralogous chromosomes in hagfish evolution. We therefore manually split these hagfish clusters to examine whether two parts would separately cluster with two lamprey scaffolds (Supplementary Figure 25). To determine the boundary, we first obtained all homologous genes on the hagfish cluster that corresponds to a specific AC and sorted them according to their actual position. We then exhaustively scan all paralogs to search for a boundary to maximise the total number of shared retained genes between the two hagfish parts. We finally split the hagfish cluster on this position and redo the overlapping ratio-based clustering (Supplementary Figure 25).

**4.10.5) Simulating the decay of overlapping ratio after a third round of WGD.** We expected that the overlapping ratio of any sister chromosome pairs drops with the increase of the number of WGD(s) since only a proportion of duplicates could be retained and retained duplicates could be scattered to more paralogous chromosomes. To make a more quantitative inference, we made a mathematical deduction (Supplementary Figure 26) based on two simple but reasonable assumptions: (1) the gene loss and gene retention are symmetrical; (2) all chromosomes have the same duplicate gene retention rate after a WGD. Specifically, for any paralogous chromosome pair, i.e., chr1 and chr2, their overlapping ratio (*OR*) is defined as,

$$OR(raw) = \frac{a}{min\,(a+b, a+c)}$$

After an additional round of WGD with duplicate gene retention rate $p$, two chromosomes became four chromosomes, which are named as chr1.1, chr1.2, chr2.1 and chr2.2. The *OR* between sister pairs (e.g., chr1.1 and chr1.2) is only dependent on the duplicate gene retention rate:

$$OR(new.1) = \frac{2p}{1+p}$$

while the *OR* between homologous chromosome pairs not directly generated by WGD (e.g., chr1.1 and chr2.1, chr1.2 and chr2.1) is

$$OR(new.2) = \frac{1+p}{2} * OR(raw)$$

Now, we would like to know the impact of a third round of WGD on *OR* among paralogous chromosomes. After two rounds of WGDs, a maximum of four paralogous chromosomes are generated and the average *OR* for two out of four paralogous chromosomes is defined as

$$OR(2R)$$

After a third round of WGD, four paralogous chromosomes became eight paralogous chromosomes. And there are in total 28 different combinations of two paralogous chromosome pairs from 8 paralogous chromosomes, of which 4 chromosome pairs were directly generated by the third round of WGD and the remaining 24 not directly generated by the third round of WGD. For the former 4 sister chromosome pairs, their OR again is

$$OR(3R.1) = \frac{2p}{1+p}$$

For the later 24 pairs, their overlapped ratio is

$$OR(3R.2) = \frac{1+p}{2} * OR(2R)$$

Therefore, the average *OR* between any 2 out of 8 paralogous chromosomes after the third round of WGD is

$$OR(3R) = \frac{4}{28} * OR(3R.1) + \frac{24}{28} * OR(3R.2) = \frac{4}{28} * \frac{2p}{1+p} + \frac{24}{28} * \frac{1+p}{2} * OR(2R)$$

We then approximated *OR(2R)* in cyclostomes with the median *OR* of paralogous chromosomes contributed by 2R WGD (0.52) in gnathostomes (Supplementary Table 43). That is,

$$OR(2R) = 0.52$$

For teleost which was subject to the third round of WGD about 320-350 million years (myr) ago, the duplicate gene retention rate in extant species were 0.15-0.20 (REFs. 138-140). We used this range to approximate the value of $p$ for the third round of cyclostome WGD, which likely occurred before the split of hagfish and lamprey, that is, earlier than 470 mya. Performing in silico simulation with a reasonable p ranging between 0.08 and 0.30, we found that the mean *OR* after the third round of WGD is generally between 0.26 and 0.36. That is,

$$OR(3R) \sim (0.26, 0.36)$$

Thus, as expected, after a third round of WGD, the mean *OR* of paralogous chromosomes would further drop. Actually, the observed median *OR* in cyclostome species is 0.30 (Supplementary Table 44), fitting well with the simulated scenario where cyclostomes were subject to three rounds of WGDs.

**4.11) Further scaffolding of sea lamprey scaffolds.**

We took advantage of closely related Pacific lamprey linkage group information to further increase the continuity of sea lamprey assembly. That is, although the sea lamprey genome used in this study has been assembled at the pseudo-chromosome level, numerous scaffolds exist. So, we extracted the orthologous mapping information provided by Smith et al.[141]. When more than one sea lamprey scaffolds were mapped to the same Pacific lamprey LG, and when they correspond to the same AC, we interpreted such signal as incomplete assembly of one chromosome and manually fused them, which was represented as the scaffold name of the larger scaffold (Supplementary Table 45). 17 out of 18 fusion cases were confirmed by the updated sea lamprey genome release[142]. The further scaffolded sea lamprey pseudo-chromosomes are used in our overlapping ratio-based analyses.

**4.12) Estimating the chance of paralogous chromosomal fusions in hagfish evolution.** We wondered how frequent paralogous chromosomal fusions occurred in hagfish lineage since such events confound the analyses of overlapping ratios. For example, both hagfish cluster1 and cluster6 correspond to AC10, and cluster1 is paralogous to cluster6 on the AC10 derived regions according to their overlapping ratio. We firstly randomly shuffled the correspondence between ACs and hagfish clusters. We then counted the cases where two paralogous parts were shuffled to the same hagfish cluster. For example, if the cluster1-AC10 part and cluter6-AC10 part were shuffled into one cluster, it was counted as one fusion event between paralogous chromosomes. We performed two replicates of simulations with each replication consisting of 10,000 simulations (Supplementary Figure 27). The interquartile range (IQR) of paralogous chromosome fusion events were estimated to be between 3 and 6.

### 5. REGULATORY EVOLUTION

#### 5.1) *Cis*-regulatory profiling.

A previous comparative analysis of gene regulatory elements between amphioxus and several gnathostomes suggested an increase in distal gene regulatory elements during the vertebrate evolution[4]. Considering that distal regulatory elements could contribute to spatiotemporal gene expression patterns, the evolutionary increase of distal regulatory elements may lead to tissue- or cell type-specific evolution. However, it is still unknown whether this increase in complexity is a shared feature of all vertebrates. Thus, here we examined whether hagfish also have more distal regulatory elements in their genomes in order to understand general trends of gene regulatory evolution after whole genome duplication events.

We collected hagfish embryos at two different developmental stages and separated each embryo into anterior and posterior parts. Using them, we performed an assay for transposase-accessible chromatin using sequencing (ATAC-seq)[143] per duplicate (technical replicates) to estimate gene regulatory regions in the hagfish genome. For each sample, we identified more than 130,000 accessible chromatin regions (ACRs) as putative regulatory regions (Supplementary Figure 29a). It should be noted that identified hagfish ACRs would not have any information on biological variation, as we have no biological replicates but only technical replicates in our ATAC-seq data because of the extreme difficulty in hagfish sampling, especially of embryonic material.

We next confirmed that accessible chromatin landscapes determined by our hagfish ATAC-seq data were reliable by several analyses. First, we examined the fragment sizes of ATAC-seq for each sample, and confirmed the periodic distributions corresponding to integer multiples of nucleosomes (Supplementary Figure 29b). Second, quantifying the fraction of all aligned reads in the ACRs (FRiP scores), we confirmed a high signal-to-background ratio of all ATAC-seq data (Supplementary Figure 29c). In addition, identified ACRs were not distributed randomly through the genome but significantly enriched at promoter regions of transcription start sites (TSSs) (Supplementary Table 46). Moreover, the ATAC-seq signal intensities at ACRs were robustly reproduced between the two technical replicates (Supplementary Figure 29d). Collectively, the obtained results indicate that our embryonic ATAC-seq data provide a reliable accessible chromatin landscape of the hagfish genome.

To compare the genome-wide distribution of ACRs in hagfish with non-vertebrate chordate and gnathostome species, we used publicly available ATAC-seq datasets of amphioxus (*Branchiostoma lanceolatum*) and four gnathostome species (*Danio rerio*, *Oryzias latipes*, *Gallus gallus*, and *Mus musculus*)[4,144]. For all the species, we identified ACRs, and then divided them into promoter, proximal, exonic, and distal ACRs according to the distance from the closest TSSs. Consistent with the previous study[4], all the gnathostomes have smaller fractions of ACRs in the promoter or the proximal regions than those in the amphioxus, and we also observed essentially the same tendency in the hagfish (Figure 1a and Supplementary Tables 47, 48). As for distal ACRs, the hagfish and all the gnathostomes also similarly showed significant enrichments, which was not detected in the amphioxus (Figure 1a and Supplementary Tables 47, 48). Next, examining the distance between ACRs and the closest TSSs, we found that the distances in the hagfish and all the gnathostomes were significantly larger than those in the amphioxus (Figure 1b and Supplementary Table 49; Extended Data Figure 8c). In addition, we also observed that the higher number of ACRs in the *cis*-regulatory region of each gene (as defined in REF. 145) not only in all the gnathostomes, as shown previously[4], but also in the hagfish, by comparing with the amphioxus

(Figure 1c and Supplementary Table 50). This suggests that evolutionary increase in the regulatory regions per gene during the vertebrate evolution.

It is still unknown whether the evolutionary increase has occurred independently in the gnathostome and the hagfish lineage, and we cannot deny the possibility of decrease in the regulatory regions in amphioxus. Nonetheless, distal regulatory elements would mainly contribute to the differences in regulatory complexity between vertebrates and amphioxus.

**5.2) Experimental and computational procedures.**

**5.2.1) Defining ohnologs.** Because the homology between vertebrate chromosomes and ACs have already been established (Extended Data Figure 5a), a group of genes locating on different chromosomes but sharing the same homologous gene on the corresponding AC was considered an ohnolog family. To minimize the confounding effects from tandem duplicated genes, we required the count of homologous genes should be no more than one plus the count of unique chromosomes where these genes are located. It means for one ohnolog family, at most one tandem duplicate gene may be included. As a result, 2,137 hagfish and 3,594 chicken ohnologs were identified.

**5.2.2) Gene Ontology of ohnologs.** We mapped hagfish and chicken ohnologs to human genes and performed GO enrichment analysis with human homologs since human genes have far more comprehensive ontology annotation than hagfish and chicken. For 2,137 ohnologs and all 16,473 protein coding genes in hagfish, we identified 959 and 8,293 reciprocal best hits in the human genome, respectively. For 3,594 ohnologs and all 18,346 protein protein coding genes in the chicken genome, we identified 3,288 and 12,875 reciprocal best hits, respectively. The larger gene list served as the statistical background. Functional enrichment was examined with the Metascape[146] online tool. We used the 959 human orthologs of hagfish ohnologs and randomly sampled 2,999 genes (as Metascape has a limit of 3,000) from a total of 3,594 chicken orthologs. We performed GO (Biological Process only) enrichment analysis against all genes of the two species. The top 20 enriched functions of ohnologs of two species and their parental GO terms have been shown in Extended Data Figure 8a,b. Semantic similarity score[147] between the two sets of enriched GO terms was 0.694, higher than random expectation (0.20~0.45)[148]. Furthermore, genes were annotated as either developmental ohnologs, non-developmental ohnologs or non-ohnologous genes according to their GO terms annotated by PANTHER[149].

**5.2.3) ATAC-seq of hagfish embryos.** Embryonic samples of the hagfish are extremely rare[150]. In this study, we collected two embryos of the inshore hagfish *E. burgeri*, at stages Dean 45 (collected in 2018) and 53 (collected in 2017) (REF. 151; Supplementary Figure 28), and performed ATAC-seq experiments following previous descriptions[143,152] with slight variations. Embryos were divided into two parts, anterior and posterior halves, at the level of the somite pair 27 (Supplementary Figure 28), in order to gain positional information for future projects. Nuclei were extracted separately from each part using 2 mL glass bouncers in ice-cold PBS and lysed in cold lysis buffer (10 mM Tris pH 7.4, 10 mM NaCl, 3 mM MgCl2 and 0.1% Igepal). Each pool was further divided into two to obtain data from technical replicates (given the lack of biological replication). Transposition reaction was done on ~50,000 nuclei per sample and per replicate (total of 200 k cells per embryo), with Tn5 enzyme (Illumina Nextera DNA Library Prep Kit, #FC-121-1030), during 5 min at 37 ºC. DNA was purified using a QIAGEN MinElute kit following manufacturer instructions. Number of PCR cycles per sample were first determined as previously described(Buenrostro et al., 2013) using 1 uL of the Tn5'ed chromatin, and then a final PCR reaction with the rest of the chromatin was performed for each sample, with the following number of cycles: 7 for the embryo at stage 45, and 8 cycles for the embryo at stage 53. PCRs were performed with the NEBNext High-Fidelity 2x PCR Master Mix (New England Biolabs, #M0541), using primers Ad1F and Different sets of primers were used to multiplex the libraries, and obtained from Buenrostro et al.[143]. Primer combinations per sample used are as follows:

- stage 45 anterior 1: Ad2.5R; stage 45 anterior 2: Ad2.7R;
- stage 45 posterior 1: Ad2.6R; stage 45 posterior 2: Ad2.11R;
- stage 53 anterior 1:Ad2.3R; stage 53 anterior 2: Ad2.9R;
- stage 53 posterior 1, Ad2.4R; stage 53 posterior 2, Ad2.10R.

The resulting multiplexed libraries were checked in an Agilent 2100 BioAnalyzer using High Sensitivity DNA chips (nucleosome pattern was observed in the profiles) and sequenced in four lanes (2 lanes per embryo) of an Illumina HiSeq 4000 platform, at the Beijing Genome Institute (BGI).

**5.2.4) ATAC-seq data of amphioxus, zebrafish, medaka, chicken, and mouse.** For the ATAC-seq analysis of species except for hagfish, we used published ATAC-seq datasets (Accession ID: GSE106428 for amphioxus and zebrafish[4]; DRA006971 for medaka, chicken, and mouse[144] (Supplementary Table 51). While there are two replicate data for hagfish, amphioxus, and zebrafish, there are three replicate data for each sample for medaka, chicken, and mouse ATAC-seq data. In order to minimise analytical biases derived from the different numbers of the replicate ATAC-seq data, we removed the ATAC-seq data with the lowest number of aligned ATAC-seq reads for medaka, chicken, and mouse, so that we can use the same number of replicate data across different species data in the ATAC-seq analysis.

**5.2.5) ATAC-seq alignment and peak calling.** Adaptor removal of raw paired-end reads were performed by using NGmerge[153] (version 0.3) with the following parameters: -a -e 30. The filtered reads were aligned to species-specific reference genomes (this study for *E. burgeri*, BraLan2 for *Branchiostoma_lanceolatum*[4], GRCz11 for *Danio rerio*[154], ASM223467v1 for *Oryzias latipes*[155], GRCg6a for *Gallus gallus (International Chicken Genome Sequencing Consortium 2004)*, GRCm38 for *Mus musculus*[156]) by using bowtie2 (version 2.4.2) (REF. 13) with the following parameters: -k 4 -X 2000 --sensitive. These genomes were downloaded from the Ensembl database (release 101)[157]. Picard (version 2.23.8; http://broadinstitute.github.io/picard/index.html) was used to remove duplicate reads from properly paired aligned reads, which had been filtered by using SAMtools[158] (version 1.10). For peak calling, we basically followed the pipeline described in Marlétaz et al.[4]. First, to deplete ATAC-seq reads derived from the transposition events that happened on both sides of a nucleosome, we removed the properly paired reads whose mate is more than 120 bp for identification of accessible chromatin reginos (ACRs). Then, the start position of every aligned ATAC-seq read was corrected to account for the 9-bp insert between the adaptors, which was introduced by Tn5 transposase[143], and the single 5′-most base of each read was retained to obtain single-base resolution. For each sample, the data for the two replicates were pooled into a single dataset. For the pooled data and those for each replicate data, peak calling was performed by using MACS[159] (version 2.2.7.1) with the following parameters: callpeak --nomodel --keepdup 1 --llocal 10000 --extsize 74 --shift -37 -p 0.07. Finally, on the basis of replicate information, we obtained the high confidence ACRs by using the IDR framework[160] (idr 2.0.4 with the following parameter: -i 0.1). To evaluate the reproducibility of chromatin accessibility between replicates, we calculated Pearson correlation coefficients by using the ATAC-seq signal intensities (log10-RPM; reads per million mapped reads) of ACRs at different samples.

**5.2.6) Definition of the *cis*-regulatory regions.** We defined the *cis*-regulatory region of each gene according to the ‘Genomic Regions Enrichment of Annotations Tool’ (GREAT[145]). First, each gene is assigned a proximal regulatory region that extends 5 kb upstream and 1 kb downstream of the TSS, or until another TSS is found. Next, the proximal region is extended up to 1 Mbp in both directions, until another proximal region is encountered. This extended region was defined as the *cis*-regulatory regions of the gene. Peaks falling into *cis*-regulatory region of one gene are considered to be related to the regulation of transcription of it.

**5.2.7) The genomic distribution of ACRs.** To examine the genomic distribution of ACRs in each species, all the ACRs at all the samples were consolidated into a single list. In the merged list, ACRs overlapping each other were combined into a single ACR. By using bedtools[24] (version 2.29.2) and the longest transcript for each gene, these ACRs were classified into four groups: (1) ACRs in promoter regions were defined as those overlapping with regions between 1 kbp upstream and 500 bp downstream of all TSSs of genes; (2) ACRs in proximal regions were defined as those overlapping with regions between 5 kb upstream and 1 kb downstream of all the TSSs but not overlapping with promoters; (3) ACRs in exons were defined as those overlapping with exons of genes but not overlapping with proximal regions; (4) distal ACRs were defined as those not in groups 1–3. In this analysis, we used the gene sets retrieved from the Ensembl database[157] except for hagfish. Considering the length of GREAT regions are different, we also normalised the length of GREAT region to examine the peak density of genes with different roles in development (Extended Data Figure 8d).

**5.2.9) Fate of ohnologs after WGD.** As aforementioned, hagfish transcriptome data of nine somatic tissues were generated in this study. Chicken transcriptome data of nine somatic tissues were retrieved from public sources[161]. Gene level transcriptomic quantification was performed with RSEM[162] software (version 1.3.1) using STAR[163] mapping method (version 2.6.1d). Following previous practice[4], we normalized transcriptome data of different tissues with the quantile method by in-house scripts (code provided in Supplementary File 12). After normalization, TPM (transcripts per million) > 5 is used as threshold to binarize a gene state as either 'expressed' or 'not expressed'. Tissue specificity is quantitatively described by Tau index[164]; we chose 0.9 value as a cut-off to define tissue specific genes.

Herein, we only analysed ohnolog families containing exactly two ohnologs. For an ohnolog pair, we required both genes to be expressed in at least one tissue of tissues assayed in this study for the hagfish or retrieved from public databases for the chicken[161]. Since we did not have information about expressional domains of orthologous amphioxus genes in homologous tissues, we followed a different strategy than that of Marlétaz et al.[4] to define the expressional fates of ohnologous genes. Ohnolog pairs were classified into three classes based on their expressional states across tissues[4] (Supplementary Figure 30): (i) redundancy, if the two ohnologs are expressed in the same set of tissues; subfunctionalization, if both ohnologs are each expressed in a tissue not shared with the other. In other words, each of them has tissue-specific expression domains; (iii) specializatoin, if one ohnolog has a reduced set of expression domains contained in a larger set of expressional tissues of the other ohnolog. Gene families within 'specialization' can be further defined as 'strong specialization' if the number of expressed tissues for the ohnolog with the narrower expression range is less than forty percent of its ohnolog.

### 6. DATING WGD EVENTS AND ASSESSING THEIR MORPHOLOGICAL EFFECT

#### 6.1) Background.

WGD events have undoubtedly contributed to the expansion of gene families underpinning key developmental processes, but they have also been implicated causally in the evolution of the vertebrate body plan[165–169]. The relative timing of vertebrate WGD events has long been thought to coincide with dramatic increases or bursts in phenotypic innovation[165,170]. Testing such hypotheses of causality requires both an estimate of the absolute (geological) timing of the event and a measure of phenotypic diversity (disparity) pre- and post-WGD. The timing of a WGD event can be constrained relative to the timing of lineage divergence events before and after a WGD event, however, this approach is unnecessarily imprecise when applied to events that occur on long evolutionary branches, as here. Instead, molecular clock methods can be applied to directly date WGD events by exploiting the signal of paralogy in gene families where ohnologs have been retained[171,172]. Molecular clock analyses based on gene trees can be calibrated in the same way as species trees, using temporal constraints on the timing of speciation events within gene trees. This approach combines information from the fossil record with Bayesian MCMC methods to estimate probabilistically the age of a gene or genome duplication event. Existing applications have yielded precise estimates for the timing of WGD events in deep time[171,173,174].

WGD events in vertebrates have been inferred on important branches on the vertebrate tree with respect to the key evolutionary innovations including stem-vertebrates, stem-gnathostomes and stem-cyclostomes. Fossil evidence demonstrates that at the very least in the stem-gnathostomes phenotypic innovation played out over a protracted time[72,175]. If WGD events were causal to developmental and phenotypic innovation on these lineages, we should anticipate that they occurred prior to the appearance of key phenotypes. Alternatively, if the relationship between WGD events, developmental and phenotypic innovation is complementary, WGD would occur after the appearance of these characteristics[175]. Here, we attempt to test these competing hypotheses by establishing the timing of WGD events relative to phenotypic innovation manifest as phenotypic disparity. Phenotypic disparity describes the variance in form between species and lineages[176]. Estimates of disparity can be obtained from living and fossil species and, in a phylogenetic framework, can be used to contrast patterns of phenotypic diversity before and after WGD events[173,177–179]. With that in mind, here we employ Bayesian divergence time estimation to place these genome duplication events temporally, relating this to the evolution of phenotypic disparity within the clade.

**6.2) Dating Whole Genome Duplications.**

The age of the 1R duplication event is inferred to be 536-525 Ma (early Cambrian; Figure 2, Supplementary Figure 31). Across both subtrees, the age of crown vertebrates was estimated at 510-502 Ma (middle Cambrian), suggesting a lag of 15-34 Myr between the WGD event and the divergence of crown vertebrates. The age of the 2R event was estimated at 498-486 Ma (late Cambrian) across both subtrees, with the age of the crown gnathostomes between 450-445 Ma (Katian, Late Ordovician; Figure 2, Supplementary Figure 31), suggesting a longer lag of between 46-53 Myr. The CR1 and CR2 events are dated in a rapid succession to 505-494 (late Cambrian) and 491-473 Ma (latest Cambrian - earliest Ordovician), respectively, 10-39 Myr prior to the 463-452 Ma (Middle-Late Ordovician) divergence of crown-cyclostomes.

**6.3) Phenotypic disparity.**

The phylomorphospace exhibits clustering of clades along both axes, as anticipated, though with stem representatives often plotting intermediate of their respective crown-clades and their nearest living clades. For example, stem-cyclostomes occupy a position within the morphospace that is intermediate of invertebrate deuterostomes and crown-cyclostomes. Similarly, ostracoderms (stem-gnathostomes) are positioned intermediate of cyclostomes and jawed vertebrates, i.e. placoderms+crown gnathostomes (Figure 5a).

Generally, the occupation of morphospace exhibits a strong phylogenetic structure, with clades occupying distinct regions of morphospace. However, placoderms are nested entirely among the chondrichthyan and osteichthyan total groups and, similarly, thelodonts nest within pteraspidomorphs.

Each duplication coincides with an increase in overall disparity and the exploration of a novel region of morphospace (Figure 5b), but the majority of vertebrate disparity emerged subsequent to the 2R WGD event – between 88-97% of the morphospace encompassed by a vertebrate convex hull is attributable to descendants of the 2R event.

**6.4) Discussion.**

Here we have addressed a well-studied problem in phylogenetics, regarding the phylogenetic position of the 2R event in vertebrate evolution. Our high-throughput approach considers all gene families, to incorporate as much evidence as possible without focusing on selected individual gene families. We expect inferred gene duplication events from OrthoFinder to be accurate, because it has some *a priori* expectations of gene tree errors, which it corrects for while predicting duplications[180]. This avoids inferring very large numbers of losses for individual gene families, because in these cases, a gene tree reconstruction error is likely to be a better explanation for the inferred tree. Further, we only report duplication events from OrthoFinder that have greater than 50% support. Here support means the number of species that we would expect to have both duplicates present as paralogs, without loss events. This means that gene loss is allowed, but duplication events followed by extensive loss would be ignored, because there is a distinct possibility that these patterns represent the consequence of a phylogenetic, gene-tree reconstruction, error. This would further reduce the effect of stochastic gene-tree reconstruction error, at the expense of some correctly inferred gene trees, but this strategy has the advantage of being conservative, only identifying a WGD when substantial, consistent data is available. As a measure to validate our approach we considered whether it robustly and correctly identified uncontroversial vertebrate WGD, i.e. the 1R and 3R (teleost) duplication events, which are here used as positive controls.

In this analysis, we do not distinguish between whole genome duplication and other forms of large-scale gene duplication, and a large-scale duplication does not necessarily correspond to a WGD event. Whole genome duplications appear to be exceptional events in animals, although WGDs have been identified in diverse lineages[181–183]. We are therefore cautious in making conclusions about the potential for WGDs to have happened along branches where they are not expected, based on our results only. However, the evidence we present strongly suggests the 2R duplication happened after the divergence of gnathostomes and cyclostomes. In addition, we identify two duplication events in cyclostomes. The four duplications we identify close to the vertebrate root (1R, 2R and cyclostome duplications) lead to potentially confusing gene-tree patterns, and we argue that these confusing patterns underpinned erroneous suggestions that 2R occurred in stem vertebrate lineage, or not at all.

We present the first estimates for the timing of the four genome duplication events associated with early vertebrate evolution (Figure 2). Compared to previous estimates, our results represent a significant advancement in that (i) they are derived through integration of palaeontological and molecular evidence, and (ii) they simultaneously estimate the age of the WGD and speciation events. The 1R event occurred within 0-34 million years before the divergence of crown vertebrates. Species which underwent the 1R event but not the subsequent 2R event, i.e. cyclostomes, are not morphologically diverse even though they have experienced their own WGD events and their lineage is of equal longevity.

The timing of the 2R event indicates that it occurred very early within the gnathostome stem-lineage. This leaves a considerable time lag between the 2R WGD event and the divergence of crown gnathostomes. Such extensive lag periods have also been observed in salmonids, in both the seed plant and flowering plant WGD events, and marshalled in evidence against a causal relationship between WGD and macroevolutionary outcomes[171,172]. However, the 2R duplication event also appears to give rise to the largest increase in morphospace occupation in vertebrates, with the descendants of the event representing the vast majority of vertebrate disparity. Furthermore, the timing of the 2R event at the base of the gnathostome stem-lineage indicates that it occurred before the sequential addition of gnathostome characters revealed by fossil stem-gnathostomes[175,184]. Our lower estimate for the age of the 2R event is older than the earliest unequivocal ostracoderm fossils[184–186]. This indicates that all ostracoderms are descendants of the 2R event. Consequently, the emergence of gnathostome disparity was neither explosive nor delayed. Rather, phenotypic characters were acquired along the gnathostome stem during the protracted period between WGD and crown-gnathostome divergence. The timing of the 2R event, early within the gnathostome stem-lineage, also demonstrates that WGDs do not appear to confer extinction resistance in any material sense[187]. This observation is particularly significant since the link between WGD and extinction resistance was based in large part on the predicted relationship between the 2R event and the extensive gnathostome stem-lineage diversity[187].

While both crown-vertebrate lineages (cyclostome and gnathostome) that descended from the 1R event are of equal antiquity and underwent subsequent, independent, WGD events (2R, CR1, CR2), there is a stark imbalance in both species diversity and phenotypic disparity. The CR1, CR2 and 2R events occurred early in the history of their respective lineages. However, unlike the 2R event, the CR1 and CR2 events are not associated with a substantive increased disparity and diversity. Thus, the evolutionary consequences of WGD are not equal, suggesting that 2R in itself is not sufficient to explain phenotypic innovation in early vertebrate evolution. Rather, it is likely the specific conspiracy of circumstances between genetic potential and ecological opportunity, facilitated by developmental and phenotypic innovation, that underpins the differences in evolutionary potential between clades like cyclostomes and gnathostomes (cf. REF. 188).

#### SUPPLEMENTARY FILES

**Supplementary File 1 -** Hi-C maps of all cases of misassembly in version 3.2 before manual correction (related to Supplementary Figure 2).

**Supplementary File 2 -** Multifasta file with all hagfish vision-related genes.

**Supplementary File 3 -** Excel file containing Datasets 1-3 with relational information about orthologs of different species.

- Dataset 1: Between chicken and spotted gar, between chicken and elephant shark, between spotted gar and elephant shark.
- Dataset 2: Between hagfish and lamprey
- Dataset 3: Between vertebrates (chicken, spotted gar, elephant shark, hagfish and sea lampery) and invertebrates (sea cucumber and Belcher's lancelet)

**Supplementary File 4 -** Multifasta file with 1,957 genes used in phylotenetic analyses supporting 1R/2R (132 in sea cucumber, 132 in Belcher's lancelet, 24 in Florida lancelet, 363 in hagfish, 703 in sea lamprey, 94 in elephant shark, 316 in chicken, 323 in spotted gar). Gene alignments were visually inspected and manually curated if necessary as described in Supplementary Information. Related to Figure 4a.

**Supplementary File 5 -** All 193 gene trees describing the phylogenetic position of elephant shark *C. milii* genes in chromosomal trees. Related to Figure 4a.

**Supplementary File 6 -** All 190 gene trees describing the phylogenetic position of inshore hagfish *E. burgeri* genes in chromosomal trees. Related to Figure 4a.

**Supplementary File 7 -** All 172 gene trees describing the phylogenetic position of the sea lamprey *P. marinus* genes in chromosomal trees. Related to Figure 4a.

**Supplementary File 8 -** All 115 gene trees describing the phylogenetic position of both inshore hagfish and sea lamprey genes in chromosomal trees. Related to Figure 4a.

**Supplementary File 9 -** Overlapping ratios between hagfish chromosomes (left panels) and sea lamprey scaffolds (right panels) deriving from the same AC. Related to Extended Data Figure 7a,b. Mutually (best-to-best) paralogous chromosomes are listed at the bottom of each panel.

**Supplementary File 10 -** Clustering analysis of duplicate retention profiles of orthologous chromosomes in chicken and spotted gar. Related to Extended Data Figure 7c.

**Supplementary File 11 -** Clustering analysis of duplicate retention profiles of putative orthologous chromosomes of the inshore hagfish and sea lamprey. Related to Figure 4g.

**Supplementary File 12 -** R script used for transcriptomics data normalization. Related to Figures 5f,g, and Extended Data Figure 8g-j.

**Supplementary File 13 -** Compressed folder containing the alignment used in the phylogenomics tree and the resulting tree.

**Supplementary File 14 -** Compressed folder containing the alignments, control files and tree used for dating species divergence (see Methods).

**Supplementary File 15 -** Orthogroups obtained by OrthoFinder.

**Supplementary File 16 -** Multiple sequence alignment and MrBayes options and parameter values of Agap genes, in Nexus format. Related to Extended Data Figure 4d.

**Supplementary File 17 -** Multiple sequence alignment and MrBayes options and parameter values of Gbx genes, in Nexus format. Related to Extended Data Figure 4a.

**Supplementary File 18 -** Multiple sequence alignment and MrBayes options and parameter values of Cbx genes, in Nexus format. Related to Extended Data Figure 4b.

**Supplementary File 19 -** Multiple sequence alignment and MrBayes options and parameter values of Hnrnpa genes, in Nexus format. Related to Extended Data Figure 4c.

**Supplementary File 20 -** Compressed folder containing alignments, MCMCtree control files and calibrations used for dating genome duplications in vertebrates.

**Supplementary File 21 -** Character matrix and descriptions (Vertebrate_disparity_matrix.nex) and tree with estimated ancestral character states (Disparity.tre) used for the phenotypic disparity analyses. Related to Figure 5g,h.

#### SUPPLEMENTARY TABLES

**Supplementary Table 1 -** Sources of genomic and annotation data used.

**Supplementary Table 2 -** Sequencing libraries and basic statistics of clean sequencing data, after filtering, and processing steps they were used for.

**Supplementary Table 3 -** Statistics of reads before and after error correction of data from short-insert libraries.

**Supplementary Table 4 -** Stats of data used for genome size estimation based on 17-mer distribution.

**Supplementary Table 5 -** Dovetail Genomics's HiRise performance and library stats.

**Supplementary Table 6 -** Basic statistics of different genome assembly versions.

**Supplementary Table 7 -** Misassembled scaffolds in the hagfish Dovetail assembly (v3.2). 280 misassembled scaffolds involving 364 misjoining boundaries are listed here.

**Supplementary Table 8 -** After correction, hagfish scaffolds were regrouped as 19 clusters based on HiC contact information. From top to bottom, the rank of scaffolds denotes their relative orders in one cluster (pseudo-chromosome). Each scaffold is either in sense (orientation = 0) or antisense (orientation = 1) orientation in one cluster.

**Supplementary Table 9 -** All hagfish clusters have similar read coverage from DNA-seq reads.

**Supplementary Table 10 -** 41 hagfish genes were found at misassembly boundaries and removed from further analyses.

**Supplementary Table 11 -** Transcript structures identified from short read RNA-seq data. The table shows the number of unique transcript structures identified in each tissue prior to downstream filtering and QC.

**Supplementary Table 12 -**Overall GC-content percentages at genomic and coding sequence (CDS) levels of selected chordates.

**Supplementary Table 13 -** LastZ parameters used to align hagfish to lamprey, zebrafish, and human.

**Supplementary Table 14 -** Comparison of the coverage of the hagfish and lamprey genesets (coding and non-coding) by the gene-trees analysis. Ensembl v93 annotations were used.

**Supplementary Table 15 -** Number of orthologues, sequence identity, and number of high-confidence calls for hagfish orthologies against lamprey, zebrafish, and human.

**Supplementary Table 16 -** Number of orthologues, sequence identity, and number of high-confidence calls for lamprey orthologies against zebrafish and human.

**Supplementary Table 17 -** Number of genes that are structurally different from their orthologues in other species.

**Supplementary Table 18 -** A size summary of coding exons from 11 species.

**Supplementary Table 19 -** A size summary of introns from 11 species.

**Supplementary Table 20 -** A size summary of coding transcript from 11 species.

**Supplementary Table 21 -** A size summary of genes from 11 species.

**Supplementary Table 22 -** A summary of intergenic regions from 11 species.

**Supplementary Table 23 -** GO enrichment analysis of gene families present in the reconstructed ancestral vertebrate genome.

**Supplementary Table 24 -** Number of Homology groups (HG) gained at different notes (Novel) and their Panther GO enrichment analysis.

**Supplementary Table 25 -** Panther GO enrichment analysis of vertebrate Novel Core HGs.

**Supplementary Table 26 -** Panther GO enrichment analysis of gnathostome Novel Core HGs.

**Supplementary Table 27 -** Immune-related genes in the current version of hagfish genome.

**Supplementary Table 28 -** List of circadian clock and clock-controlled genes found in the genome of Eptatretus burgeri and their Ensembl IDs.

**Supplementary Table 29 -** Annotation information of Hox genes and their linked non-Hox genes in the genome of E. burgeri.

**Supplementary Table 30 -** Nomenclature update of Hox genes from Pascual-Anaya et al., 2018 (REF. 19) and this study after orthology assignment to lamprey Hox clusters.

**Supplementary Table 31 -** Numbers of best-to-best genes between back-mutated gar chromosomes and back-mutated chicken chromosomes.

**Supplementary Table 32 -** Homology between sea cucumber chromosomes, ancestral chromosomes (ACs), chicken chromosomes and gar chromosomes.

**Supplementary Table 33 -** 16 ACs reconstructed on the basis of amphioxus genes.

**Supplementary Table 34 -** The correspondance between 17 ACs, 17 tetrads, 17 CLGs and 18 PVCs.

**Supplementary Table 35 -** Numbers of best-to-best genes between ACs and Florida lancelet chromosomes.

**Supplementary Table 36 -** Homologous gene sets underlying chromosomal phylogenetic analyses of chicken and gar.

**Supplementary Table 37 -** Comparison of post-1R chromosomal fusion events identified in this study with those identified in three previous studies.

**Supplementary Table 38 -** Frequency distribution of "pairwise post-1R fusion" based on random sampling.

**Supplementary Table 39 -** Homologous gene sets used to infer the position of cyclostome genes in single gene phylogenies.

**Supplementary Table 40 -** The overlapping ratio of two parts of a chromosome in chicken, spotted gar, hagfish, or sea lamprey genome after artificial splitting.

**Supplementary Table 41 -** Overlapping ratio of orthologous chromosome pairs of chicken and gar.

**Supplementary Table 42 -** Overlapping ratio of orthologous chromosome pairs in hagfish and lamprey.

**Supplementary Table 43 -** The overlapping ratio of paralogous chromosomes in chicken or spotted gar genome that correspond to one same AC.

**Supplementary Table 44 -** The overlapping ratio of putative paralogous chromosomes in hagfish or sea lamprey genome that correspond to one same AC.

**Supplementary Table 45 -** Merging Sea lamprey scaffolds.

**Supplementary Table 46 -** Number of ACRs in promoters.

**Supplementary Table 47 -** Statistic information for Figure 5b (ACRs in promoters/proximal regions).

**Supplementary Table 48 -** Statistic information for Figure 5b (ACRs in distal regions).

**Supplementary Table 49 -** Statistic information for Figure 5c.

**Supplementary Table 50 -** Statistic information for Figure 5a.

**Supplementary Table 51 -** Sources of ATAC-seq data amphioxus, hagfish, zebrafish, medaka, chicken and mouse used for regulatory genome profilings.

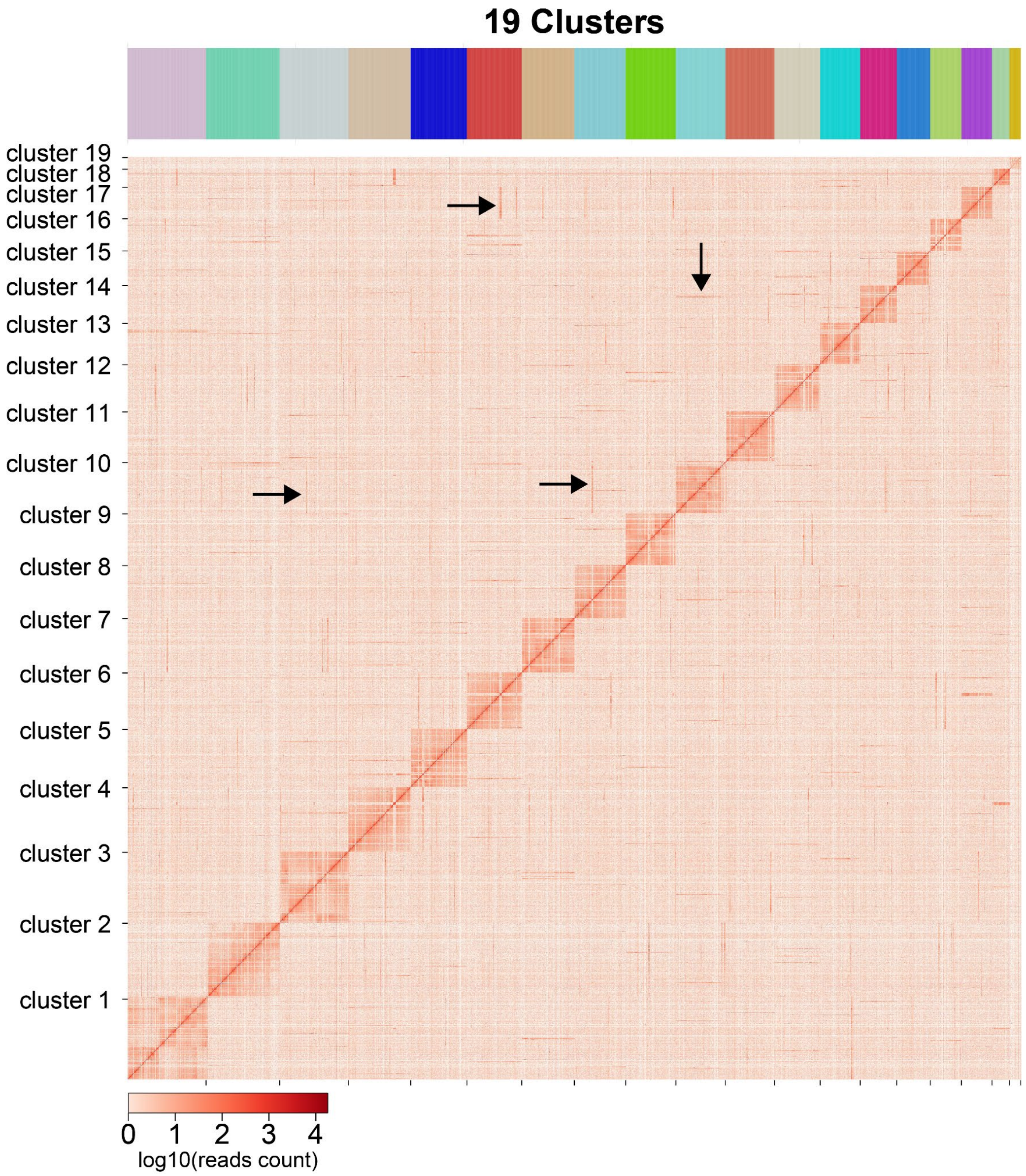


**Supplementary Figure 1. Preliminary 19-cluster Hi-C heatmap of the chromosome-scale genome assembly of *E. burgeri*.** Numerous off-diagonal long stretches of Hi-C contacts are noticeable, and a few examples are marked with black arrows. They may represent misassembled scaffolds.

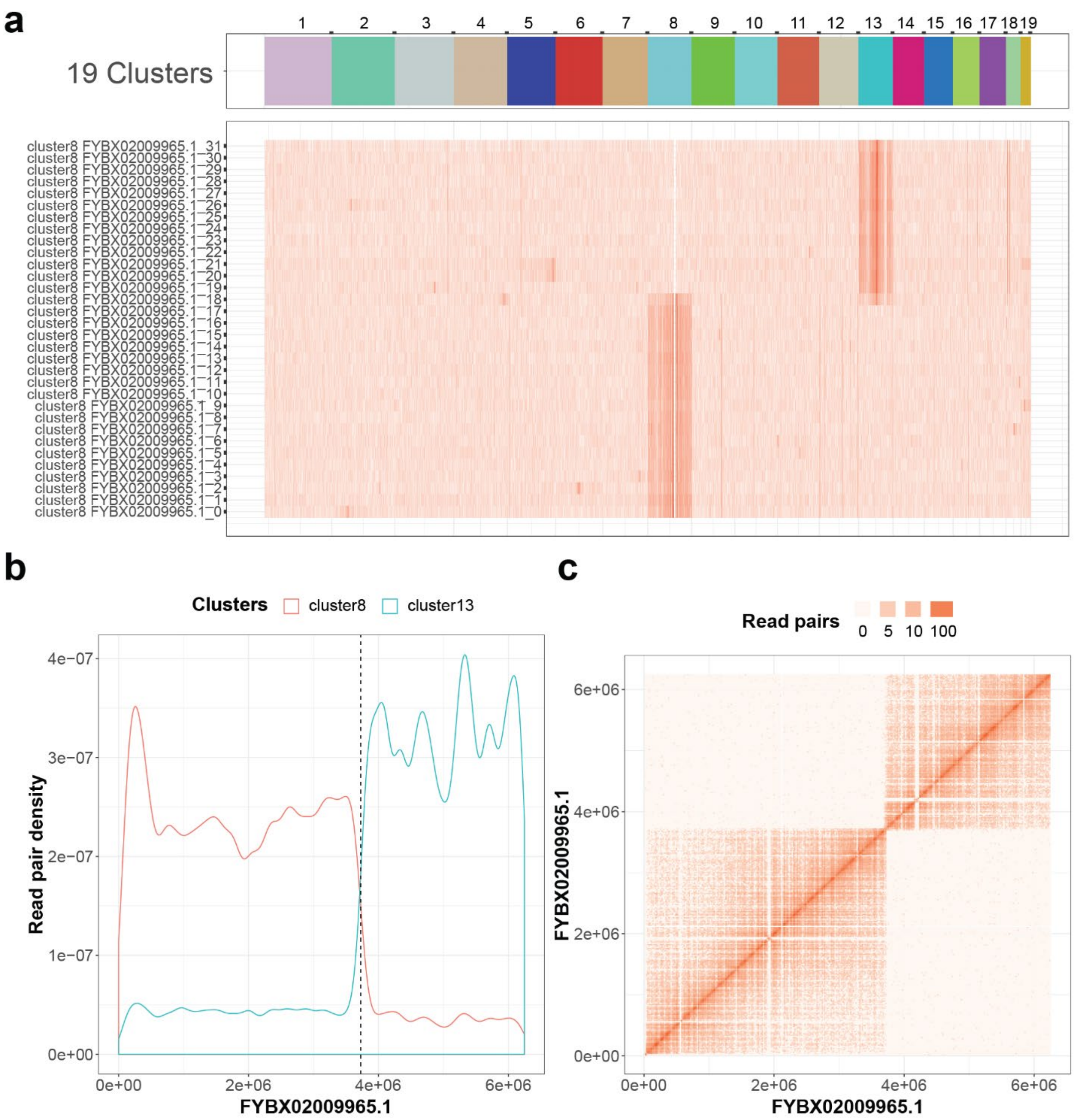


**Supplementary Figure 2. Example of a misassembly in the genome assembly version 3.2 detected with three types of Hi-C contact signals. a,** A scaffold named FYBX02009965.1 (harboring 32 200-kb segments, 0~31) is grouped into cluster8 but has abundant Hi-C contacts with cluster13 in its last thirteen segments, 19~31. **b,** The distribution of Hi-C contacts in FYBX02009965.1 relative to cluster8 or cluster13 could be divided into two partitions where the first 3.8 Mb (mega base) mainly interacts with cluster8 and the remaining 2.6 Mb mainly interacts with cluster 13. A dashed line denotes the boundary. **c,** Within-scaffold Hi-C contacts of FYBX02009965.1 clearly shows two separate blocks. The consistency across all three panels in terms of partition boundary suggests that FYBX02009965.1 represents a misassembly of cluster8 (3.8 Mb) and cluster13 (2.6 Mb) derived segments.

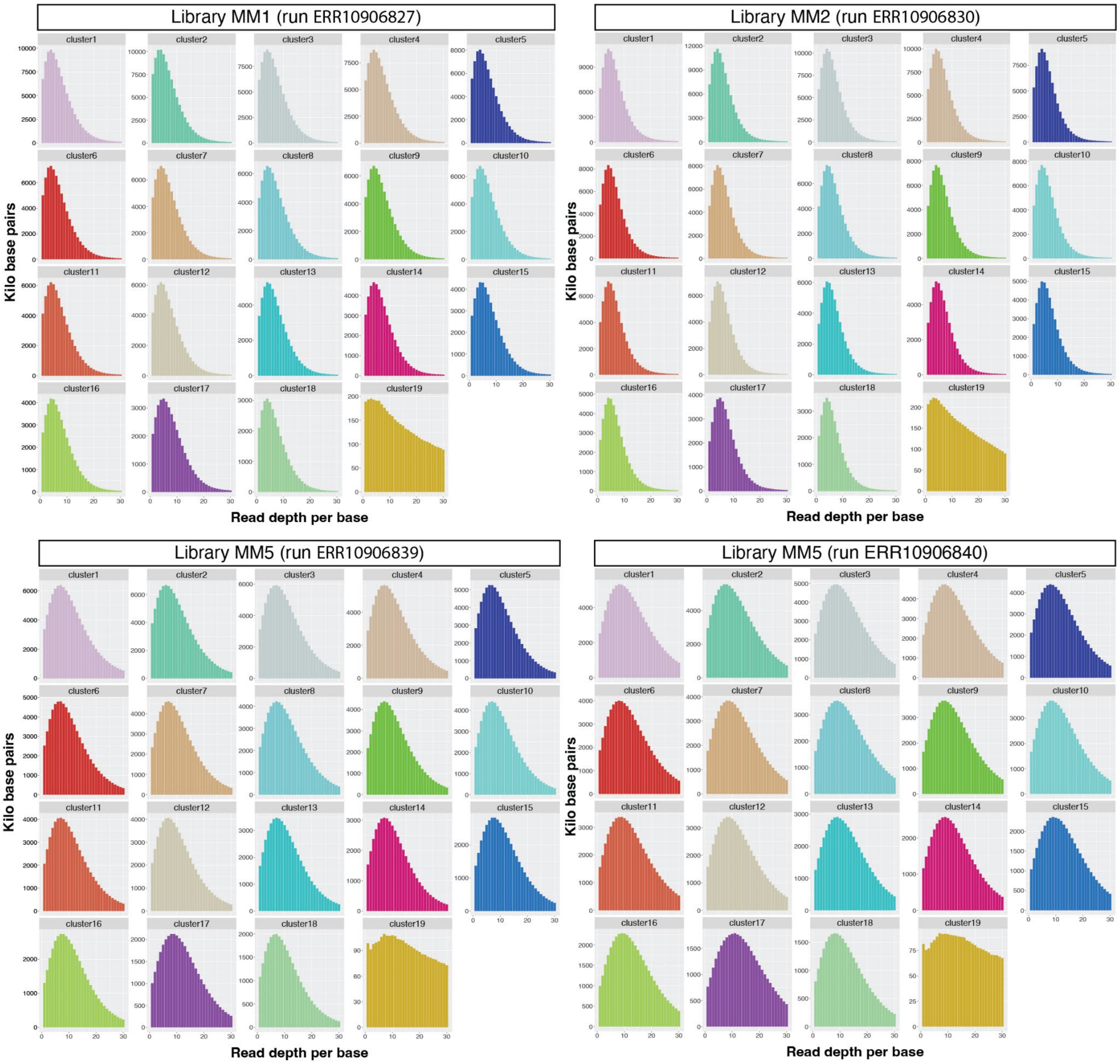


**Supplementary Figure 3. DNA sequencing read depth distribution across all 19 contact clusters of the E. burgeri genome Hi-C assembly.** Read depth per kilobases calculated using 4 randomly chosen Illumina lanes are plotted. For simplicity, read depths higher than 30 are not shown. Peak values per each run and cluster can be found in Supplementary Table 8.

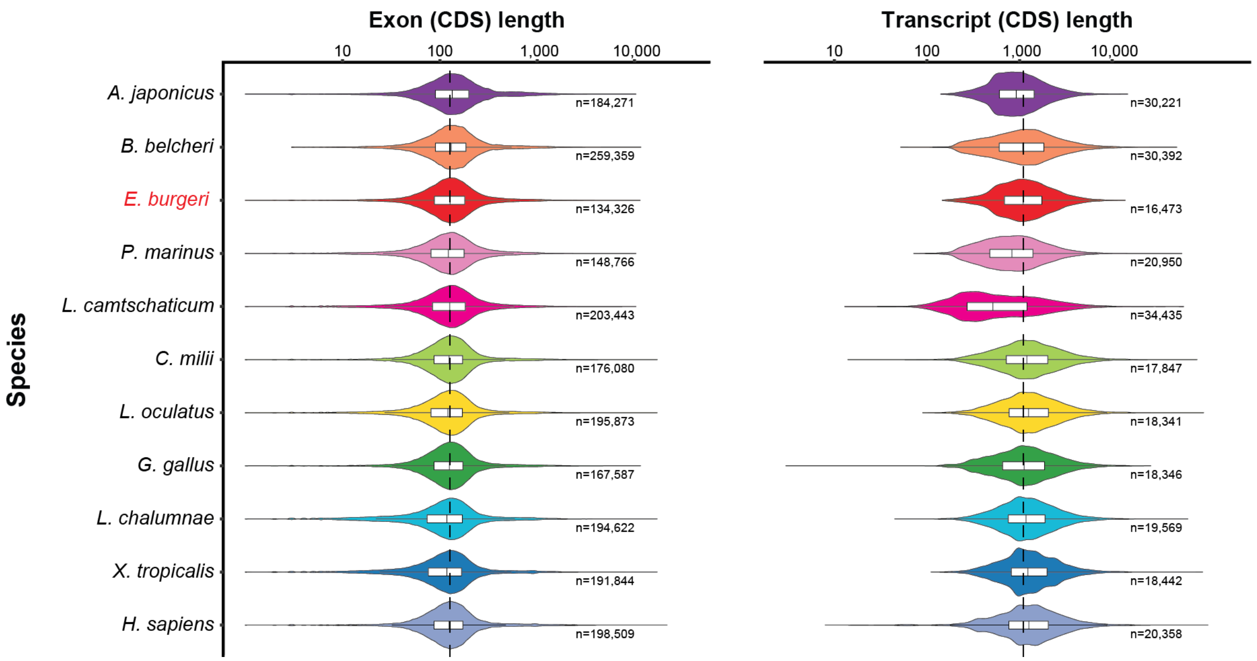


**Supplementary Figure 4. Exon and transcript lengths of vertebrate genes.** Violin plots of size distribution of exon and transcript lengths over a logarithmic scale of all annotated genes of the hagfish (*E. burgeri*), two lamprey species (sea lamprey, *P. marinus*; Arctic lamprey, *L. camtschaticum*), six gnathostome vertebrates (human, *H. sapiens*; frog, *X. tropicalis*; coelacanth, *L. chalumnae*; chicken, *G. gallus*; spotted gar, *L. oculatus*; and the elephant shark, *C. milii*) and two invertebrate deuterostomes (sea cucumber, *A. japonicus*; amphioxus, *B. belcheri*). For each genomic feature and each species, the median and IQR (interquartile range) length statistics are indicated with a white rectangle. Dashed vertical line indicates median size of *E. burgeri* features.

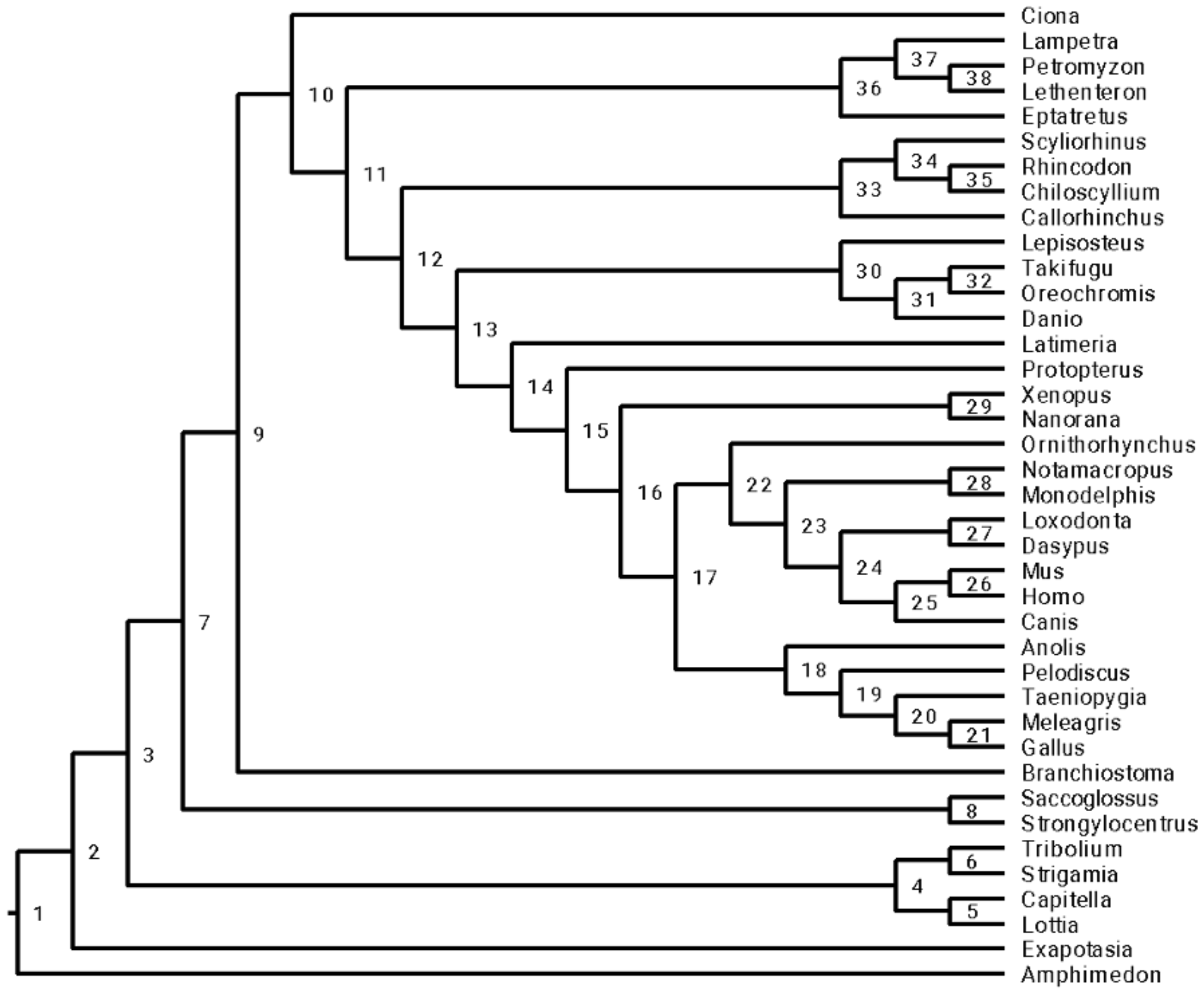


**Supplementary Figure 5. Node calibrations.** Node labels are used to indicate the node being calibrated (see section 2.2).

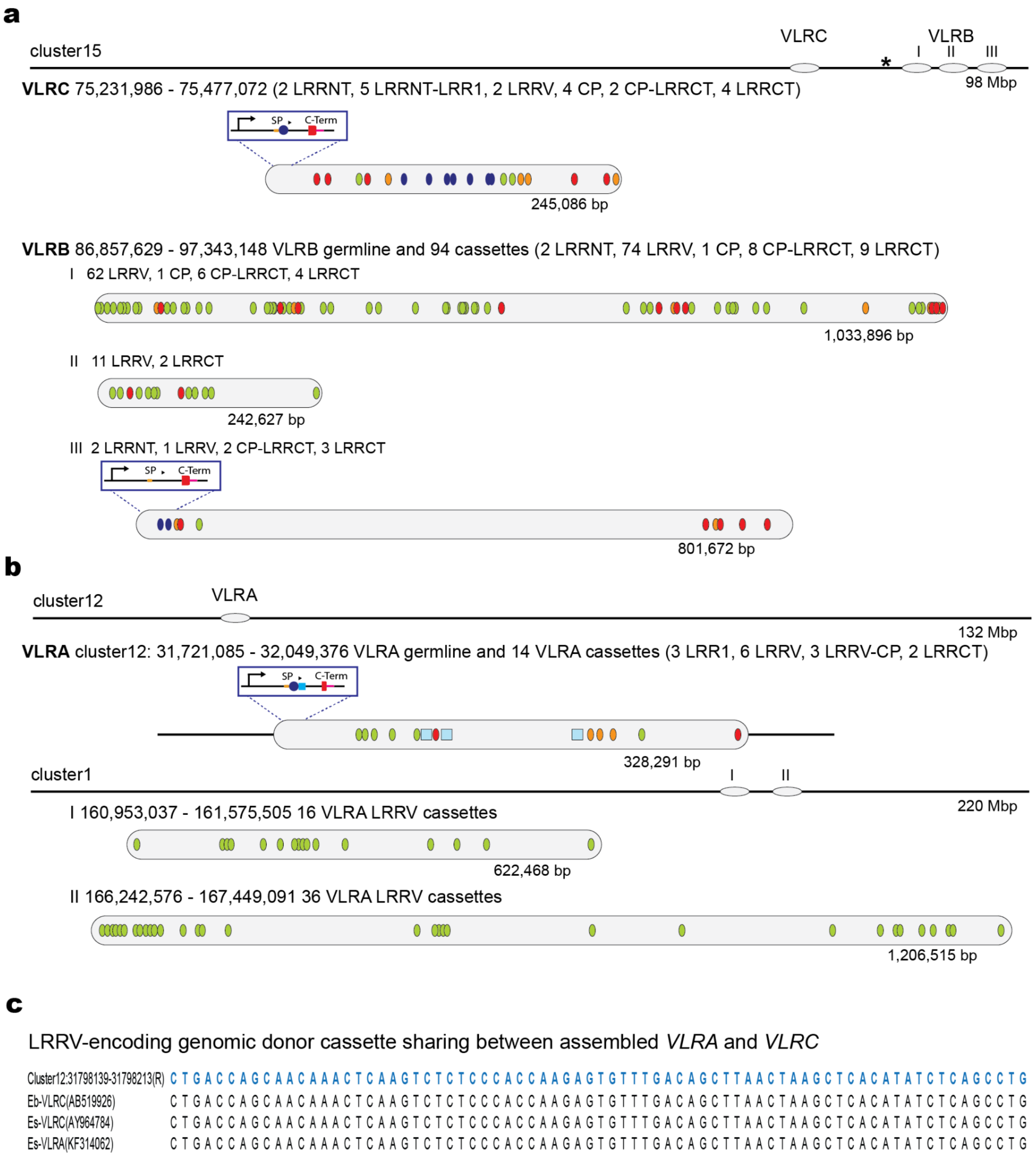


**Supplementary Figure 6. *VLR* loci organisation in the hagfish. a**, The genomic organisation of *E. burgeri VLRB* and *VLRC*. The germline *VLR* genes are located near clusters of donor cassettes. A map of assembly cluster15 is shown at the top. *VLRB* donor cassettes are encoded in three subclusters designated I, II, III and shown as light blue oval. The *VLRC* and *VLRB* germline genes are located approximately 21 Mbp apart with 12.5 Mbp separating their associated cassette clusters. Details of the cassette organisation within clusters are shown in the larger grey ovals. Cluster sizes are given under the grey oval diagrams. Cassette icons are spaced proportionally to positions in the assembly sequence but are not drawn to scale. *VLR* germline gene structures are shown in boxes above the detailed cassette cluster representations. Cassette-types are illustrated with colored icons: dark blue ovals, *LRRNT* or *LRRNT-LRR1*; green ovals, *LRRV*; orange ovals, connecting peptide (*CP*); red ovals, *LRRCT*. For germline gene diagrams: *SP*, signal peptide; *C-*Term, invariant C-terminal region; dark blue circle, germline LRRNT; light blue square, LRR1. Four isolated *VLRB* donor cassettes located upstream of *VLRB* subcluster I are indicated by an asterisk at the top. **b**, *VLRA* is encoded on assembly cluster12 with two subclusters of *VLRA*-like *LRRV* cassettes located on assembly cluster1. Colors and icons are as in (A) with the addition of light blue squares representing individual *VLRA LRR1* cassettes. **c**, Examples of genomic donor cassette sharing between assembled *VLRA* and *VLRC* in hagfish. The genomic donor cassette sequence from the *E. burgeri* genome is shown in blue. The position of this cassette in the genome and the accession number of the mature *VLRA* and *VLRC* sequences are indicated. NB: Since only mature *VLRC* sequence (previously identified as *VLRA*) from *E. burgeri* is available in NCBI, we included *VLRA* (the third VLR) and *VLRC* from *Eptatretus stoutii*.

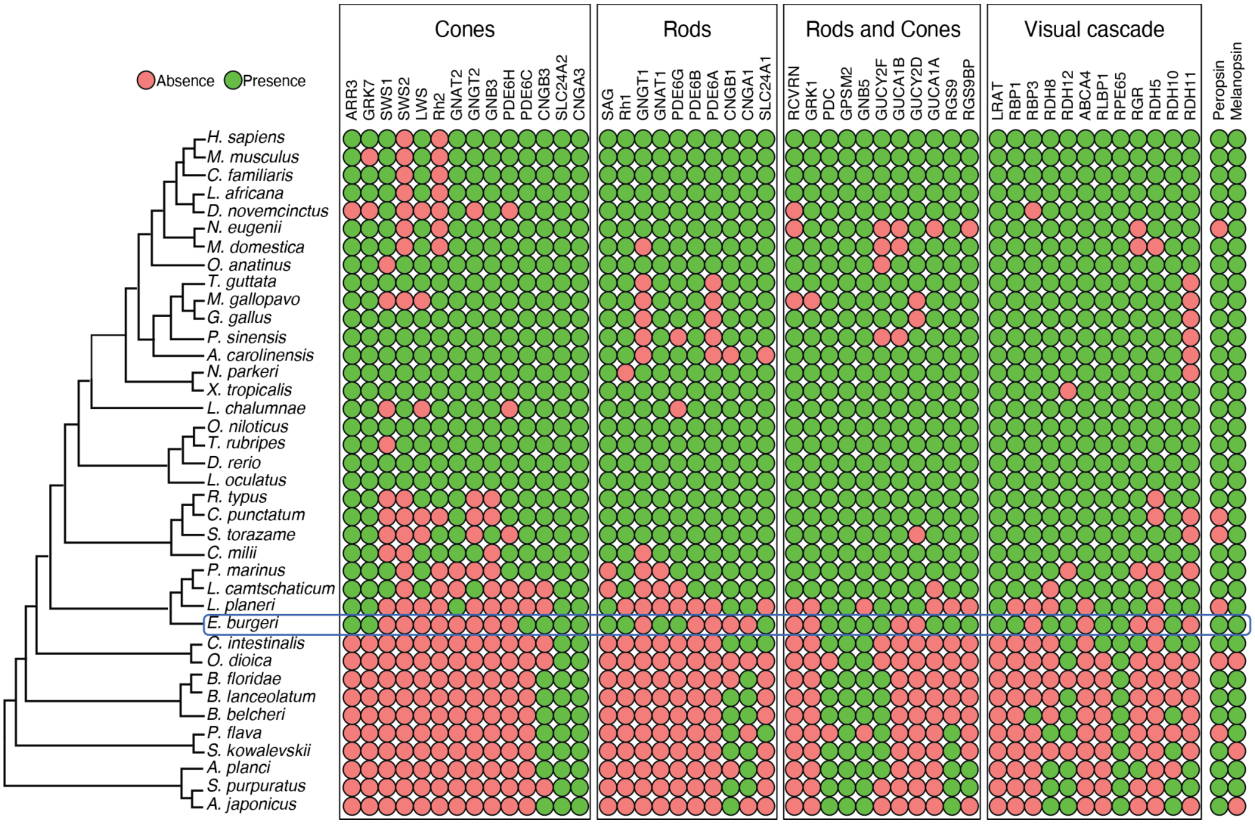


**Supplementary Figure 7. Evolution of visual genes deuterostomes.** Phylogeny of deuterostomes (left) with the presence (green) and absence (red) of vision-associated genes. Hagfish genome possess three opsin genes: Melanopsin (Opn4), Peropsin (RRH) and Rhodopsin 1 (RH1 or RHO), but have secondarily lost most cone-specific genes, as well as several rod-specific genes. Species and the taxa they belong to can be found in Supplementary Table 1. A fasta file with *E. burgeri* gene sequences is provided in Supplementary File 2.

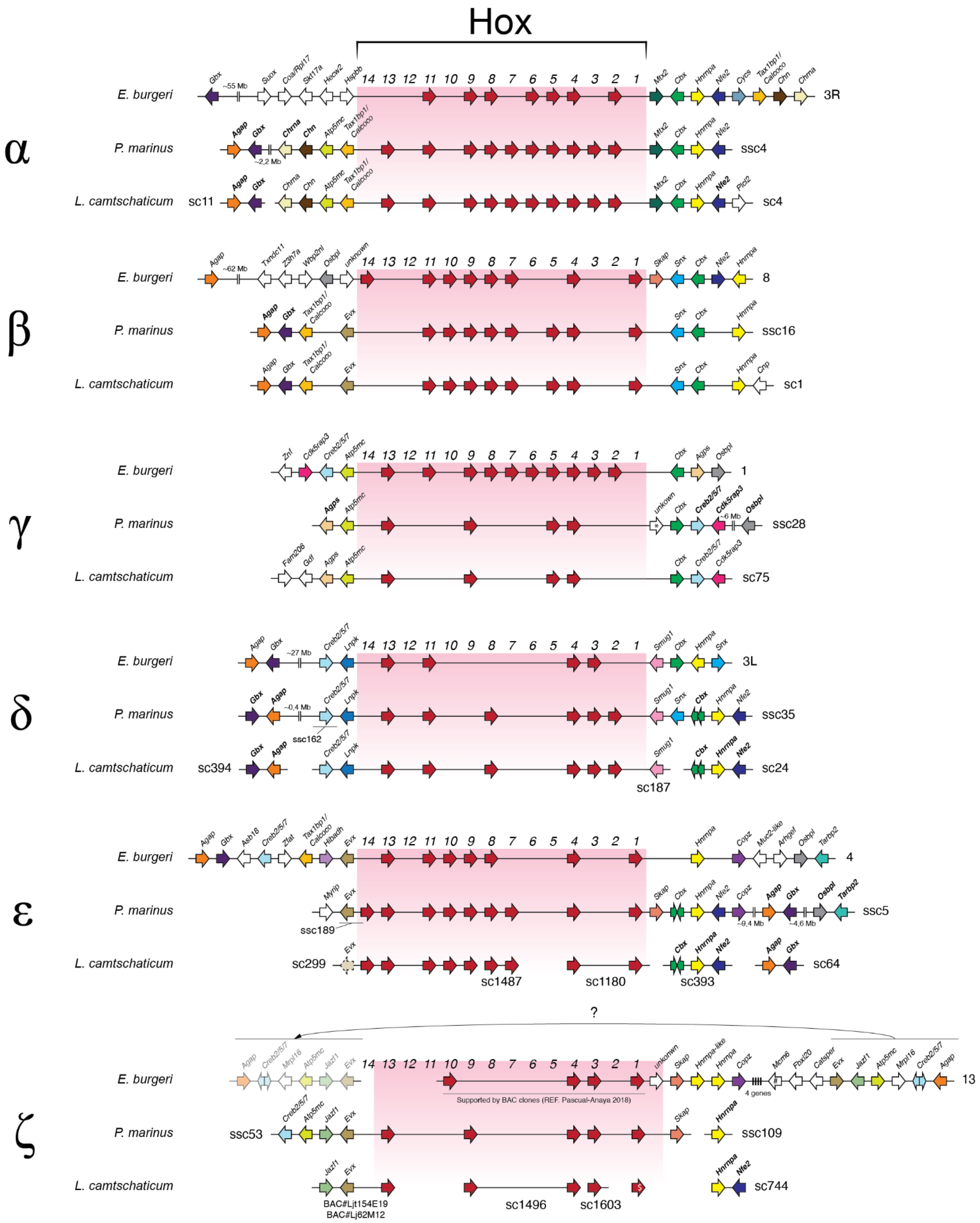
Hox
14 13 12 11 10 9 8 7 6 5 4 3 2 1
α
E. burgeri
Gbx ~55 Mb Suox Coa/Rpl17 Skt17a Hecw2 Hspbb
Mtx2 Cbx Hnrnpa Nfe2 Cyrs Tax1bp1/Calcoco Chn Chrna
3R
P. marinus
Agap Gbx ~2,2 Mb Chrna Chn Atp5mc Tax1bp1/Calcoco
Mtx2 Cbx Hnrnpa Nfe2
ssc4
L. camtschaticum
sc11
Agap Gbx Chrna Chn Atp5mc Tax1bp1/Calcoco
Mtx2 Cbx Hnrnpa Nfe2 Plcl2
sc4
β
E. burgeri
Agap ~62 Mb Txndc11 Z3h7a Wbp2nl Osbpl unknown
Skap Snx Cbx Nfe2 Hnrnpa
8
P. marinus
Agap Gbx Tax1bp1/Calcoco Evx
Snx Cbx Hnrnpa
ssc16
L. camtschaticum
Agap Gbx Tax1bp1/Calcoco Evx
Snx Cbx Hnrnpa Cnp
sc1
γ
E. burgeri
Znf Cdk5rap3 Creb2/5/7 Atp5mc
Cbx Agps Osbpl
1
P. marinus
Agps Atp5mc
unkown Cbx Creb2/5/7 Cdk5rap3 ~6 Mb Osbpl
ssc28
L. camtschaticum
Fam206 Gdf Agps Atp5mc
Cbx Creb2/5/7 Cdk5rap3
sc75
δ
E. burgeri
Agap Gbx ~27 Mb Creb2/5/7 Lnpk
Smug1 Cbx Hnrnpa Snx
3L
P. marinus
Gbx Agap ~0,4 Mb Creb2/5/7 Lnpk
ssc162
Smug1 Snx Cbx Hnrnpa Nfe2
ssc35
L. camtschaticum
sc394
Gbx Agap Creb2/5/7 Lnpk
Smug1
sc187
Cbx Hnrnpa Nfe2
sc24
ε
E. burgeri
Agap Gbx Asb18 Creb2/5/7 Zfat Tax1bp1/Calcoco Hibadh Evx
Hnrnpa Copz Muc2-like Arhgef Osbpl Tarbp2
4
P. marinus
Myrip Evx
ssc189
Skap Cbx Hnrnpa Nfe2 Copz ~9,4 Mb Agap Gbx ~4,6 Mb Osbpl Tarbp2
ssc5
L. camtschaticum
sc299
Evx
sc1487
sc1180
Cbx Hnrnpa Nfe2
sc393
Agap Gbx
sc64
?
ζ
E. burgeri
Agap Creb2/5/7 Mrpl16 Atp5mc Jazf1 Evx
Supported by BAC clones (REF. Pascual-Anaya 2018)
unknown Skap Hnrnpa-like Hnrnpa Copz 4 genes Mcm6 Fbxl20 Catsper Evx Jazf1 Atp5mc Mrpl16 Creb2/5/7 Agap
13
P. marinus
ssc53
Creb2/5/7 Atp5mc Jazf1 Evx
Skap
Hnrnpa
ssc109
L. camtschaticum
Jazf1 Evx
BAC#Ljt154E19
BAC#Lj62M12
sc1496
sc1603
Hnrnpa Nfe2
sc744

2175

**Supplementary Figure 8. Orthology relationships of cyclostome Hox clusters.** Orthologous *Hox* clusters of the inshore hagfish and two lamprey species (sea lamprey, *P. marinus*[141], and Arctic lamprey, *L. camstchaticum*[189]) are shown by cluster orthology. Hagfish *Hox* clusters have been assigned to each orthology group based on (i) microsynteny conservation of both Hox gene composition of each cluster and syntenic non-Hox genes, (ii) phylogenetic analyses of candidate genes (*Gbx*, *Agap*, *Hnrnpa* and *Cbx*; Extended Data Figure 4a-d), and (iii) clustering analysis of retention profiles (Extended Data Figure 4e). (i) and (ii) are sufficient to assign orthology to clusters α, γ, δ, and ζ. Assignment to Hox clusters β and ε is resolved after (iii). All genes are represented by color coded arrows according to their homology, with Hox genes in red, and whose direction indicates its sense of transcription. Separate scaffolds harbouring *Hnrnpa*, *Agap*, *Gbx* or *Cbx* genes are assigned to a particular clusters based on phylogenetic analyses. The block encompassing genes from Evx-ζ to Agap-ζ downstream to its syntenic Hox cluster might be the result of missassembly. Syntenic non-Hox genes with bold font are found in this study. The assembly scaffold or chromosome where they are located in their respective assemblies are indicated to the right of each cluster (ssc, superscaffold of the sea lamprey 2018 assembly; sc, scaffold of the 2013 Arctic lamprey assembly). #, Mcmc6 gene is also present in amphioxus, implying its possible ancestral link to a Hox cluster in chordates; *, PMZ_0048273 in REF. [141]; $, Described in REF. 25.

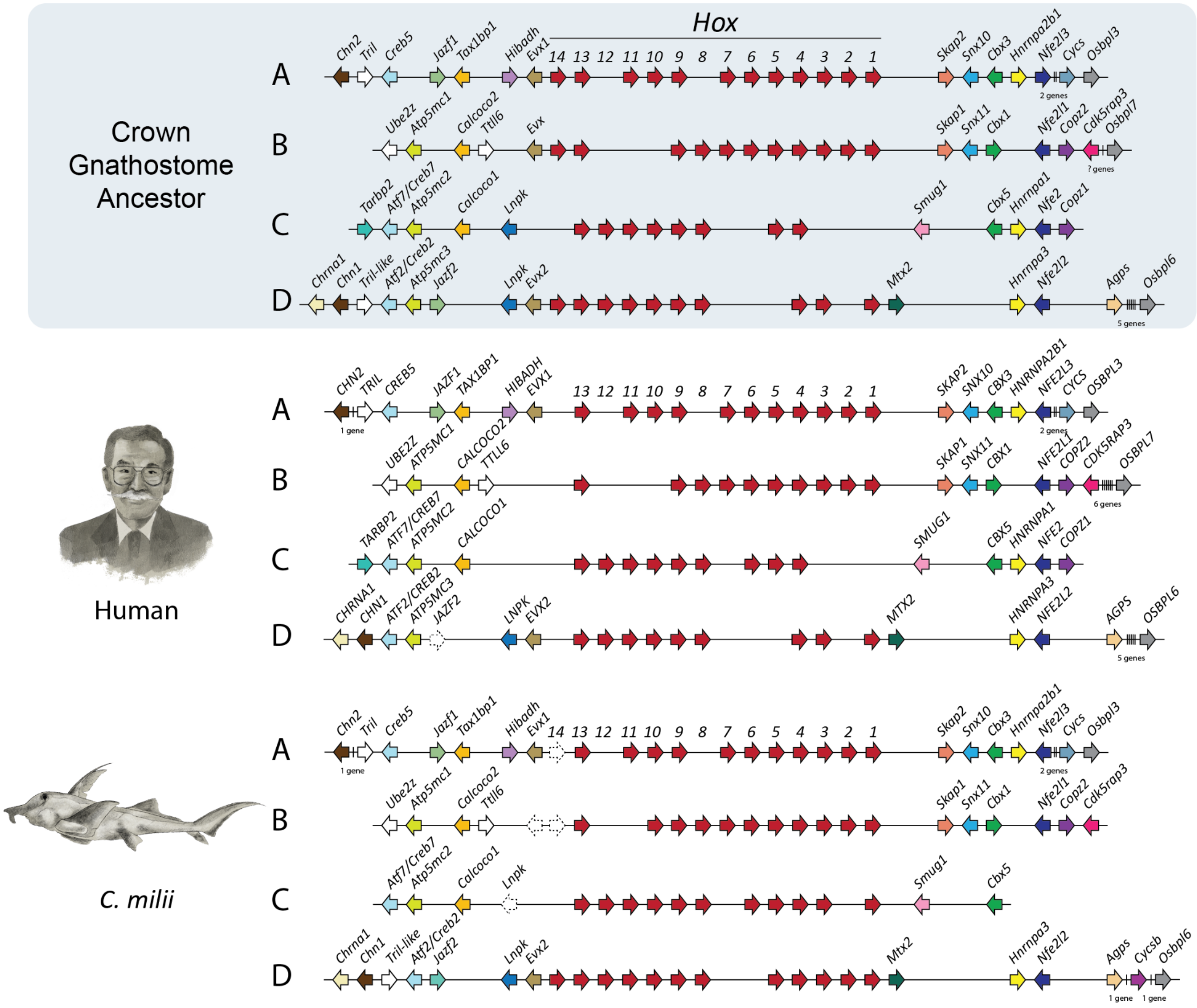


**Supplementary Figure 9. Gnathostome Hox clusters.** Schematic representations of Hox clusters and syntenic genes of the elephant shark (*C. milii*; bottom), human (middle), and a reconstruction of the complement of the last common ancestor of gnathostomes (top). Genes are represented by coloured-coded arrows, whose direction marks the sense of transcription: Hox genes in red, non-Hox genes coloured by homology. Information about syntenic genes was based on Acemel et al.[152] and manual inspection at Ensembl genome browser.

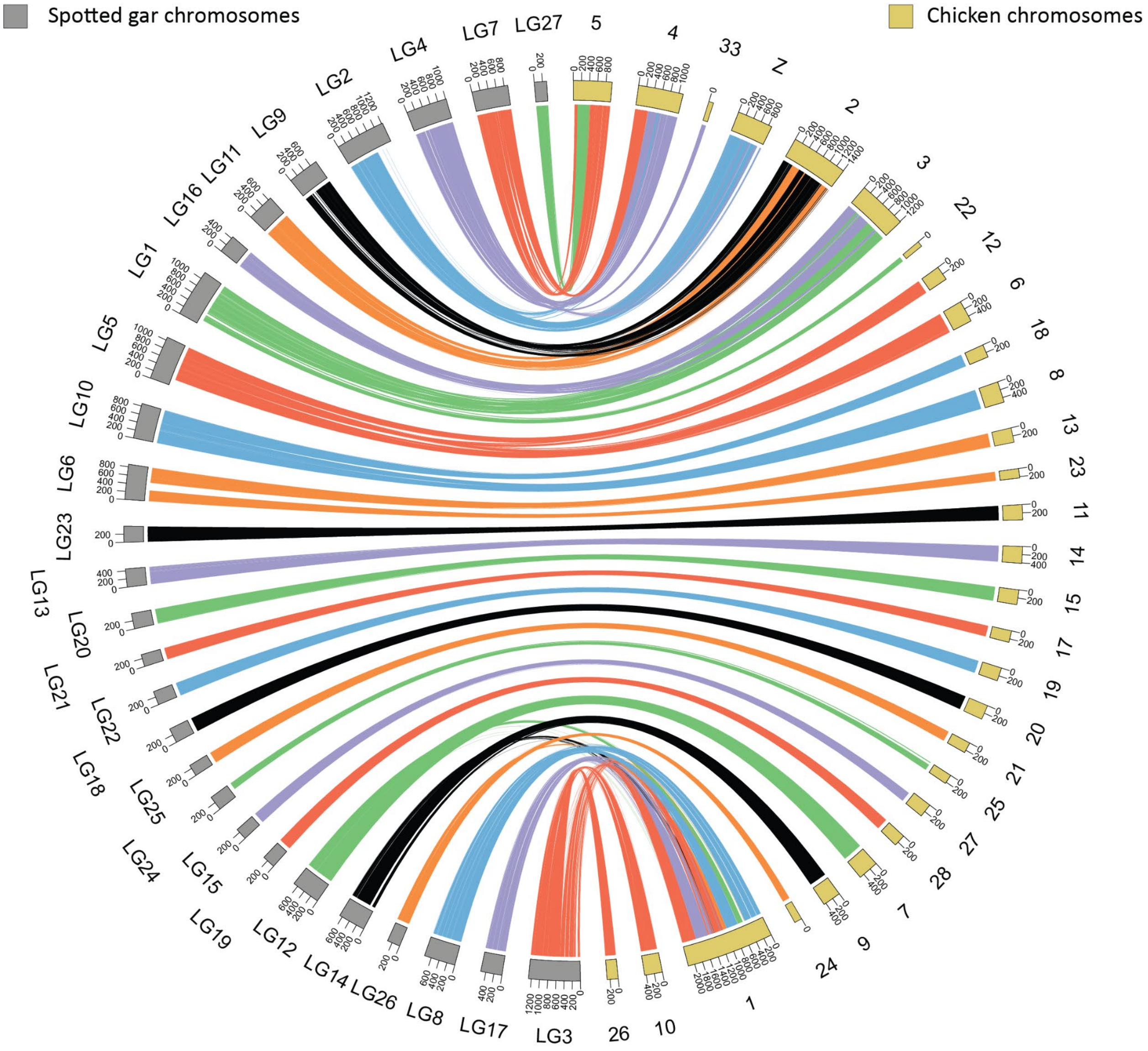


**Supplementary Figure 10. Chromosomal correspondence between chicken and gar.** A circus plot is used to show macrosynteny conservation. Each chromosome is scaled to the number of genes it contains. Chromosomes are arranged in a way to better visualise the correspondence between species. Each line represents one pair of homologous genes between chromosomes, which is colour-coded according to different spotted gar chromosomes and from a set of six colors (red/purple/blue/black/orage/green).

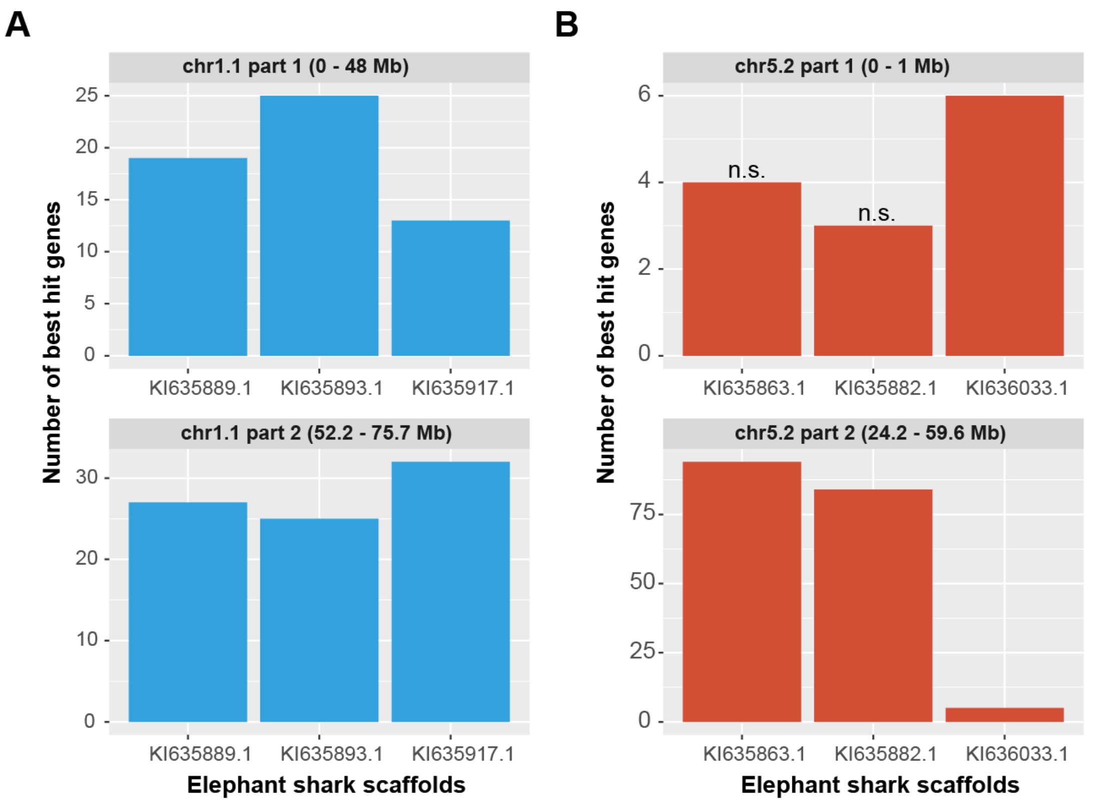


**Supplementary Figure 11. Two chicken chromosomal sections, chr1.1 or chr5.2, are homologous to the same set of shark scaffolds.** All correspondence relationships of chr1.1 (**a**) and chr5.2 (**b**) are statistically significant except that the homology of chr5.2 part 1 with elephant shark KI635863.1 or KI35882.1 is not (marked by 'n.s.'). We used 'homologous gene pair number > 5 and FDR q value < 0.05' as the cutoff to define homologous relationships. Since this segment does not harbour many genes (79 for chr5.2 part 1), only 4 and 3 homologous genes were identified in KI635863.1 and KI35882.1, respectively. We thus considered them as non-significant homologous relationships.

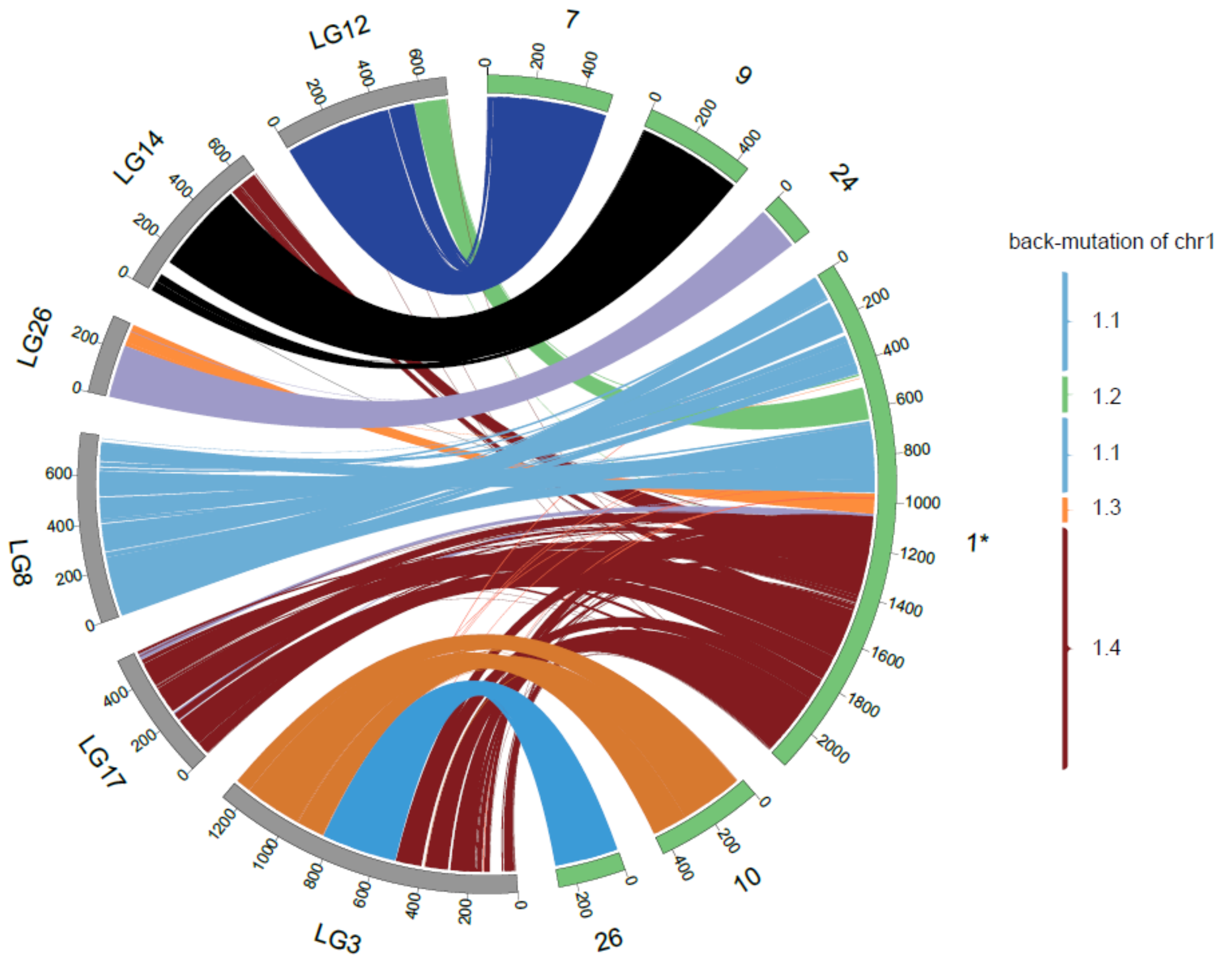


**Supplementary Figure 12. Chicken chr1 is homologous to multiple gar chromosomes.** A circus plot is used to show the macrosynteny. Each chromosome is scaled to the number of genes encoded by this chromosome and is arranged in a way to better visualize the correspondence of chromosomes. Each line represents one pair of homologous genes between chromosomes in different species, which is color-coded according to chicken and gar chromosomes (green for chicken and grey for gar). Chicken chr1 corresponds to multiple gar chromosomes (e.g., LG12), which in turn correspond to additional chicken chromosomes (e.g., chr7). Such a complex relationship suggests chicken- or gar-specific chromosomal fusions or fissions. After polarization with the outgroup, i.e., elephant shark scaffold, we inferred that chicken chr1 has been subject to four fusions and thus manually split it back as chr1.1, 1.2, 1.3 and 1.4 to approximate gnathostome ancestral chromosomes (right panel). “*” marks the chromosome subject to back-mutation.

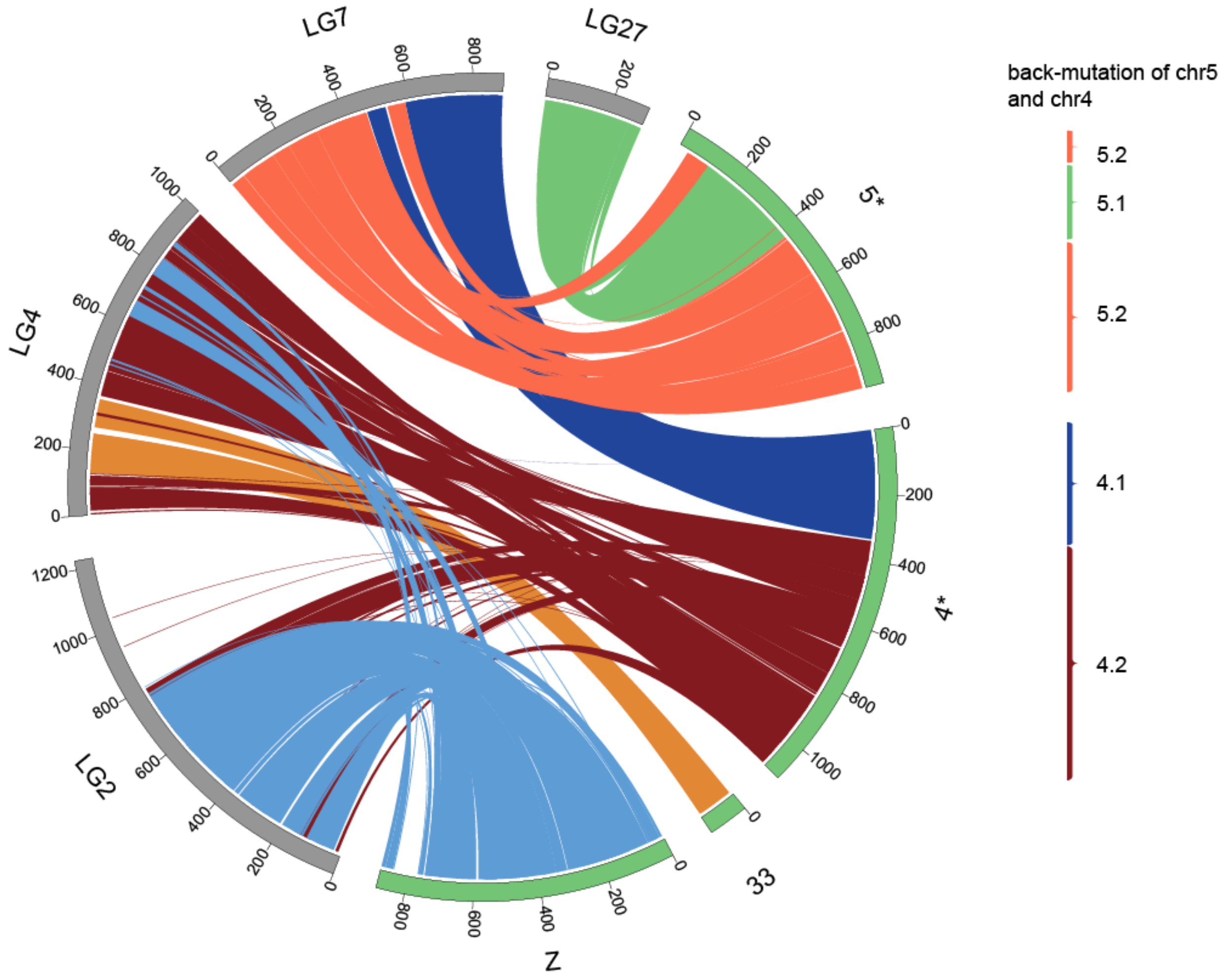


**Supplementary Figure 13. Both chicken chr4 and chr5 are homologous to two gar chromosomes, respectively.** The figure convention follows Supplementary Figure 12. Note that gar LG2 and LG4 are co-homologous with chicken chrZ and chr4.2.

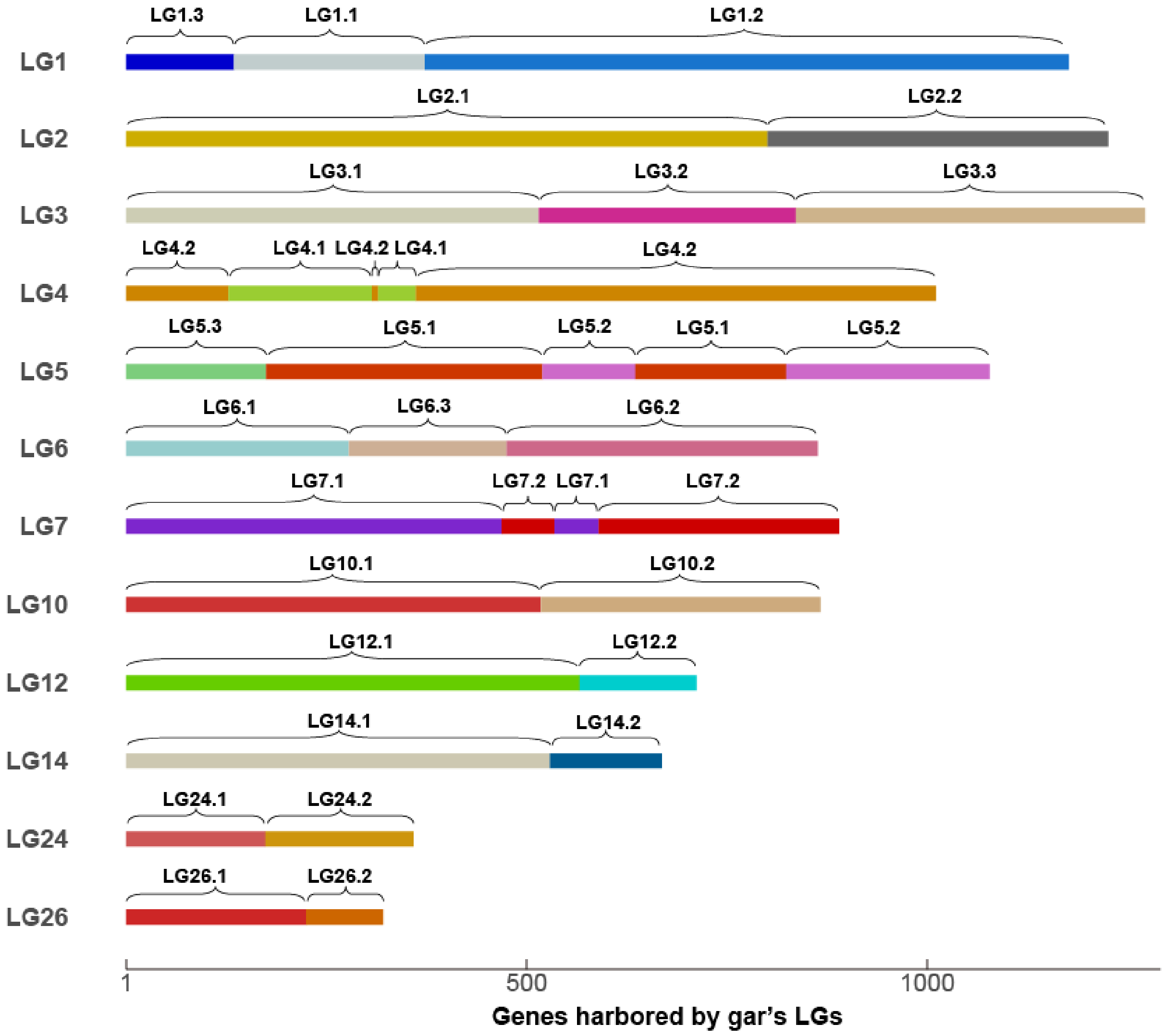


**Supplementary Figure 14. A schematic view of back-mutations in the gar genome.** Only gar LGs with lineage-specific fusions are shown. The length of LGs is proportional to the number of genes harboured by the corresponding LGs.

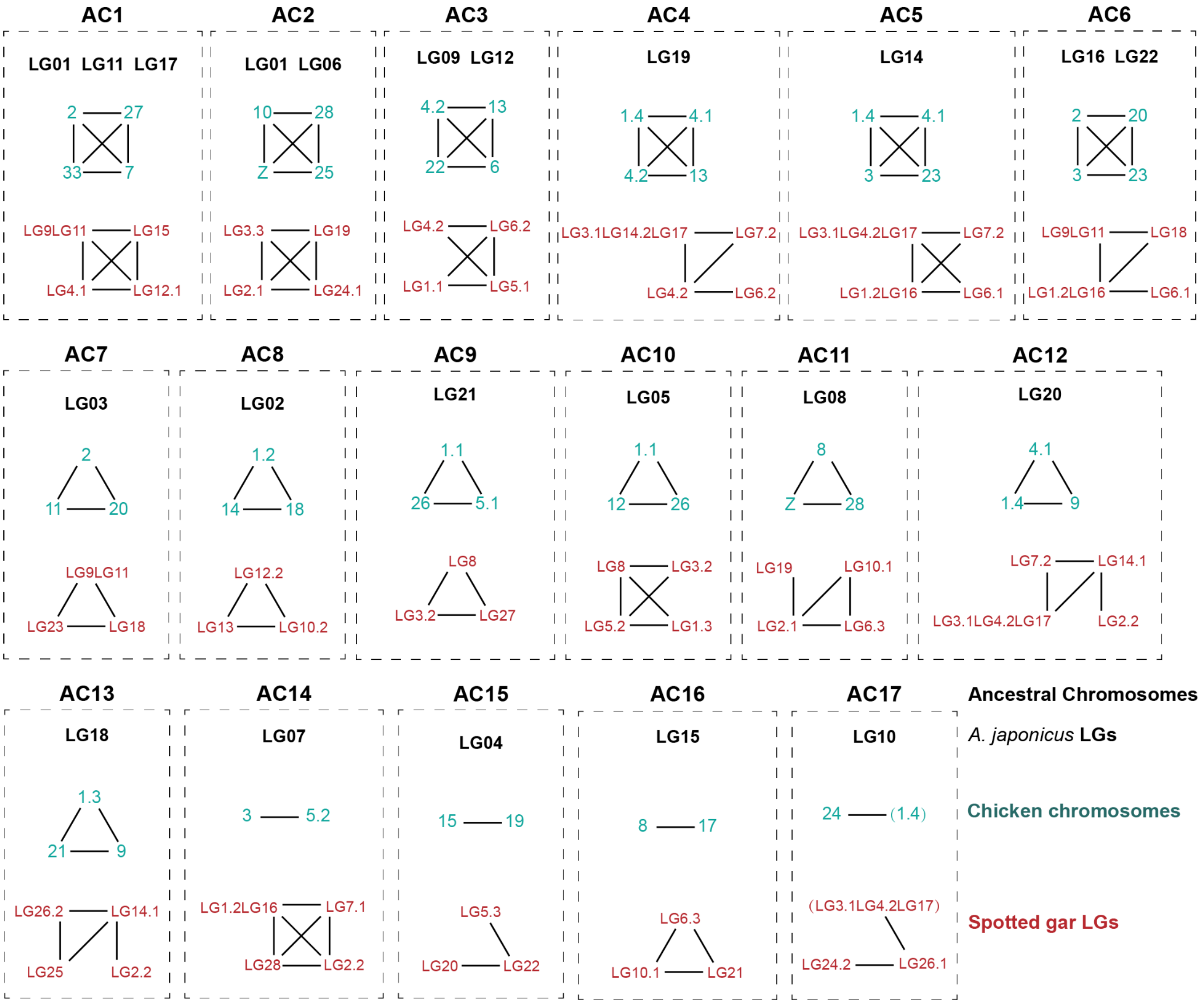


**Supplementary Figure 15. The homology between sea cucumber LGs and chromosomes of chicken and gar.** By examining whether a pair of chromosomes share an excess of homologous genes, we divided chicken and gar chromosomes homologous to the same sea cucumber LGs as 17 groups, where chromosomes connected with lines are paralogous between each other. For sea cucumber LG10, chicken chr1.4 and spotted gar LG3.1LG14.2LG17 are marked with parentheses because these two chromosomes are not significantly homologous to sea cucumber LG10. However, chicken chr24 is homologous to chr1.4 and gar LG26.1 is homologous to LG3.1LG14.2LG17, while chr24, LG24.2 and LG26.1 are homologous to sea cucumber LG10. So chr1.4 and LG3.1LG14.2LG17 appears to correspond to sea cucumber LG10 and the insignificance of correspondence could be due to the fact that chicken chr1.4 and gar LG3.1LG14.2LG17 already correspond to sea cucumber LG14, LG19 and LG20. Actually, after excluding chicken or gar genes mapping to sea cucumber LG14, LG19 and LG20, the relationship between chicken chr1.4 or gar LG3.1LG14.2LG17, and sea cucumber LG10 becomes significant or marginally significant ($q$ = 0.03, 0.06).

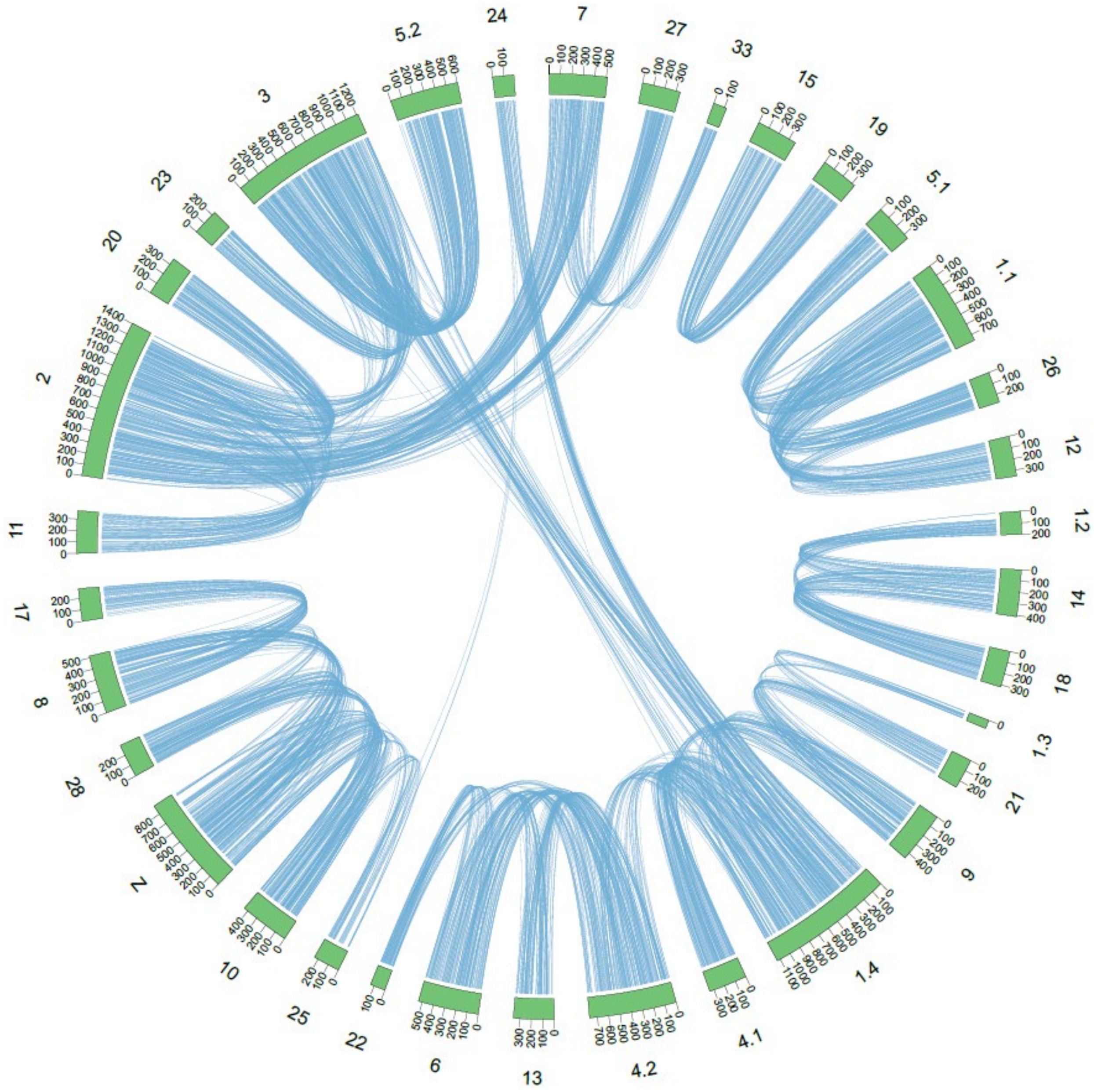


**Supplementary Figure 16. Extensive homology among chicken chromosomes.** To remove within-chromosome small scale duplications, only homologous gene pairs from homologous chromosome pairs are shown. This circus plot view supports the occurrence of at least one round of whole genome duplication.

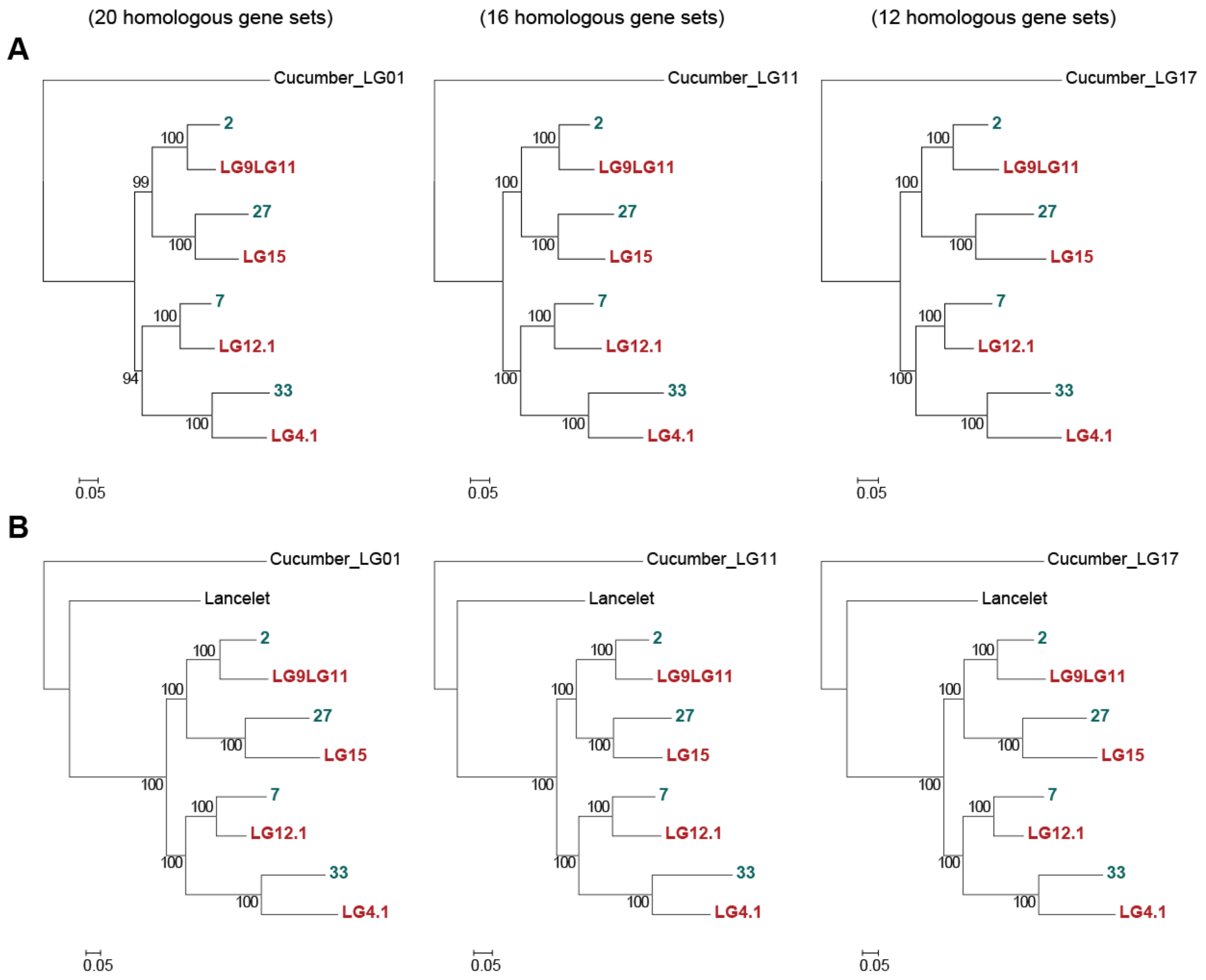


**Supplementary Figure 17. The phylogeny of chicken chromosomes chr2, 7, 27, and 33. a,** Chromosomal level trees with sea cucumber LG01, LG11 and LG17 as outgroups, respectively. Due to gene loss after WGD(s), only a small proportion of genes retains four duplicates in both chicken and gar. This is why we only have 12 to 20 homologous gene sets applicable for each chromosomal level tree reconstruction. **b**, Same as **a** except that lancelet homologous genes are added into each homologous gene set. Lancelet is not assembled at the chromosomal level and thus no chromosomal information is available. Cyan and red denote chicken chromosomes and gar LGs, respectively. Bootstrap scores are shown along the internal branches and the protein divergence is measured with the unit as 0.05 (5%). With sea cucumber LG01, LG11 or LG17 as outgroups, the same tree topologies were generated. Thus, the consensus tree of chicken chromosomes should be (((chr2, chr27), (chr7, chr33)), LG01/LG11/LG17).

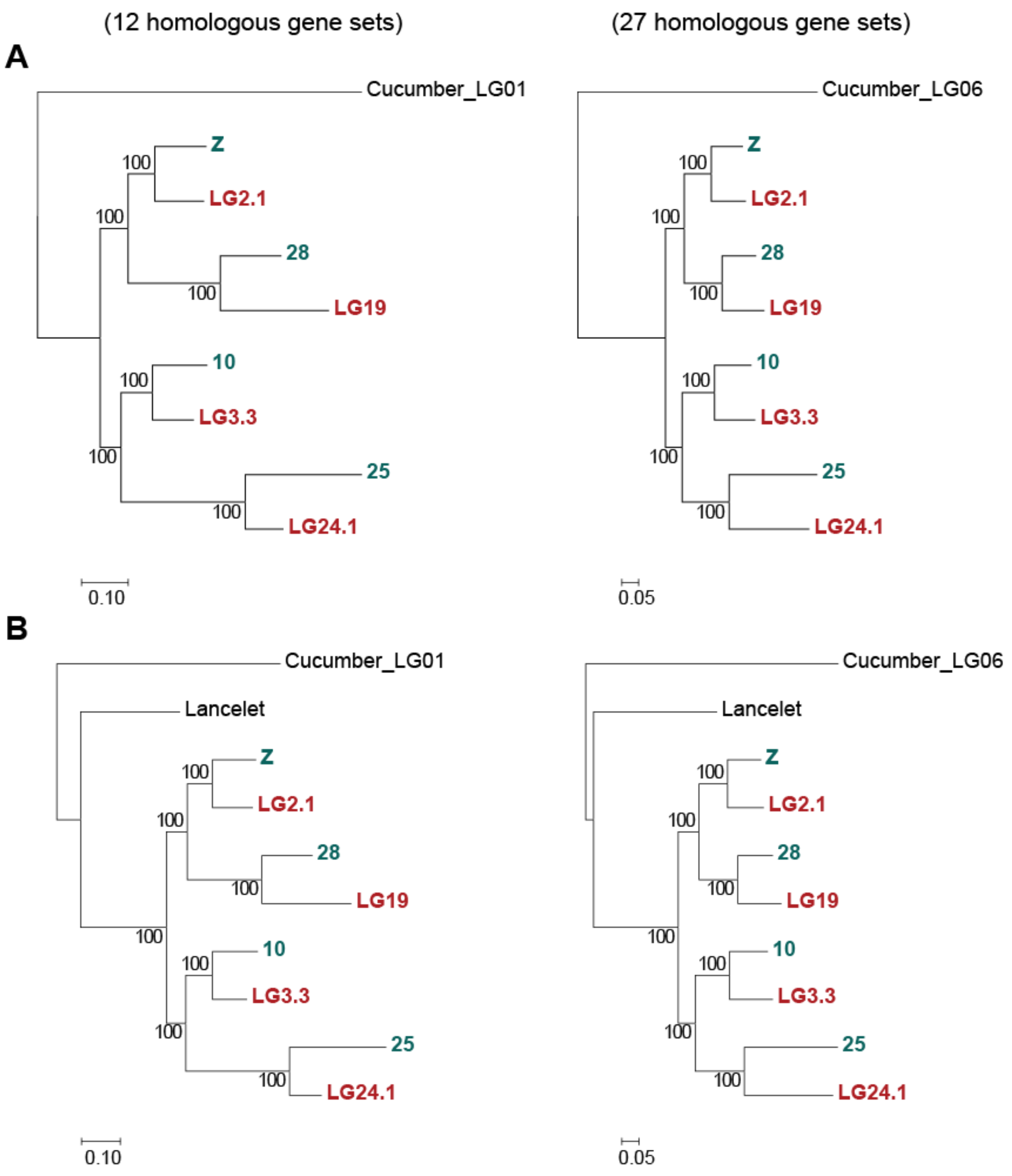


**Supplementary Figure 18. The phylogeny of chicken chromosomes chrZ, 28, 10, and 25.** This figure follows the same convention as Supplementary Figure 17. The consensus tree of chicken chromosomes appears to be (((chrZ, chr28), (chr10, chr25)), LG01/LG06).

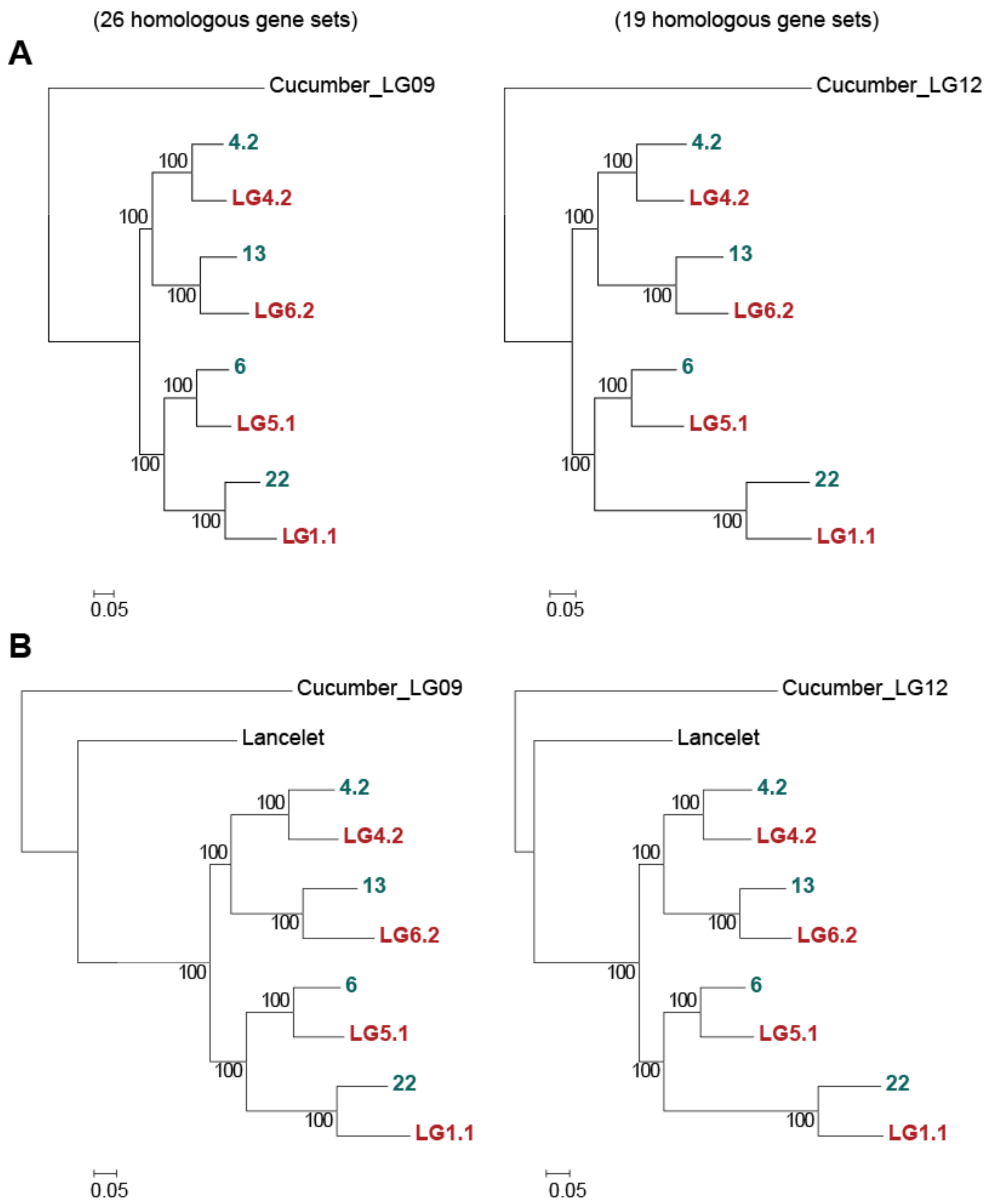


**Supplementary Figure 19. The phylogeny of chicken chromosomes chr4.2, 13, 6, and 22.** This figure follows the same convention as Supplementary Figure 17. The consensus tree of chicken chromosomes appears to be (((chr4.2, chr13), (chr6, chr22)), LG09/LG12).

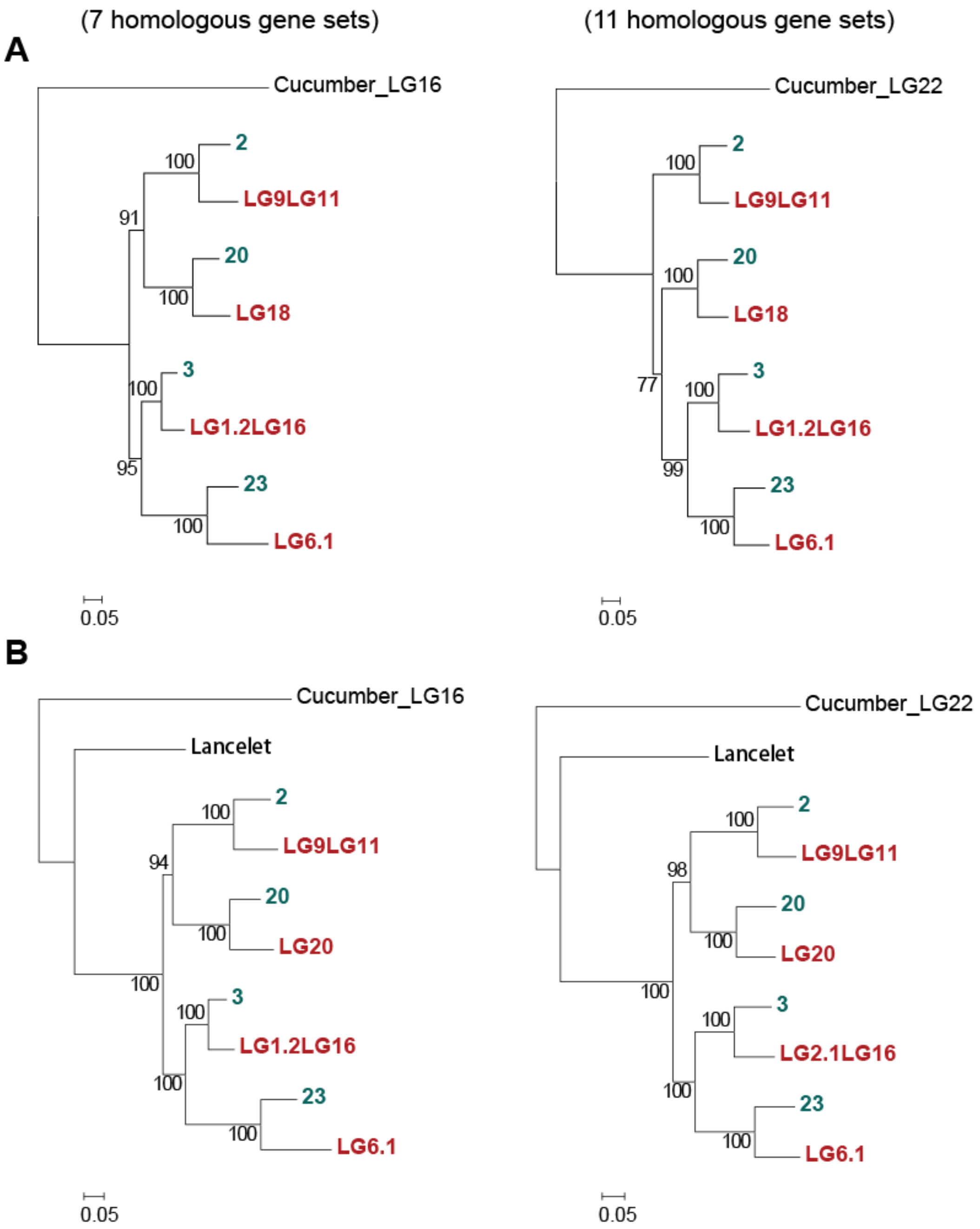


**Supplementary Figure 20. The phylogeny of chicken chromosomes chr2, 20, 3, and 23.** This figure follows the same convention as Supplementary Figure 17. The consensus tree of chicken chromosomes appears to be (((chr2, chr20), (chr3, chr23)), LG16/LG22).

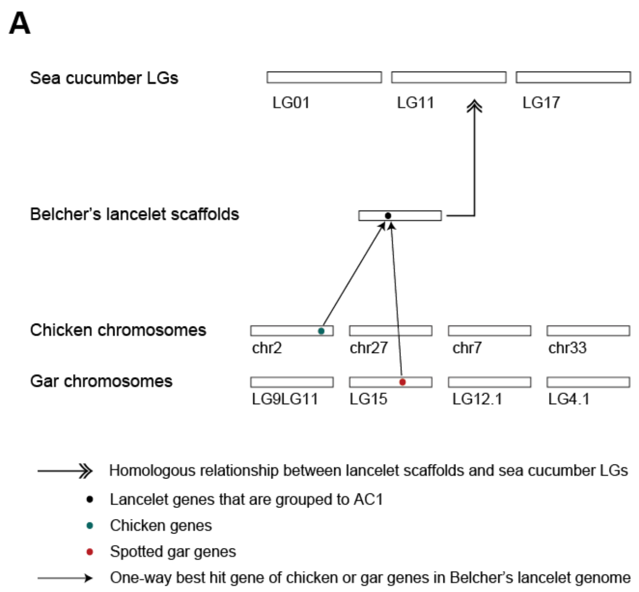


**Supplementary Figure 21. Two complementary strategies to reconstruct the gene content of ACs based on amphioxus genes.** We took AC1 as an example, which corresponds to sea cucumber LG01/LG11/LG17, chicken chr2, 7, 27, 33 and gar LG9LG11, LG15 LG12.1, LG4.1 (Supplementary table 9). **a,** For a lancelet scaffold that is homologous to sea cucumber LG01, LG11 or LG17, if at least one chicken gene from chr2, 7, 27, or 33 and at least one gene from gar LG9_11, LG15, LG12.1 and LG4.1 are co-homologous toward one gene encoded by this lancelet scaffold, we grouped the latter gene to AC1. **b,** If at least five genes from five different chromosome of chicken chr2, 7, 27, 33 and gar LG9_11, LG15, LG12.1, LG4.1 have one lancelet gene as their homolog, we grouped the later gene to AC1 without consideration of the homology of the corresponding lancelet scaffold and sea cucumber LGs. For the second strategy, the reason that we require five genes is to ensure that this small gene family should be generated by 2R WGDs in the ancestor of chicken and gar.

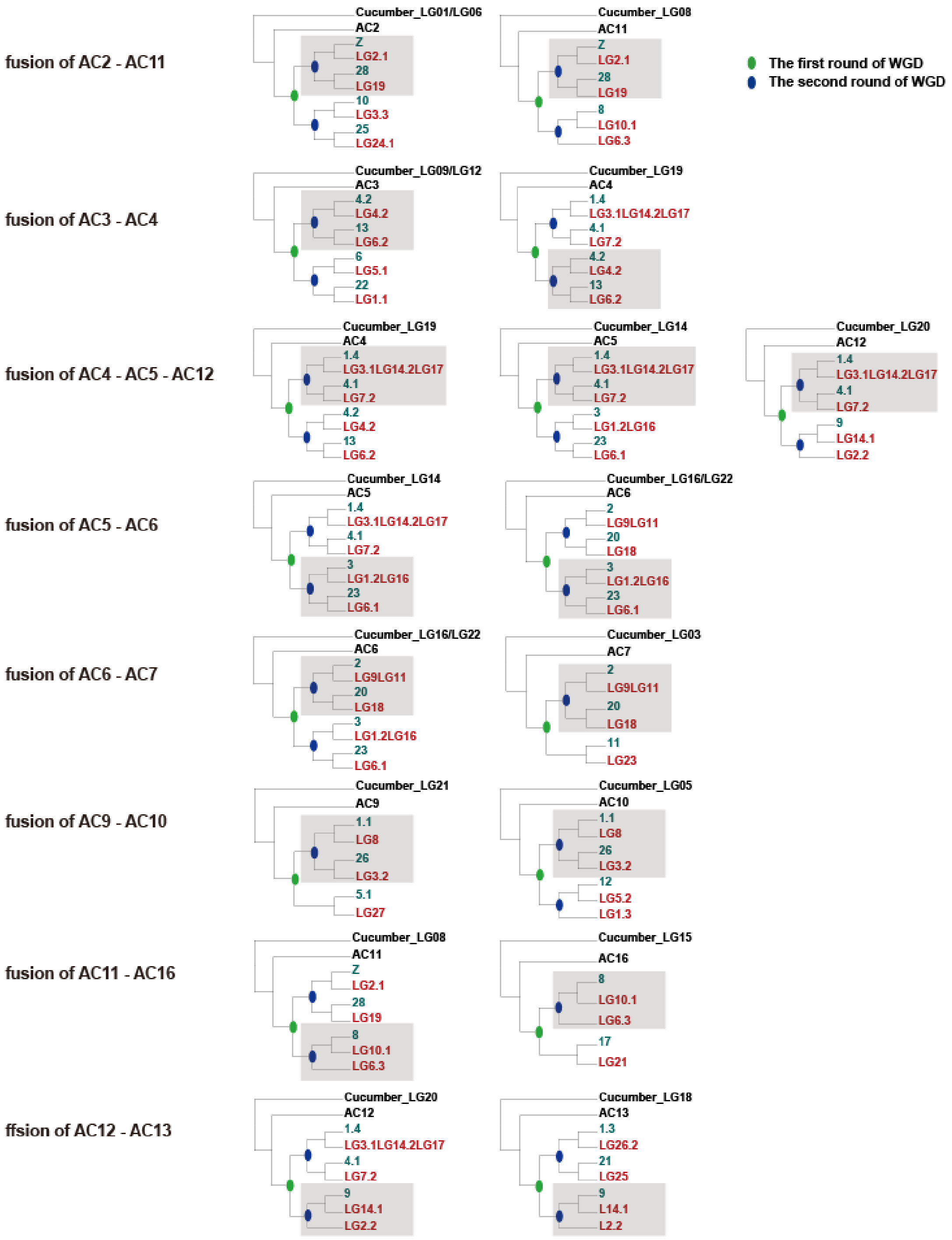


**Supplementary Figure 22. The eight post-1R fusions proposed based on chromosomal phylogenetic trees.** The products of the fused chromosomes after the second round of gnathostome WGD are marked in grey blocks.

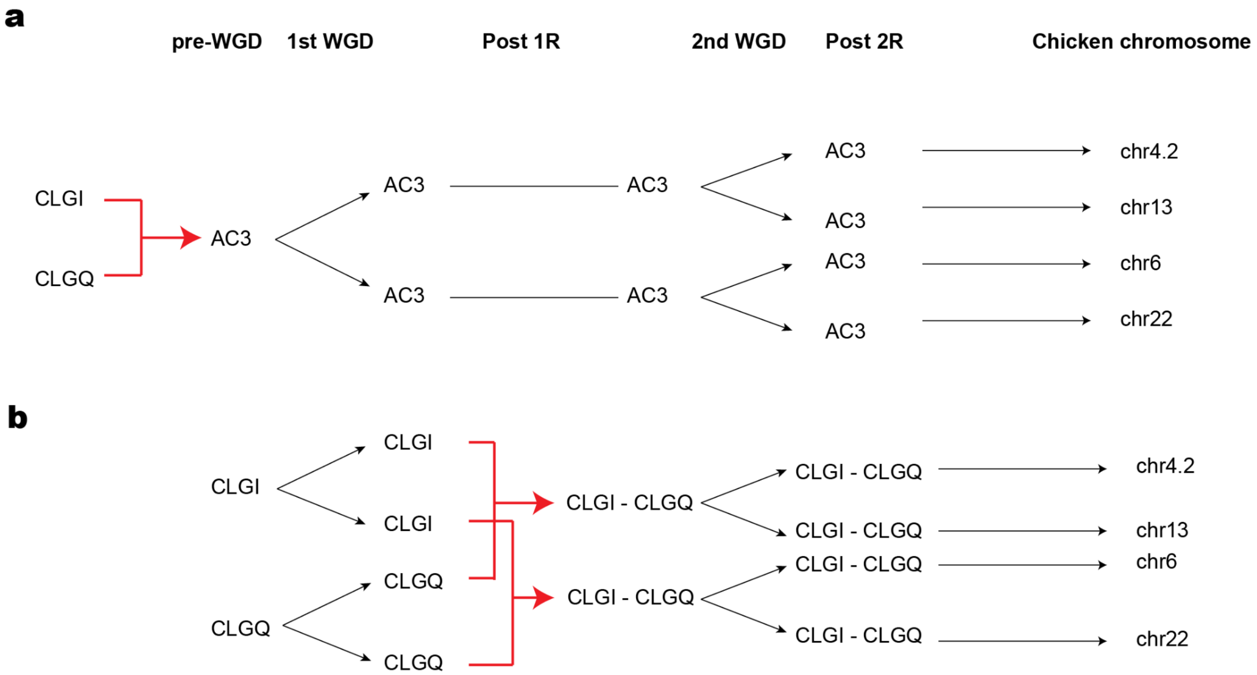


**Supplementary Figure 23. Two hypothetical hypotheses explaining the origin of chicken chr4.2, 13, 6 and 22. a,** A pre-1R fusion event is hypothesized in our 17 ACs model where AC3 is equivalent with a fusion product of CLGI and CLGQ in Simakov's 17 CLGs model[121]. **b,** Two pairwise post-1R fusion events were hypothesized in 17 CLGs model.

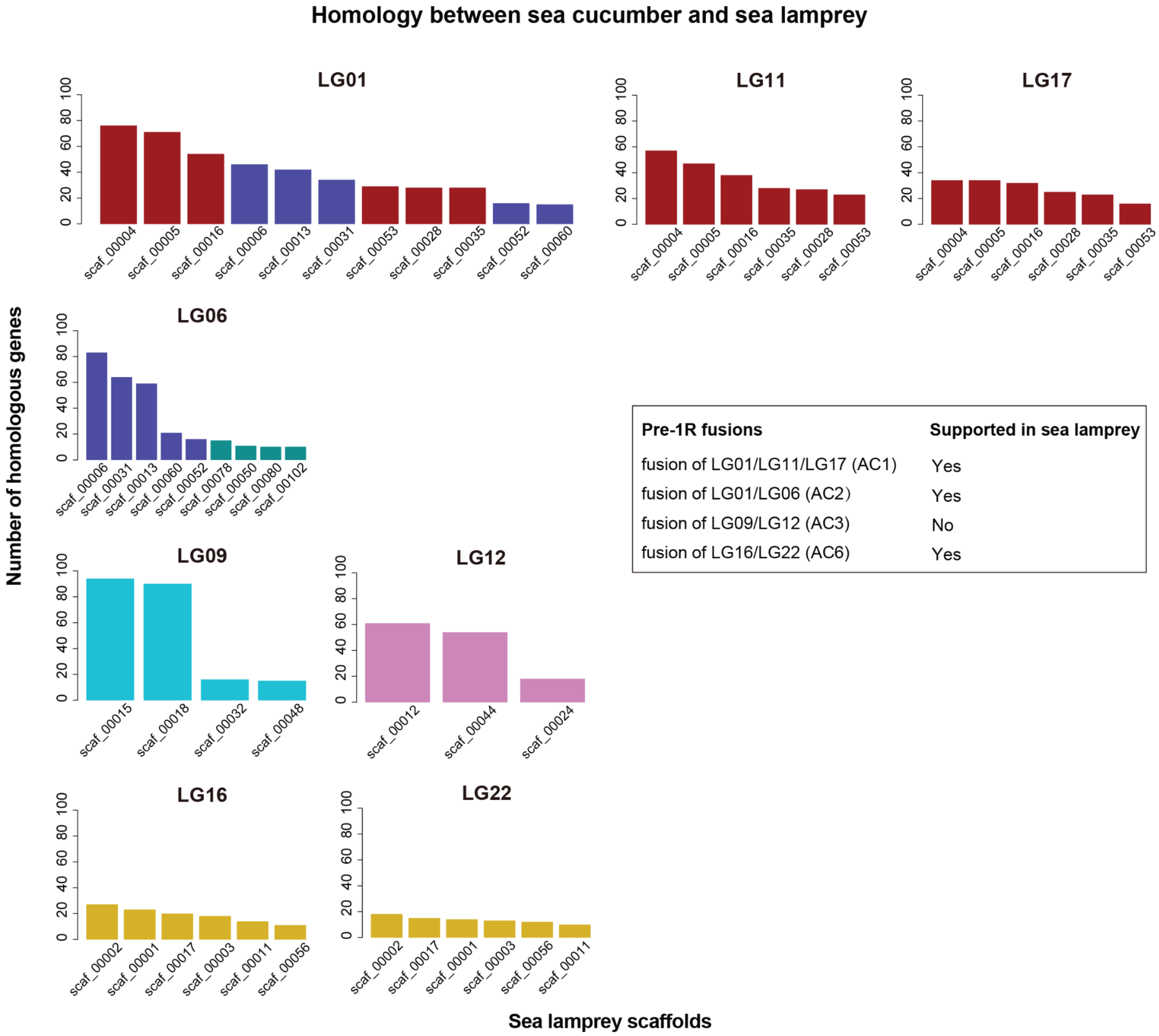


**Supplementary Figure 24. Three but one pre-1R fusions are shared by sea lamprey genome.** Four pre-1R fusions of ancestral chromosomal regions, corresponding to LG01/LG11/LG17, LG01/LG06, LG09/LG12, and LG16/LG22 of the sea cucumber genome, were proposed based on the analyses of sea cucumber, lancelet, chicken and gar genomes. Herein, sea lamprey scaffolds homologous to sea cucumber LG01, LG06, LG11, LG17, LG09, LG12, LG16 and LG12 are shown. They correspond to AC1 (LG01/LG11/LG17), AC2 (LG01/LG06), AC3 (LG09/LG12), and AC6 (LG16/LG22) respectively. Sea cucumber scaffolds supporting fusions (LG01/LG11/LG17, LG01/LG06 and LG16/LG22) are marked in the same color.

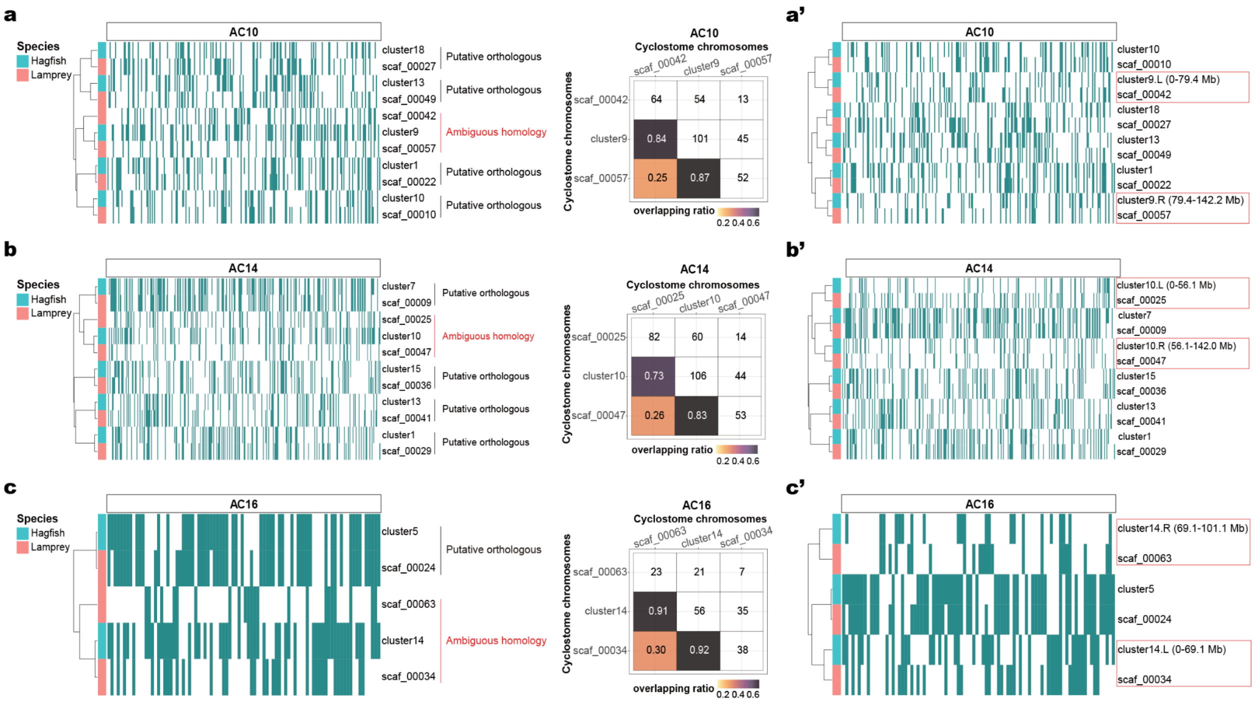


**Supplementary Figure 25. Three cases of hagfish-specific fusion of paralogous chromosomes were identified and back-mutated. a,** The left panel shows that hagfish Hi-C cluster 9 is most similar to sea lamprey scaffolds 42 and 57 in terms of AC10 derived homologs, while the right panel shows that the two lamprey scaffolds are likely paralogous suggested by their decent overlapping ratio. **a',** After manually splitting of hagfish Hi-C cluster 9 into two parts (split at 79.4 Mb) into cluster9.L and cluster9.R, these cluster well separately with sea lamprey scaffolds 42 and 57, respectively, indicating that hagfish cluster9 is the result of a fusion of two paralogous chromosomes. **b, b', c** and **c'** show two further examples with chromosomes deriving from AC10 and AC16. Another example is that of *Hox*-containing hagfish Hi-C cluster 3 (see Extended Data Figure 4e).

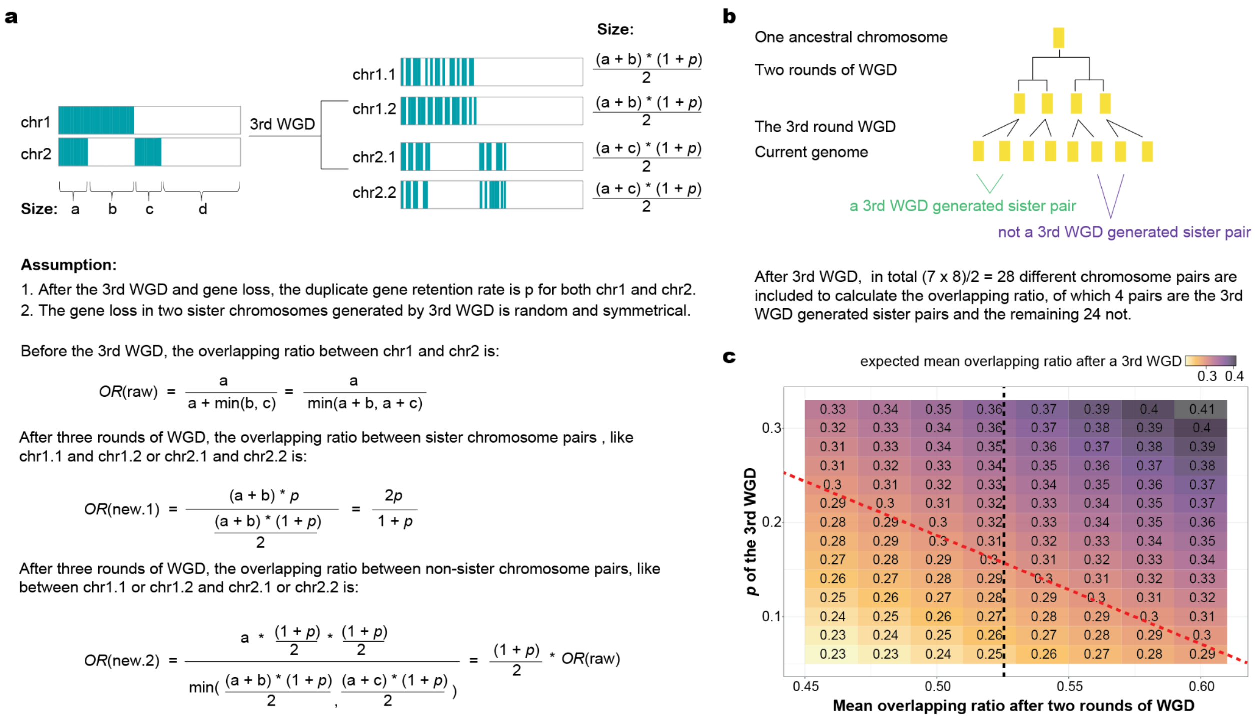


**Supplementary Figure 26. More rounds of WGDs cause the continuous decrease of overlapping ratio. a,** With one more round of WGD, the overlapping ratio drops with a rate (1+p)/2, where p is equal to the percentage of retained duplicates. **b,** A schema on how a third round of WGD changes the number of sister chromosomes and thus the calculation of average overlapping ratio between paralogous chromosomes. **c,** The mean overlapping ratio after the third round of WGD is calculated based on a different combination of parameters including the duplicate retention rate (p) and mean overlapping ratio after two rounds of WGDs. A black dashed line indicates the observed median overlapping ratio (0.52) of paralogous chromosomes in gnathostomes shaped by two rounds of WGDs, while a red dashed line indicates the observed median overlapping ratio (0.30) in cyclostomes.

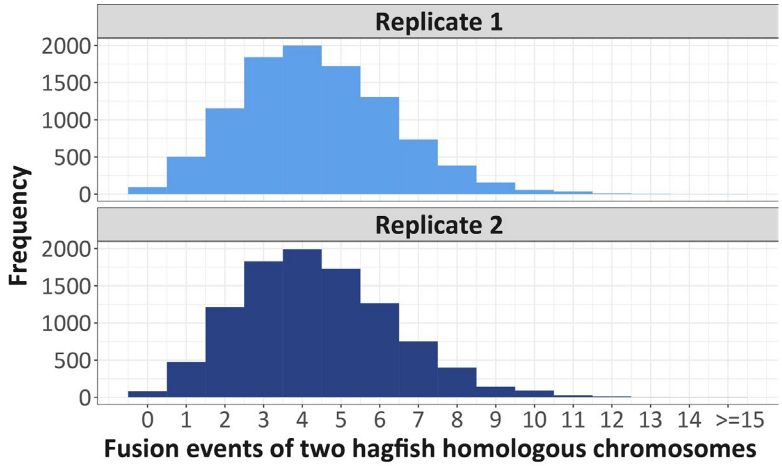


**Supplementary Figure 27.** Frequency distribution of fusion events of two hagfish paralogous chromosomes. We randomly fused hagfish chromosomes and counted how many times that paralogous chromosomes were fused.

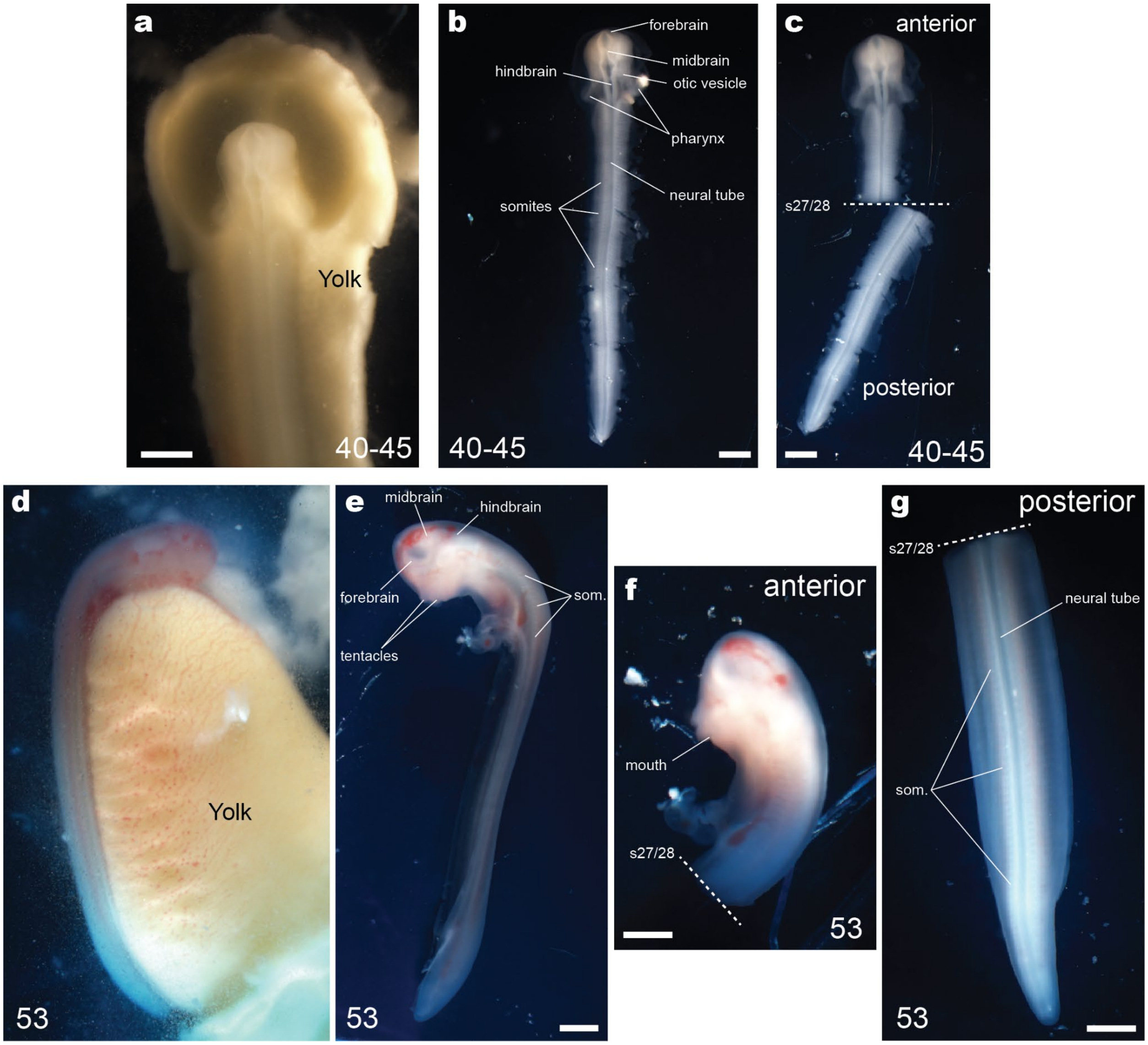


**Supplementary Figure 28. Hagfish embryos used for ATAC-seq. a-c,** Embryo of the inshore hagfish *E. burgeri* at stage Dean ~40-45 extracted from the eggshell. Yolk was removed almost completely in (**a**), and completely in (**b**,**c**). Key anatomical features are labeled. The embryo was divided into anterior and posterior halves (**c**) at the level between somites 27 and 28 (dashed line) and each was assayed separately. **d-g**, Embryo at stage Dean 53. As with before, we divided the embryo into anterior (**f**) and posterior (**g**) halves at an equivalent A-P level (somites 27 and 28, dashed lines). Staging was performed according to a previous study[151]. Scale bars, 1 mm.

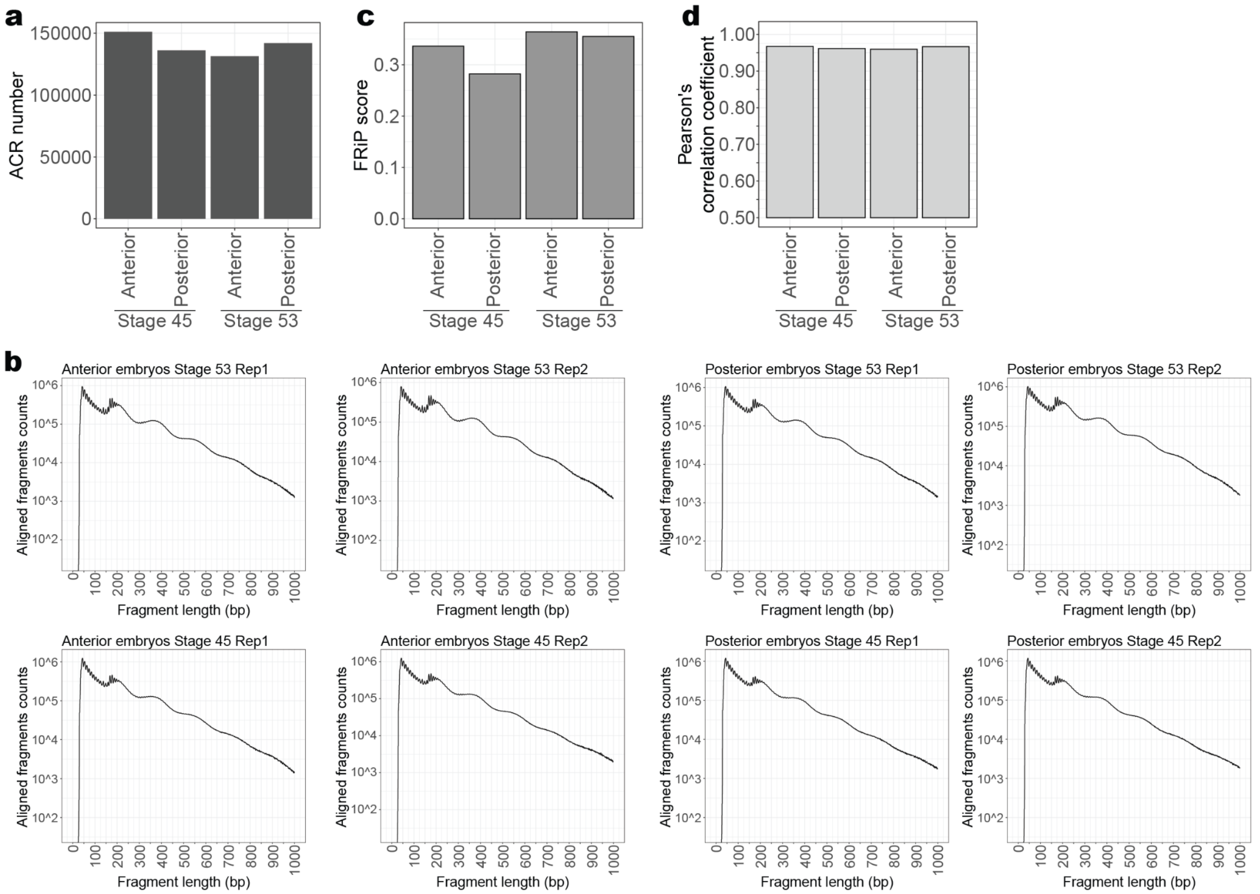


**Supplementary Figure 29. Validation of ATAC-seq data of hagfish embryos. a,** Number of identified ACRs in each sample. **b,** Distributions of fragment size of properly-paired aligned reads in each replicate data. **c,** A fraction of all aligned reads in the called ACRs (FRiP score, which is a genome-wide, quality metric of the signal-to-background ratio[190]). **d,** The Pearson correlation coefficients of ATAC-seq signal intensities (log10-RPM; reads per million mapped reads) of ACRs between the technical replicates at different samples.

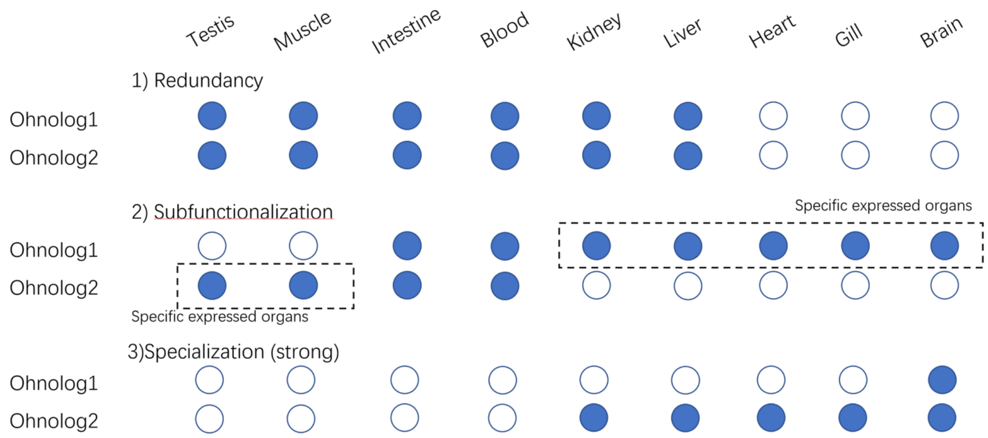


**Supplementary Figure 30. Possible fates of ohonologs after WGD according to their expressional domains.** Ohnologous genes with the same expressional tissues are classified as redundant (1); Ohnologs with each a tissue-specific expression pattern are considered as subfunctionalized (2); and ohnologous families in which one of the ohnologs is expressed in a subset of the expressional domains of the other corresponding ohnolog is considered specialized (3). Shown in the scheme is a case of ‘strong specialization’, in which the ohnolog with the narrower pattern is expressed in < 40% of domains than the ohnolog with the larger pattern.

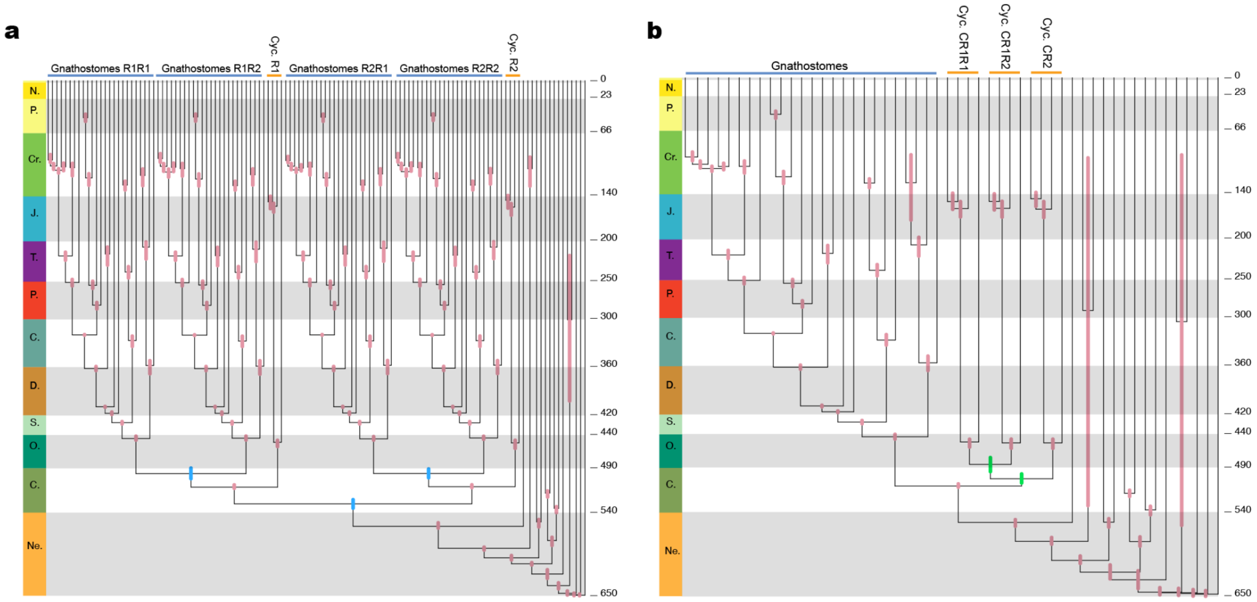


**Supplementary Figure 31. Dating Whole Genome Duplication events.** **a**, Dating using a multigene alignment focussed on the 1R and 2R events. **b**, Dating using a multigene alignment focussed on the CR1 and CR2 events.
